# Inhibiting the GPI Transamidase Subunit GPAA1 Abolishes CD24 Surface Localization and Enhances Macrophage-Mediated Phagocytosis of Ovarian Cancer Cells

**DOI:** 10.1101/2023.10.21.563315

**Authors:** Alok K. Mishra, Tianyi Ye, Shahid Banday, Ritesh P. Thakare, Chinh Tran-To Su, Ngoc N. H. Pham, Amjad Ali, Ankur Kulshreshtha, Shreya Roy Chowdhury, Tessa M. Simone, Kai Hu, Lihua Julie Zhu, Birgit Eisenhaber, Sara K. Deibler, Karl Simin, Paul R. Thompson, Frank Eisenhaber, Sunil K. Malonia, Michael R. Green

## Abstract

The CD24-Siglec10 signaling axis is an immune checkpoint pathway that shields ovarian cancer cells from phagocytosis by tumor-associated macrophages (TAMs), making it an appealing immunotherapeutic target. Here, we investigate factors influencing CD24 cell surface expression and assess their suitability as drug targets. Using a CRISPR-based knockout screen, we identify GPAA1 (glycosylphosphatidylinositol anchor attachment-1) as a positive regulator of CD24 cell surface expression. GPAA1 is a crucial component of the multi-subunit GPI transamidase complex, which facilitates the attachment of GPI lipid anchor to the C-terminus of CD24, enabling its surface localization. Reducing the activity of GPAA1 in ovarian cancer cells, either by genetic ablation or targeting with an aminopeptidase inhibitor bestatin, disrupts GPI attachment to CD24. This disruption impairs CD24 cell surface localization, enhances phagocytosis by TAMs, and suppresses tumor growth in mice. Our study highlights the potential of GPAA1 targeting as a therapeutic approach for CD24-positive ovarian cancers.

## INTRODUCTION

Ovarian cancer is the second most common gynecologic cancer and the leading cause of gynecologic malignancy-related deaths in the United States.^1^ The standard treatment for newly diagnosed ovarian cancer includes cytoreductive (debulking) surgery followed by chemotherapy.^2^ However, recurrence affects 25% of early-stage patients and 70-80% of advanced-stage patients,^3,4^ highlighting the urgent need to explore alternative therapeutic strategies.

Immuno-oncology strategies have demonstrated remarkable success in stimulating immune responses against diverse tumor types and have significantly improved patient outcomes. For example, conventional immune checkpoint inhibitors targeting the PDL1/PD1 pathway have shown efficacious responses in several solid cancers, such as malignant melanoma, non-small-cell lung cancer, and urothelial cancer.^5^ However, in ovarian cancers, PDL1/PD1 inhibitors have shown limited efficacy.^6–8^

Recent studies have identified the CD24-Siglec10 immune checkpoint as a promising therapeutic target in ovarian cancer.^9^ CD24 is a highly glycosylated cell adhesion protein that is linked to the plasma membrane by a glycosylphosphatidylinositol (GPI) anchor.^9^ CD24 is primarily expressed by immune cells but is often highly expressed in many cancers, most notably ovarian cancer.^10,11^ CD24 facilitates immune evasion by interacting with its receptor, Siglec10, a transmembrane protein present on the surface of tumor-associated macrophages (TAMs) and providing an anti-phagocytosis (“don’t eat me”) signal that hinders TAMs from targeting and engulfing tumor cells.^10^ Previous studies have shown that CRISPR-mediated knockout of CD24 or its blockade with an anti-CD24 monoclonal antibody enhances TAM-mediated phagocytosis of ovarian cancer cells and inhibits tumor growth in mouse models.^10,12^

In this study, we aimed to identify factors and pathways that could be pharmacologically inhibited to reduce CD24 expression and its function; such factors could represent novel immunotherapeutic targets for ovarian cancer. As a first step, we performed a genome-wide CRISPR/Cas9 screen to identify factors that promote CD24 cell surface expression. Using this approach, we identified GPAA1 (GPI anchor attachment 1), a critical component of a multi-subunit complex that facilitates attachment of a GPI anchor to substrate proteins, thereby directing them to the cell surface membrane.^13^ Interestingly, GPAA1 shares structural similarities with a specific class of aminopeptidases^14^ and we found that an aminopeptidase inhibitor bestatin binds to GPAA1, inhibiting its function. Genetic or pharmacological inhibition of GPAA1 in ovarian cancer cells decreased CD24 cell surface expression, increased phagocytosis by TAMs, and reduced ovarian tumor growth in mice xenografts. Our findings open a new avenue for developing small-molecule immunotherapeutics for CD24-positive ovarian cancers.

## RESULTS

### A genome-scale CRISPR-Cas9 knockout screen identifies factors that promote CD24 cell surface expression in ovarian cancer

To identify factors and pathways regulating CD24 cell surface expression in ovarian cancer, we performed a genome-wide CRISPR/Cas9 screen. To select an appropriate cell line for the screen, we performed CD24 flow cytometry analysis in a panel of human ovarian cancer cell lines. Our analysis revealed robust CD24 cell surface expression among seven out of eight ovarian cancer cell lines tested (A1847, IGROV1, NCI/ADR-RES, OVCAR3, OVCAR4, OVCAR8, and SKOV3), all of which exhibited greater than ∼85% CD24-positive cells (Figure S1A). The exception was the A2780 cell line, which was CD24 negative. We selected the OVCAR8 cell line for the screen because it had the highest percentage of CD24-positive cells, and it is a well-characterized high-grade serous ovarian cancer (HGSOC) cell line^15^. To ensure the suitability of OVCAR8 cells for the CRISPR/Cas9 knockout screen, we confirmed that the proliferation of OVCAR8 cells was unaffected by the loss of CD24 (Figures S1B and S1C).

For the screen, we established OVCAR8 cell line that stably expressed active Cas9 (Figures S1D and S1E). These cells were transduced with the Human Brunello CRISPR knockout pooled library, which consists of ∼76,000 sgRNAs targeting ∼19,000 genes (4 sgRNAs per gene).^16^ After 15 days of puromycin selection, cells were stained with an anti-CD24 antibody, and those with substantially reduced cell surface expression of CD24 (CD24^low^ cells) were isolated by fluorescence-activated cell sorting (FACS). We identified sgRNAs that were significantly enriched in the CD24^low^ population relative to the unsorted population by bioinformatic analysis of deep sequencing data (Figure 1A).

**Figure 1.**
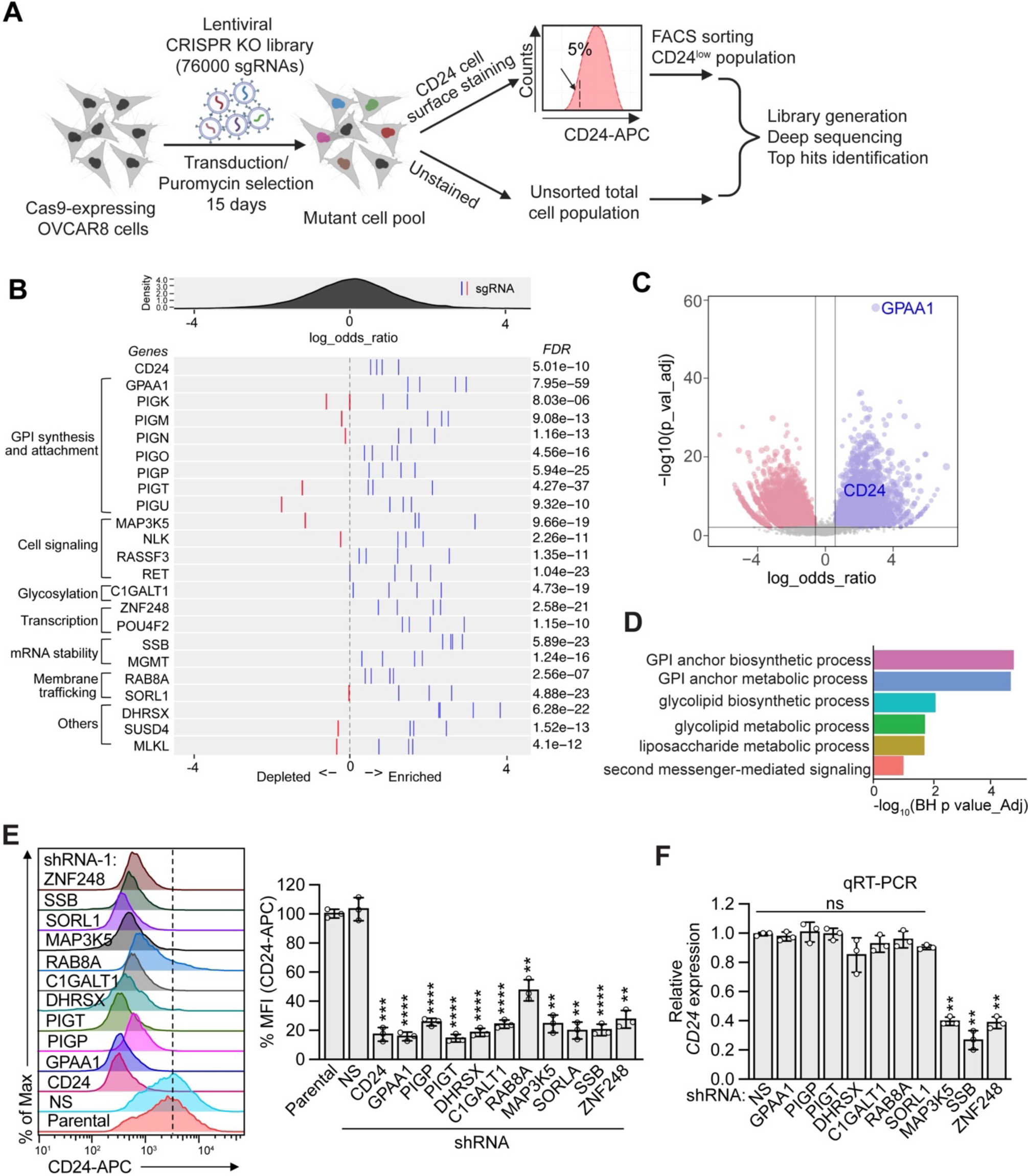
A genome-scale CRISPR/Cas9 screen identifies factors that promote cell surface expression of CD24 in ovarian cancer. (A) Schematic of the pooled CRISPR/Cas9 screen, created with BioRender.com. (B) FDR plot showing the distribution of sgRNAs targeting selected genes enriched (blue lines) or depleted (red lines) in CD24^low^ cells. (C) Volcano plot displaying significantly enriched (blue dots) or depleted (red dots) candidate genes identified from the screen. The top candidates GPAA1 and CD24 are shown. (D) GO analysis showing the most significant biological processes related to the enriched sgRNAs in CD24^low^ cells. (E) (Left) Representative flow cytometry histograms of CD24 cell surface expression in OVCAR8 cells expressing a shRNA targeting each candidate factor, or as a control a non-silencing (NS) shRNA. (Right) Quantification of mean fluorescence intensity (MFI). The results were normalized to that obtained in parental OVCAR8 cells. (F) QRT-PCR analysis monitoring *CD24* expression in OVCAR8 cells expressing an NS shRNA or shRNA targeting each candidate factor. The results were normalized to that obtained with an NS shRNA. Data represents ± SD (n=3). P values were calculated using one-way ANOVA followed by Dunnett’s multiple comparisons test. *\*\*P<0.01, ***P<0.001, ****P<0.0001.* See also Figure S1

Our screen yielded candidate genes representing a broad range of functional categories including intracellular signalling, transcriptional regulation, mRNA stability, post-translational modification, and membrane trafficking (Figure 1B). As anticipated, all four sgRNAs targeting CD24 exhibited statistically significant enrichment (Figure 1B). Notably, the most highly enriched candidate in the screen was GPAA1 (Figures 1B and 1C), a component of the multi-subunit GPI transamidase (GPIT) complex that mediates attachment of a GPI anchor to substrate proteins, which serves as a mechanism for directing GPI associated proteins (GPI-APs) to the cell membrane.^17^ Proteins that are destined to become GPI anchored harbor an N-terminal endoplasmic reticulum (ER) localization sequence and a C-terminal GPI-attachment signal peptide, and are translocated to the ER where the attachment of a pre-synthesized GPI lipid anchor occurs.^13,18^ Once the GPI anchor is attached, the protein is shuttled to the Golgi, where it undergoes fatty acid remodeling before being transported to the outer leaflet of the cell membrane. The attachment of GPI to substrate proteins is crucial for their proper localization and function.^17^ In addition to GPAA1, the primary screen identified other components of the GPIT complex, including PIGK, PIGT and PIGU (phosphatidylinositol glycan anchor biosynthesis class K, T and U, respectively),^19^ as well as other proteins involved in GPI anchor biosynthesis, including PIGM, PIGN, PIGO and PIGP^13,20^ (Figure 1B). Gene ontology enrichment analysis of the significant hits underscored GPI anchor biosynthesis and metabolism as the most significant biological processes (Figure 1D). We selected GPAA1 and nine additional genes representing various functional categories for validation. We knocked down each gene using two independent shRNAs and monitored CD24 cell surface expression by flow cytometry. As shown in figure 1E, stable knockdown of each of the 10 candidates significantly reduced CD24 cell surface expression when compared to a control non-silencing (NS) shRNA (see also Figures S1F and S1G). To determine whether knockdown of these candidates reduced CD24 expression at the transcriptional or post-transcriptional level, we assessed *CD24* mRNA levels by quantitative RT-PCR (qRT-PCR). Knockdown of three of the ten candidates (MAP3K5, SSB and ZNF248) significantly reduced *CD24* mRNA levels, suggesting that they promote CD24 expression at the transcriptional level (Figures 1F and S1H). Conversely, knockdown of the other seven candidates (GPAA1, PIGP, PIGT, DHRSX, C1GALT1, RAB8A and SORL1) did not alter *CD24* mRNA levels, suggesting their potential roles in either post-transcriptional or post-translational regulation of CD24.

### GPAA1 promotes CD24-mediated inhibition of cellular phagocytosis

We focused our further investigation on GPAA1 for several reasons. First, as mentioned above, GPAA1 emerged as the top-scoring factor from the primary screen (Figures 1B and 1C). Second, previous studies have shown that GPAA1 is overexpressed in several types of cancers including ovarian cancer.^21–24^ Third, an examination of the cBio Cancer Genomics Portal^25^ revealed that GPAA1 is amplified in approximately 30% of ovarian cancers (Figure S2A). In addition, analysis of publicly available single-cell RNA sequencing (scRNA-seq) data from ovarian tumors indicated high expression of GPAA1 in the ovarian epithelial cancer cell cluster compared to other cell types in the tumor microenvironment (Figure 2A). Interestingly GPAA1 was found to be co-expressed with CD24 in the ovarian cancer cell cluster (Figure 2B). Furthermore, Kaplan-Meier survival analysis of publicly available gene expression datasets from HGSOC patients^26^ indicated that high expression of GPAA1 correlates with a reduced probability of overall survival (Figure S2B). Finally, as elaborated more below, GPAA1 possesses a domain with primary sequence and predicted structure similar to metallo-aminopeptidases,^14,27^ suggesting it could potentially be inhibited by small-molecule aminopeptidase inhibitors, several of which are in advanced stages of clinical development.^28^ Collectively, these observations strongly suggest that GPAA1 represents a targetable component of the GPI synthesis pathway, and its inhibition could represent an immunotherapeutic approach for ovarian cancer.

**Figure 2.**
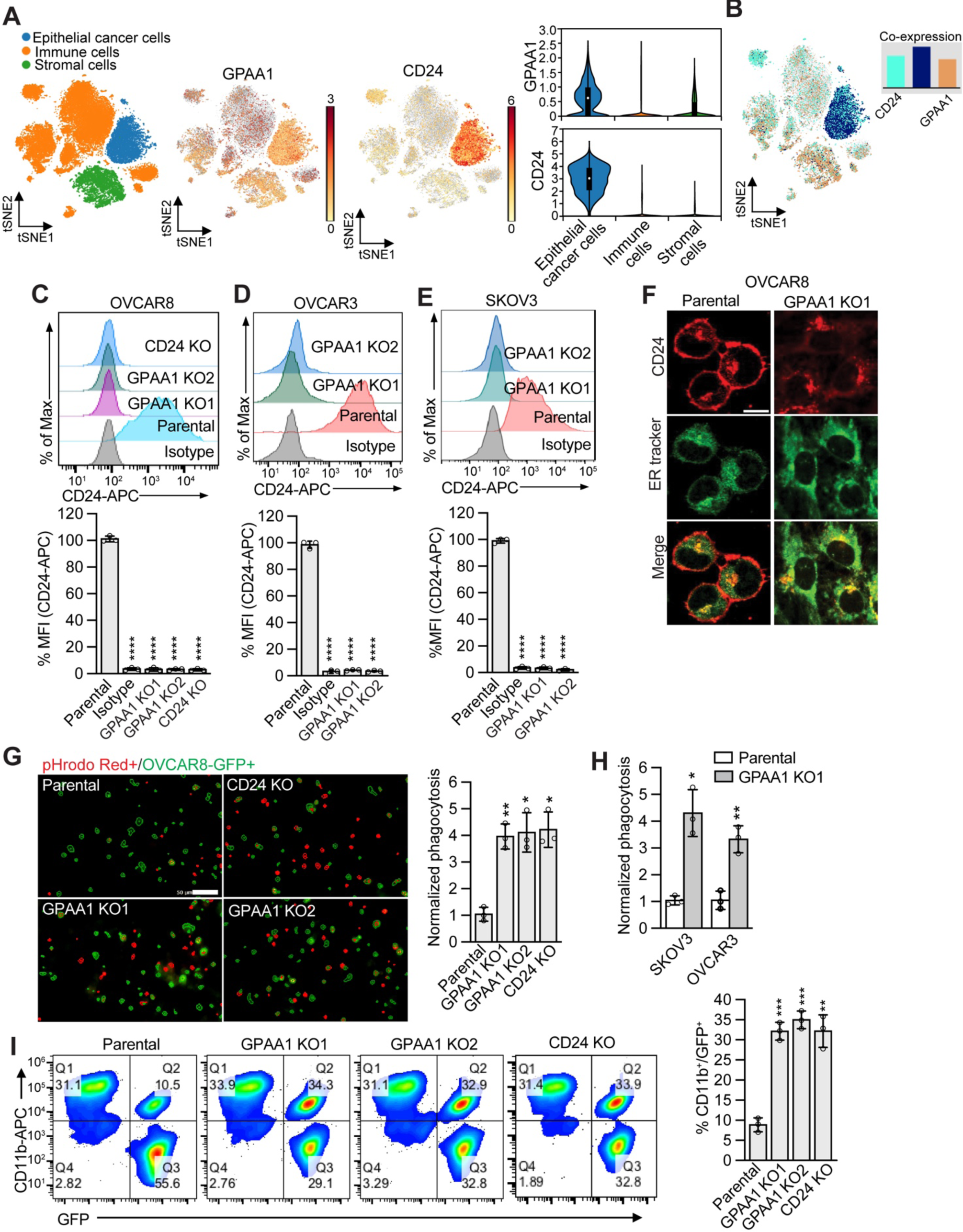
GPAA1 inhibition promotes ovarian cancer cell phagocytosis. (A) t-SNE and violin plots generated from the scRNA-seq dataset (GSE165897) showing expression of *GPAA1* and *CD24* in different cell types in high-grade serous ovarian cancer (HGSOC) patient samples. (B) t-SNE plot of scRNA-seq data showing co-expression of *GPAA1* and *CD24* in HGSOC patient samples. (C-E) (Top) Representative flow cytometry histograms of CD24 cell surface expression in parental, two independently derived single-cell GPAA1 knockout clones (KO1 and KO2) in OVCAR8 (C), OVCAR3 (D) and SKOV3 (E) cells, and in a CD24 knockout OVCAR8 clone. (Bottom) Quantification of MFI. (F) Representative confocal microscopy images of parental OVCAR8 or OVCAR8 GPAA1 KO1 cells stained with an anti-CD24 antibody or ER tracker. Merged images are shown. Scale bar, 10 µm. (G) *In vitro* phagocytosis assay (Left) Representative microscopy images of pHrodo Red^+^ GFP^+^ parental, GPAA1 KO, and CD24 KO OVCAR8 cells. Scale bar, 50 µm. (Right) Plot showing phagocytic events (pHrodo Red^+^ GFP^+^ cells), which was normalized to that obtained in parental OVCAR8 cells. (H) Plot showing normalized phagocytic events (pHrodo Red^+^ GFP^+^cells) in parental, GPAA1 KO OVCAR3 and SKOV3 cells. (I) (Left) Representative flow cytometry plots depicting the macrophage-mediated phagocytosis of parental, GPAA1 KO, and CD24 KO OVCAR8 cells. (Right) Quantification of phagocytic events showing double-positive macrophages (GFP+ and CD11b+). Data represents ± SD (n=3). P values were calculated using one-way ANOVA followed by Dunnett’s multiple comparisons test. *\*P<0.05, **P<0.01, ***P<0.001, ****P<0.001*. See also Figure S2.

To delve deeper into the role of GPAA1 in regulating CD24 localization, we employed CRISPR/Cas9-mediated genome editing to generate two independent GPAA1 knockout clones (GPAA1 KO1 and KO2) in OVCAR8, OVCAR3 and SKOV3 cell lines (Figures S2C-E). As a control, we also generated a CD24 knockout OVCAR8 cell line (Figure S2C). Remarkably, GPAA1 knockout in ovarian cancer cells did not affect cell proliferation (Figure S2F). As anticipated, GPAA1 knockout in ovarian cancer cells completely abrogated CD24 cell surface expression (Figures 2C-E), without affecting *CD24* mRNA levels (Figure S2G). The loss of CD24 cell surface expression in GPAA1 knockout OVCAR8 cells was also confirmed by immunocytochemistry, which revealed an accompanying accumulation of CD24 in the ER (Figure 2F).

As previously mentioned, CD24 protects cancer cells from phagocytosis by Siglec10-expressing TAMs.^10,29,30^ Given that GPAA1 knockout resulted in the loss of CD24 cell surface expression, we hypothesized that this would enhance macrophage-mediated phagocytosis of ovarian cancer cells. To investigate this idea, we conducted *in vitro* phagocytosis assays following established protocols.^10,31,32^ Parental, GPAA1 knockout or CD24 knockout OVCAR8 cells stably expressing green fluorescent protein (GFP) were labeled with a pH-sensitive fluorescent dye called pHrodo and co-cultured with macrophages that were derived from human peripheral blood mononuclear cells stimulated with M2-polarizing cytokines IL-4 and IL-13 to induce Siglec10 expression^10^ (Figure S2H). The total number of phagocytic events (pHrodo-positive cells) per well was quantified by imaging cytometry. The analysis revealed that GPAA1 knockout significantly enhanced phagocytosis of OVCAR8 cells (Figure 2G). Similar results were observed with GPAA1 knockout OVCAR3 and SKOV3 cells (Figure 2H). We confirmed these results using an alternative phagocytosis assay, in which phagocytic events were determined by flow cytometry to identify cells that were double positive for GFP and the macrophage marker CD11b, indicative of macrophage-engulfed cancer cells (Figures 2I and S2I). Finally, using A2780 cells in which CD24 expression is absent, GPAA1 knockout did not enhance macrophage-mediated phagocytosis (Figures S2J and S2K), strongly suggesting that GPAA1 regulates cancer cell phagocytosis specifically through CD24-dependent mechanism. Collectively, these results indicate that genetic depletion of GPAA1 not only abrogates CD24 cell surface expression but also enhances macrophage-mediated phagocytosis of ovarian cancer cells.

### GPAA1 knockout suppresses the growth of ovarian tumors and increases survival in mice

Previous studies have shown that genetic depletion of CD24 or disruption of the CD24-Siglec10 axis enhances *in vivo* phagocytosis and suppresses tumor growth in mice.^10,30,33^ Based on our results with the *in vitro* phagocytosis assays, we hypothesized that GPAA1 knockout ovarian cancer cells would be susceptible to TAM-mediated phagocytosis *in vivo*, which could potentially inhibit tumor growth. To test this idea, we intraperitoneally implanted parental, GPAA1 knockout or CD24 knockout OVCAR8 cells expressing GFP and luciferase into female NOD *scid* gamma (NSG) mice and monitored *in vivo* phagocytosis^10^ and tumor growth (Figure 3A). To assess *in vivo* phagocytosis, three weeks post-implantation, peritoneal fluid was harvested and analyzed by flow cytometry to identify cells that were double positive for GFP and the murine macrophage marker F4/80, indicative of macrophage-engulfed OVCAR8 cells (see Figure S3A). Our results revealed that GPAA1 knockout OVCAR8 tumors exhibited increased levels of *in vivo* phagocytosis by infiltrating TAMs, comparable to those observed in mice implanted with CD24 knockout cells (Figure 3B). Consistent with previous findings showing a pro-inflammatory polarization of macrophages following loss of CD24^10^, we observed a higher occurrence of pro-inflammatory M1-like macrophages (CD80+) and reduced frequency of M2-like macrophages (CD206+) in the peritoneal fluid of mice implanted with GPAA1 knockout OVCAR8 cells compared to those implanted with parental OVCAR8 cells (Figures S3B and S3C).

**Figure 3.**
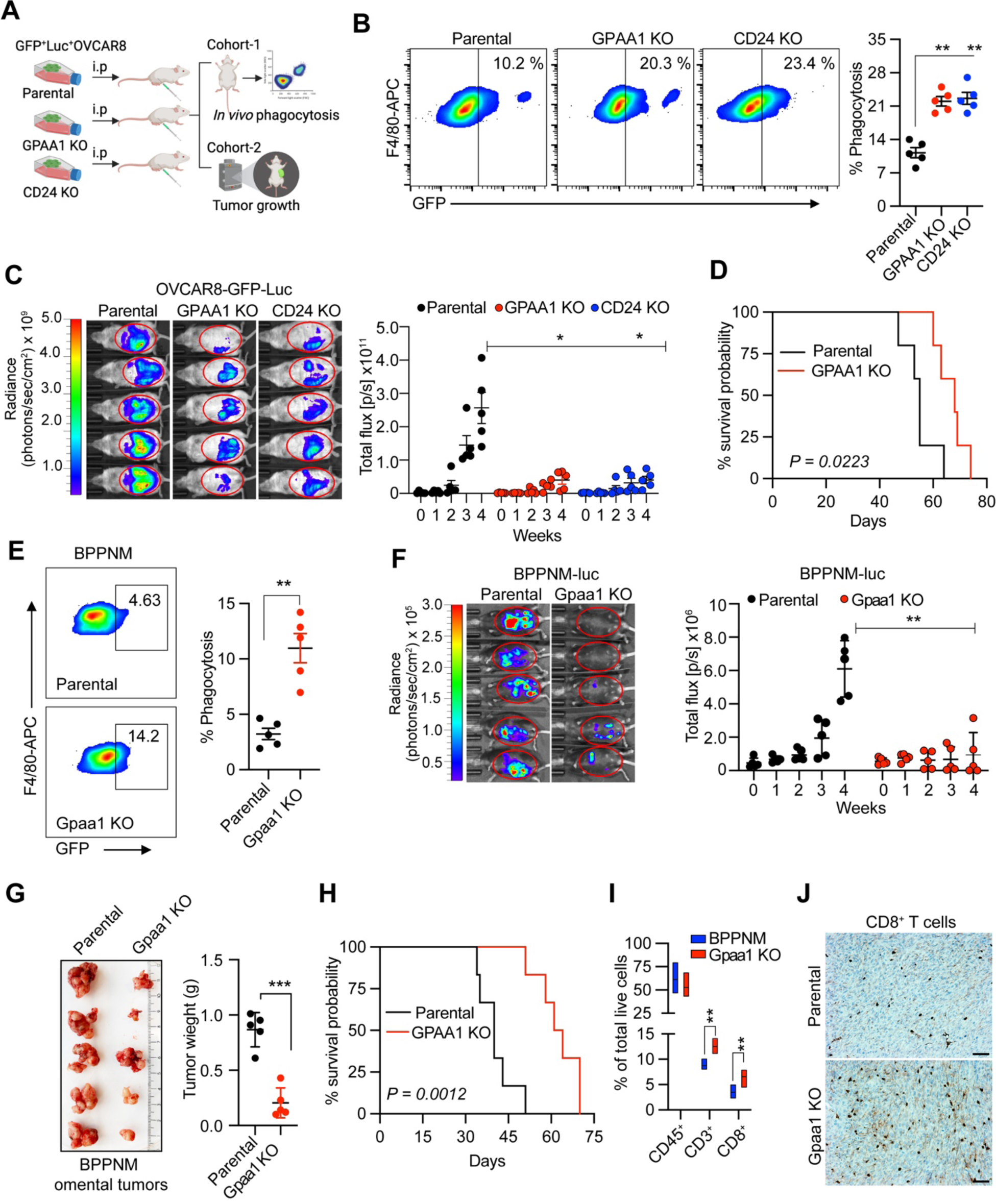
GPAA1 knockout suppresses the growth of ovarian tumors and increases survival in mice. (A) Schematic outline of *in vivo* phagocytosis and tumor growth monitoring in mice, created with BioRender.com. (B) (Left) Representative flow cytometry plots demonstrating TAM-mediated phagocytosis of parental, GPAA1 KO or CD24 KO OVCAR8 tumor cells; the number indicates the frequency of phagocytosis events out of all TAMs. (Right) Quantification of phagocytic events out of all TAMs in tumors derived from parental, GPAA1 KO, or CD24 KO OVCAR8 cells 21 days after implantation. Error bars indicate SD (n=5 mice per group). P values were calculated using one-way ANOVA followed by Dunnett’s multiple comparisons test. (C) (Left) Representative *in vivo* bioluminescence images of NSG mice implanted with parental, GPAA1 KO or CD24 KO OVCAR8 cells. (Right) Quantification of tumor growth as measured by total flux. Error bars indicate SEM (n=5). P values were calculated using two-way ANOVA with multiple comparisons. (D) Survival analysis of NSG mice harboring tumors derived from implantation with parental or GPAA1 KO OVCAR8 cells. P value was calculated by a log-rank (Mantel-Cox) test (n=5). (E) (Left) Representative flow cytometry plots demonstrating TAM-mediated phagocytosis of parental or Gpaa1 KO BPPNM tumor cells; the number indicates the frequency of phagocytosis events out of all TAMs. (Right) Quantification of phagocytotic events out of all TAMs in tumors derived from parental, or GPAA1 KO BPPNM cells 21 days after implantation. Error bars indicate SD (n=5). (F) (Left) Representative *in vivo* bioluminescence images in C57BL/6 mice implanted with parental or Gpaa1 KO BPPNM cells. (Right) Quantitation of tumor growth, as measured by total flux. Error bars indicate SEM (n=5). P value was calculated using two-way ANOVA with multiple comparisons. (G) (Right) Image showing omental tumors derived from implantation of parental or Gpaa1 KO BPPNM cells. (Right) Quantification of tumor weight. Error bars indicate SEM (n=5). P value was calculated using an unpaired student’s t-test. (H) Survival analysis of C57BL/6 mice harboring tumors derived from implantation of parental or Gpaa1 KO BPPNM cells. P value calculated by a log-rank (Mantel-Cox) test (n=6). (I) Box plot showing Immunophenotypic quantification of CD45^+^, CD3^+^ or CD8^+^ immune cells in ascites derived from parental or Gpaa1 KO BPPNM tumors. Error bars indicate SD. P values were calculated using an unpaired student’s t-test. (J) Immunohistochemistry showing CD8+ T cell infiltration in omental tumors derived from implantation of parental or GPAA1 KO BPPNM cells. Scale bar, 10 μM. *\*P<0.05, **P<0.01, ***P<0.001.* See also Figure S3.

Tumor growth was measured weekly by *in vivo* bioluminescence imaging. We observed a significant decrease in the growth of tumors derived from implantation of GPAA1 knockout OVCAR8 cells compared to those derived from parental OVCAR8 cells (Figure 3C). This reduction in tumor growth was comparable to that observed upon implantation with CD24 knockout cells (Figure 3C). Additionally, mice harboring tumors derived from GPAA1 knockout cells had significantly increased survival relative to mice bearing tumors derived from parental OVCAR8 cells (Figure 3D). Consistent with a previous study^10^ showing that the reduction in the growth of CD24-deficient tumors was TAM dependent, we also observed that the ability of GPAA1 knockout cells to form tumors was abrogated by TAM depletion (Figures S3D and S3E), confirming that the decreased tumor growth of GPAA1 knockout cells was TAM dependent.

To further extend our results in an immunocompetent setting, we utilized a genetically defined syngeneic ovarian cancer mouse model. In this model, murine fallopian tube epithelial cells bearing genetic alterations typical of HGSOC are implanted into syngeneic immunocompetent mice^34^. Specifically, we used the cell line BPPNM (*Brca1*^−/−^*Trp53*^−/−*R*172*H*^*Pten*^−/−^*Nf1*^−/−^*Myc*^OE^)^34^ and generated a Gpaa1 knockout derivative using CRISPR-mediated genome editing (Figure S3F). As expected, we found that CD24 cell surface expression was undetectable in Gpaa1 knockout BPPNM cells (Figure S3G). Furthermore, akin to our findings in human ovarian cancer cell lines, Gpaa1 knockout BPPNM cells exhibited enhanced macrophage-mediated phagocytosis *in vitro* (Figure S3H). Next, we intraperitoneally implanted parental or Gpaa1 KO BPPNM cells, which stably expressed luciferase and GFP, into female C57BL/6 mice, and after three weeks of engraftment, we performed an *in vivo* phagocytosis assay. The results demonstrated that Gpaa1 knockout BPPNM cells exhibited elevated levels of *in vivo* phagocytosis by infiltrating TAMs compared to parental BPPNM cells (Figure 3E and S3I). In addition, we observed a substantial reduction in the growth of tumors derived from Gpaa1 knockout BPPNM cells compared to those derived from parental BPPNM cells (Figure 3F, 3G and S3J). Consistent with our observations in NSG mice, C57BL/6 mice bearing tumors derived from Gpaa1 knockout BPPNM cells demonstrated significantly improved survival compared to mice bearing tumors derived from parental BPPNM cells (Figure 3H). Collectively, these findings indicate that the genetic ablation of GPAA1 is sufficient to enhance *in vivo* phagocytosis and impede tumor growth in xenografted mice.

Previous studies have shown that targeting phagocytic checkpoints can stimulate a robust antitumor T-cell response through priming of T cells by macrophages^35^. Interestingly, correlation analysis using the Tumor Immune Estimation Resource (TIMER)^36^ showed that *CD24* and *GPAA1* expression were negatively correlated with CD8+ T cell infiltration in ovarian cancer (Figure S3K). To examine T cell infiltration in GPAA1 knockout tumors, we performed flow cytometric analysis of peritoneal fluid and immunohistochemistry in omental tumors. The analysis revealed a significant increase in total CD3+ T cells and cytotoxic CD8+ T cells within GPAA1 knockout tumors compared to tumors derived from parental BPPNM cells (Figures 3I and 3J). The elevated infiltration of CD3+ T cells and CD8+ T cells within GPAA1 knockout tumors underscores the subsequent activation of the adaptive immune response following enhanced phagocytosis.

### The metallo-aminopeptidase inhibitor bestatin binds to GPAA1 and impairs GPI anchoring

GPAA1 adopts a structural fold similar to that of proteins belonging to the metallo-aminopeptidase family^14,27^, which carry one or more co-catalytic metal ions (most often Zn). We therefore hypothesized that the function of GPAA1 in GPI lipid anchoring could be inhibited by small-molecule aminopeptidase inhibitors (APIs). To test this possibility, we assessed several small-molecule APIs, including actinonin^37^, amastatin^38^, acebilustat^39^, ARM-1^39^, bestatin^40^, DG051^41^, firibastat^42^, HFI-142^43^, SC-57461A^44^, and tosedostat (CHR-2797)^45^, for their ability to inhibit GPAA1 function and consequently reduce CD24 cell surface expression. Flow cytometry analysis in OVCAR8 cells following treatment with these APIs revealed that the compounds actinonin, ARM1, bestatin, HFI-142, and tosedostat significantly decreased CD24 cell surface expression (Figure S4A). Notably, actinonin, ARM1, bestatin, and tosedostat share similar pharmacophores, such as hydrophobic and/or aromatic groups and hydrogen bond acceptors (Figure S4B), indicating their similar interaction mechanism to their potential binding partners/receptors. As expected, none of these APIs altered the cell surface expression of the non-GPI-linked protein MHC1 (Figure S4C)

We selected bestatin (also known as Ubenimex) for further analysis due to its well-documented potency in inhibiting multiple metallo-aminopeptidases.^40^ Additionally, bestatin is a natural compound derived from *Streptomyces oliverticuli* and is known to be orally active, safe, and well-tolerated.^46,47^ We employed computational methods to explore the potential binding of bestatin and other APIs to GPAA1 (see Supplementary-File-Computational-Analysis.pdf with sections I-III for more detail and Supplementary Files CSF1 and CSF2 for accompanying data). A thorough analysis of known 3D structures of aminopeptidase-inhibitor complexes revealed that the compounds that we found to inhibit GPAA1 interact with the aminopeptidase active site via the metal ion(s), most likely Zn, located there. In this dominant arrangement, two hydrogen acceptor functional groups from the inhibitor compound encircle the metal ion. Notably, one Zn ion is the preferred interaction partner for the inhibitor molecule, even in aminopeptidases with two metal ions in the active site (Figure 4A). The molecular mechanics and dynamics calculation results, including blind molecular docking attempts without prior knowledge of the bestatin-binding regions using AutoDock Vina,^48^ show that the binding mode of bestatin and similar inhibitor compounds observed in aminopeptidase-inhibitor complex structures is the most plausible, from the energetic/binding energy point of view, for our published GPAA1 structural models, which include GPAA1^Zn^ (with one Zn ion in the active site) and GPAA1^ZnZn^ (with two Zn ions).^14^ We also find that all observed, clinically relevant mutations of GPAA1 (except for L291P) energetically destabilize GPAA1, further supporting an active functional role for GPAA1 in the genesis of GPI lipid anchored proteins.

**Figure 4.**
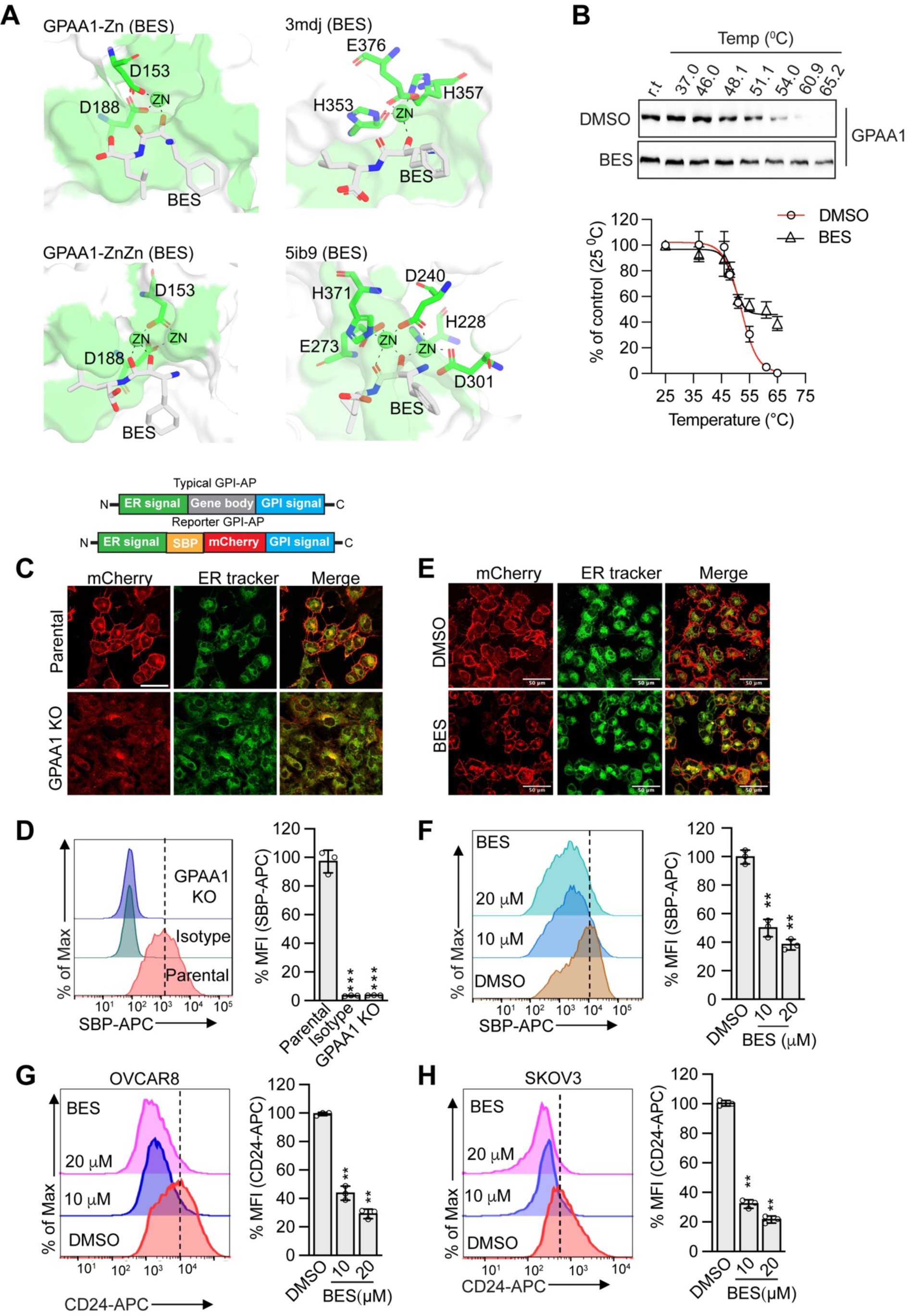
Bestatin binds to GPAA1 and inhibits GPIT activity. (A) Molecular modeling of bestatin binding modes in the GPAA1^Zn^ (upper left) and GPAA1^ZnZn^ (bottom left) models as compared to the referenced aminopeptidase-bestatin complexes PDB: 3mdj (one Zn; upper right) and PDB: 5ib9 (two Zn; upper right). The bestatin molecule is shown in sticks (carbon: white, nitrogen: blue, oxygen: red, others: yellow, hydrogen not shown for simplicity), the aminopeptidase pockets are shown in surface with the Zn in spheres. The pocket residues that are within 4 Å around bestatin are shown in green. The Zn-interacting residues are labeled. (B) Cellular thermal stability shift assay (CETSA) of GPAA1 in the presence of bestatin relative to that obtained with vehicle (DMSO). Immunoblot analysis with a GPAA1 antibody. (Bottom) The shift in bestatin binding to GPAA1 was analyzed by the Boltzmann sigmoid equation. All data were normalized to the response observed at DMSO-treated conditions at room temperature. (C) (Top) Schematics of a typical GPI-linked protein and the mCherry-SBP-tagged GPI reporter protein. (Bottom) Representative immunofluorescence images showing localization of mCherry in parental or GPAA1 KO OVCAR8 cells expressing the mCherry-SBP-tagged GPI reporter. ER tracker and merged images are shown. Scale bar, 50 µm. (D) (Left) Representative flow cytometry histograms of SBP cell surface expression in parental or GPAA1 KO OVCAR8 cells. (Right) Quantification of MFI. (E) Representative immunofluorescence images showing localization of mCherry in OVCAR8 cells expressing the mCherry-SBP tagged GPI reporter, treated with DMSO or 20 µM BES. ER tracker and merge images are shown. Scale bars, 50 µm. (F) (Left) Representative flow cytometry histograms of SBP cell surface expression by OVCAR8 cells treated with DMSO or 10 or 20 µM bestatin. (Right) Quantification of MFI. (G-H) (Left) Representative flow cytometry histograms of CD24 cell surface expression in OVCAR8 or SKOV3 cells treated with DMSO or 10 or 20 µM bestatin. (Right) Quantification of MFI. The results were normalized to that obtained in DMSO-treated cells. Data represents ± SD (n=3). P values were calculated using one-way ANOVA followed by Dunnett’s multiple comparisons test. *\*\*P<0.01, ***P<0.001.* See also Figure S4, supplementary-file-computational-analysisS1, supplementary-file-CSE1 and supplementary-file-CSE2.

In Figure 4A, the bestatin-binding pocket involving the GPAA1 Zn-binding residues (D153, D188, Y328 and/or E226)^27^ is illustrated for both the GPAA1^Zn^ and GPAA1^ZnZn^ models. This site was found to be the most frequently visited by the bestatin molecule in docking simulations. The bestatin molecule directly contacted the Zn(s) located in the GPAA1 active site of both GPAA1^Zn^ and GPAA1^ZnZn^, with a binding energy Δg (in kcal/mol) of −5.7 ± 1.04 and −5.14 ± 0.07, respectively. The same principal arrangement of a hydrogen bond acceptor pair from bestatin interacting with a Zn ion was also found in known 3D complexes such as PDB:3mdj (one Zn) or PDB:5ib9 (two Zn).

To confirm the physical interaction of bestatin with GPAA1, we utilized a cellular thermal stability shift assay (CETSA), a method for accessing drug–target interactions in cells^49^. OVCAR8 cells were treated with bestatin and then subjected to increasing temperatures to denature and precipitate proteins. Subsequently, cells were lysed, and the soluble protein fractions were analyzed by immunoblotting to quantify the changes in thermal stability. As shown in Figure 4B, the thermal stability of GPAA1 increased in the presence of bestatin compared to DMSO, suggesting that bestatin binds to GPAA1.

Specific biochemical assays for measuring the enzymatic activity of GPAA1 and components of the GPIT complex are not yet well established. Consequently, the functional characterization of GPIT components and the identification of factors that modulate GPIT activity have largely relied on the utilization of GPI-linked protein reporter assays.^50^ We therefore utilized this reporter assay to assess the impact of bestatin treatment on GPIT activity. For this assay, we stably expressed a GPI-anchored mCherry reporter protein (fused to a streptavidin-binding peptide tag, with an ER localization sequence at the N-terminus and a GPI attachment signal at the C-terminus; Figure 4C, top panel) in OVCAR8 cells and monitored mCherry localization by immunocytochemistry. As a control, we first assessed the localization of the GPI-anchored mCherry reporter protein in GPAA1 knockout cells. As anticipated, the mCherry signal was primarily observed at the cell surface in parental OVCAR8 cells, but in GPAA1 knockout cells, the mCherry signal was not detectable on the cell surface and was instead predominantly localized within the ER (Figure 4C). Moreover, the complete absence of cell surface expression of the SBP-mCherry-GPI reporter in GPAA1 knockout cells was also confirmed by flow cytometry (Figure 4D). Subsequently, we examined the impact of bestatin treatment on mCherry-GPI cell surface expression, revealing a notable reduction in cell surface presentation, as evidenced by confocal microscopy (Figure 4E) and flow cytometry (Figure 4F). These results suggest that bestatin inhibits GPAA1 function, impairing GPI anchoring.

Next, we evaluated the impact of bestatin on cell surface expression of CD24 in ovarian cancer cell lines. As expected, bestatin treatment led to decreased CD24 cell surface expression in OVCAR8 (Figure 4G) and SKOV3 (Figure 4H) cells. Notably, bestatin treatment did not affect cell proliferation of ovarian cancer cells (Figure S4D). Importantly, bestatin had no significant effect on total CD24 levels, as assessed by flow cytometry following cell permeabilization (Figure S4E), indicating that bestatin specifically decreases cell surface expression of CD24 without affecting intracellular CD24 levels. Finally, shRNA-mediated knockdown of either CD13 or LTA4H, two well-known targets of bestatin^51,52^, had no effect on CD24 cell surface expression (Figure S4F-I), strongly suggesting that the ability of bestatin to reduce CD24 cell surface expression is due to its inhibition of GPAA1.

### Bestatin, alone or in combination with docetaxel, inhibits growth of human ovarian cancer xenografts in mice

Because bestatin treatment reduced CD24 cell surface expression, we hypothesized that bestatin could, like GPAA1 depletion, augment phagocytosis of ovarian cancer cells and inhibit tumor growth. In an *in vitro* phagocytosis assay, we found an increase in macrophage-mediated phagocytosis of bestatin-treated OVCAR8, SKOV3, and OVCAR3 cells compared to DMSO-treated cells (Figure 5A-C). As expected, bestatin treatment of A2780 cells, which lack CD24 expression, did not result in a significant increase in phagocytosis compared to DMSO-treated cells (Figure 5D), confirming the specificity of bestatin in targeting CD24-dependent mechanisms.

**Figure 5.**
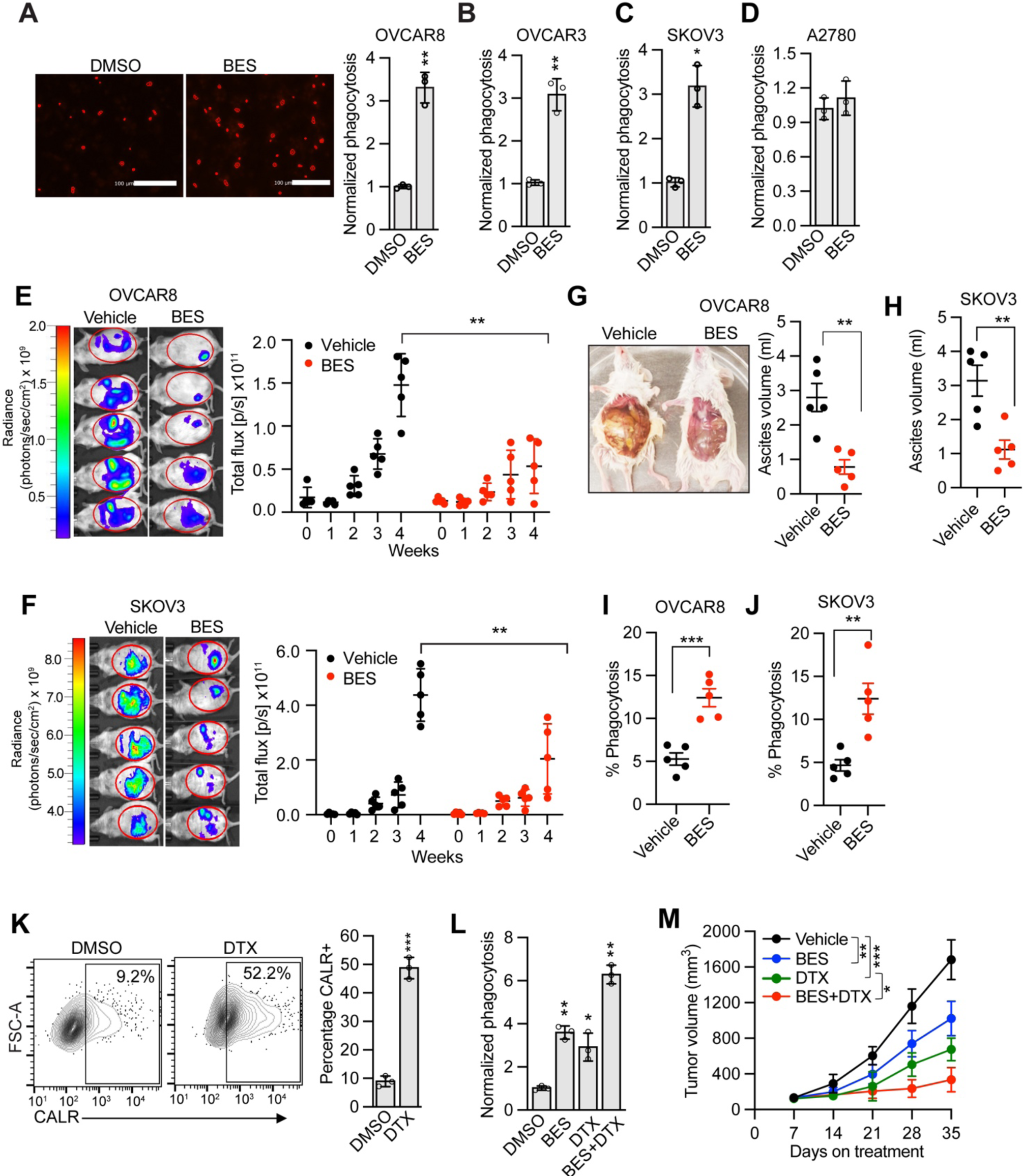
Bestatin treatment increases phagocytosis and inhibits the growth of human ovarian cancer xenografts in mice. (A) Representative images (left) and quantification (right) of macrophage-mediated phagocytosis of OVCAR8 cell treated with bestatin (20 µM) or, as a control, DMSO. The results were normalized to that obtained with DMSO. Scale bars, 100 µm. (B-D) Quantification of phagocytosis of OVCAR3 (B), SKOV3 (C), and A2780 (D) cells treated with bestatin (20 µM) or DMSO. Data represents ± SD (n=3). P values were calculated using an unpaired student’s t-test. (E and F) (Left) Representative *in vivo* bioluminescence images of NSG mice 28 days following implantation with OVCAR8 or SKOV3 cells and treated with 100 mg/kg/d bestatin or vehicle. (Right) Tumor growth, as measured by total flux. Error bars indicate SEM (n=5 mice per group). P values were calculated using two-way ANOVA with multiple comparisons. (G) (Left) Representative image depicting peritoneal ascites accumulation in mice 28 days following implantation with OVCAR8 cells and treated with either vehicle or bestatin. (Right) Quantification of ascites volume from vehicle or bestatin-treated mice (n=5 mice per group). (H) Quantification of ascites volume measurements in mice implanted with SKOV3 cells and treated with vehicle or bestatin. Error bars indicate SEM (n=5). P value was calculated using an unpaired t-test. (I-J) Quantification of TAM-mediated phagocytosis of OVCAR8 or SKOV3 tumor cells in mice treated with vehicle or bestatin. Error bars indicate SD (n=5). P values were calculated using an unpaired student’s t-test. (K) (Left) Representative flow cytometry analysis of CALR in OVCAR8 cells treated with docetaxel (DTX, 100 nM) or DMSO. (Right) Quantification of CALR-positive cells. Data represents ± SD (n=3). (L) Quantification of macrophage-mediated phagocytosis of OVCAR8 cells treated with BES, DTX or both in combination. Data represents ± SD (n=3). (M) Measurement of tumor growth in mice subcutaneously implanted with OVCAR8 cells, and treated with either vehicle, BES, DTX or both in combination. Error bars indicate SEM (n=5 mice per group). P values were calculated using P values were calculated using two-way ANOVA with multiple comparisons. *\*P<0.05, **P<0.01, ***P<0.001.* See also Figure S5.

To determine the effect of bestatin treatment on *in vivo* phagocytosis and ovarian tumor growth, we intraperitoneally implanted OVCAR8 or SKOV3 cells expressing GFP and luciferase in NSG mice. After one week, bestatin or vehicle was administered through intraperitoneal injection. A hallmark of ovarian cancer that is also observed in mouse models is the build-up of fluid (ascites) in the peritoneal cavity.^34^ As shown in Figures 5E and 5F, bestatin treatment resulted in a substantial reduction in tumor growth compared to the control vehicle-treated group. In addition, bestatin-treated mice exhibited significantly reduced ascites accumulation compared to vehicle-treated mice (Figure 5G and 5H). Furthermore, *in vivo* phagocytosis assays showed that bestatin-treated mice exhibited substantially increased GFP+ F4/80+ double-positive cells, indicative of macrophage-mediated engulfment of ovarian cancer cells (Figure 5I and 5J).

Combination therapies are often used to enhance the efficacy of treatment over monotherapies by targeting critical pathways synergistically or additively. We therefore hypothesized that simultaneous suppression of the CD24 phagocytic checkpoint and induction of a pro-phagocytic signal, such as surface localization of calreticulin (CALR),^53,54^ could augment phagocytosis and suppress tumor growth more effectively. To induce CALR, we used docetaxel, a conventional first-line chemotherapeutic drug for ovarian cancer^55,56^ that has been shown to induce surface localization of CALR. ^57^ Consistent with this idea, we found that docetaxel efficiently induced CALR cell surface expression in OVCAR8 cells, whereas two other chemotherapeutic agents, carboplatin and doxorubicin, did not (Figure 5K and S5). Moreover, combination treatment with bestatin and docetaxel promoted *in vitro* phagocytosis of OVCAR8 cells better than either treatment alone (Figure 5L). Finally, we assessed the impact of combined treatment with bestatin and docetaxel on tumor growth in mice. In this experiment, OVCAR8 cells were subcutaneously implanted into NSG mice and once tumors were established, the mice were treated with either vehicle, bestatin or docetaxel, or a combination of both drugs, and tumor volume was monitored weekly. The results showed that compared to single-agent treatment, combinatorial treatment significantly reduced tumor growth (Figure 5M). These findings suggest that activating the pro-phagocytic CALR signal may enhance the ability of bestatin to inhibit ovarian tumor growth.

## DISCUSSION

The CD24-Siglec10 axis, initially identified as an inflammatory response regulator in tissue injury^58^, now falls within the category of phagocytic immune checkpoint axes, akin to CD47-SIRPα^59^, B2M-LILRB1^60^, and PDL1-PD1^61^. Recent studies have highlighted CD24 as a potential immunotherapeutic target in various cancers.^10,29,62,63^ In the ovarian tumor microenvironment, CD24 is predominantly expressed in malignant tumor cells (see Figure 2A). In contrast, CD47, another anti-phagocytic signal, is widely expressed across all cell types^10^. Thus, targeting CD24 may offer the advantage of fewer off-target effects, potentially reducing the risk of toxicity^64^.

Preclinical studies targeting the CD24-Siglec10 axis using monoclonal antibodies (mAbs) or bispecific peptides have shown promising results, particularly in boosting phagocytic activity in CD24-positive tumors by TAMs.^10,33,65^ However, the therapeutic efficacy and safety of these interventions have not been tested in clinical settings. Although humanized mAbs targeting CD24 hold potential as cancer immunotherapeutics,^66^ small-molecule therapeutics offer several advantages, including better penetration into solid tumors, reduced immunogenicity, lack of Fc-mediated side effects often observed with mAbs, oral administration convenience, and reduced production costs.^67,68^

In this study, we identified GPAA1 as a factor that plays a crucial role in facilitating cell surface localization and expression of CD24 and showed that a small-molecule inhibitor of GPAA1, bestatin, reduced ovarian tumor growth by inducing macrophage-mediated phagocytosis. Notably, previous studies have reported that bestatin, or its derivative LYP, may reduce ovarian cancer growth by directly killing cancer cells through inhibition of aminopeptidase N (APN/CD13) activity.^69^ However, in our experiments, we did not observe any significant effect of bestatin on cell viability, even at very high concentrations, suggesting that the primary mechanism of bestatin-mediated tumor reduction in ovarian cancer is through an immune-mediated mechanism rather than a direct effect on cell viability. The results of our study highlight the potential of targeting GPAA1 as a therapeutic approach for CD24-positive ovarian cancers. In addition, we found that shRNA-mediated knockdown of other factors involved in GPI biosynthesis, such as PIGM, N, O, P, T and U, which were also identified from our primary screen, also led to a reduction in CD24 cell surface levels, suggesting the GPI biosynthesis pathway as a broader immunotherapeutic target. Interestingly, although the GPI biosynthesis pathway is crucial for the viability of yeast and protozoa, in mammals it is thought to be non-essential.^70^ Consistent with this idea, we observed that the knockout of GPAA1 in multiple ovarian cancer cell lines did not result in any proliferation defects. Moreover, the knockdown of other components of the GPIT complex, such as PIGM, N, O, P, T, and U, was well tolerated in ovarian cancer cells. Notably, like GPAA1, several other components of the GPI pathway are aberrantly expressed in many cancers, including ovarian cancer.^22^ Many of these GPI pathway proteins are enzymatic and therefore of potential therapeutic interest. However, despite attempts to target the GPI pathway in fungal and protozoan pathogens^70–72^, targeted inhibition of the pathway in mammalian systems has proven difficult.^72^ To date, targeting of GPI pathway proteins has not yet been explored in the context of cancer.

Prompted by previous studies showing that GPAA1 has a catalytic site that is structurally similar to that of metallo-aminopeptidases^29^ and is suggested to function as a metallo-peptide synthetase catalyzing the creation of a peptide bond between the carboxyl group of the substrate protein and the phosphoethanolamine of the mature GPI lipid anchor,^27^ we anticipated that the inhibition of this potential enzymatic function could be achieved using commercially-available APIs. Our results convincingly show that the implied GPAA1 function can indeed be pharmacologically targeted by several APIs including bestatin, tosedostat and ARM1. Because the three compounds share similar pharmacophores such as hydrophobic and/or aromatic groups and hydrogen bond acceptors, their interaction mechanism with GPAA1 is most likely similar. Indeed, our computational docking efforts of bestatin and GPAA1 recovered molecular arrangements observed in known 3D structures of aminopeptidase-inhibitor-complexes in which a metal ion is encircled by hydrogen acceptor groups of the inhibitor and the compound is embedded in a binding pocket that is collocated with the active site.

Our work provides the first experimental evidence for the theoretically deduced metallo-peptide synthetase activity of GPAA1.^27^ Notably, our results are in agreement with previous studies that convincingly associate GPAA1 with the process of GPI lipid anchor attachment (a process that was shown to be independent and separable of cleavage of the C-terminus of the substrate protein by the caspase-like protease PIGK, which generates the substrate for GPI anchor attachment)^13,18,20,73^ and with 3D structural data for GPAA1 (except for some details including the Zn ions near the active site, which might result from issues of sample preparation and structural resolution; see Supplementary-File-Computational-Analysis.pdf with sections IV and V for more data and discussion).

At the same time, our experimental findings in this study and previous work^14,27^ are in conflict with two published, almost identical cryo-EM structures of the GPIT complex^19,50^ that cannot rationalize a catalytic involvement of GPAA1 in the process of GPI lipid anchor attachment. How can this conflict be resolved? First, recent work by Ness *et al.* developed an *in vitro* enzymatic assay for GPI transferase activity^74^, which reaffirmed GPAA1 as an essential subunit within the minimal complex with GPI lipid transferase activity. They advanced the idea of dimerization, which speculated that the process might involve the spatial vicinity of active sites of PIGK and GPAA1 from different complex entities. Second, it can also not be excluded that previously published cryo-EM structures do not represent catalytically active GPIT conformations as a result of sample preparation or certain assumptions in the structure modeling. Supporting the latter point of view is a recently reported structure of the GPIT complex together with model ligands, which shows PIGK and GPAA1 in cooperating vicinity and with metal ions interacting with the phosphoethanolamine attached to the GPI lipid anchor. ^73^ A direct enzymatic assay with an *in vitro* reconstructed GPIT complex will be necessary to evaluate the activity of individual GPIT subunits and the mechanism of action of APIs that disrupt GPIT function.

In summary, the findings of this study highlight the potential for therapeutic targeting of CD24 through GPAA1 inhibition using small-molecule APIs. Our results encourage further exploration of the GPI pathway as a promising target for small molecule-based immunotherapeutic strategies. Although our research primarily focused on ovarian cancer, it is essential to recognize that GPAA1-mediated CD24 regulation is not limited to ovarian cancer. Consequently, targeting GPAA1 and other components of the GPI synthesis pathway could be a feasible strategy for various CD24+ cancers, including breast, lung and colorectal cancers^9^.

## Supporting information

Supplementary-file-computational analysis

Supplementary -File -CSE1

Supplementary -File -CSE2

## ACKNOWLEDGEMENTS

This work is dedicated to the memory of our mentor Michael R. Green, who suddenly passed away while this manuscript was in progress. His guidance and scientific insights were highly invaluable to this work. We would like to express our gratitude to Michelle Kelliher and Arthur Mercurio for their generous support and for critically reading the manuscript. We thank Robert. A. Weinberg for providing the BPPNM cell line; Surajit Sinha for providing the CMV51p>ffluc2-mEmerald plasmid; Magnolia L. Pak for advice on the CRISPR screen; Huseyin Mehmet for advice on this study; Kishore K. Srivastava for critically reading the manuscript; and Madison Coleman and Lynn Chamberlain for technical assistance. We also acknowledge support from the University of Massachusetts Chan Medical School (UMCMS) RNAi Core Facility for providing shRNAs; the UMCMS FACS Core Facility; and the UMCMS Deep Sequencing Core Facility.

## MATERIAL & METHODS

### Cell lines

Human ovarian cancer cell lines A2780, A1847, IGROV3, OVCAR3, OVCAR4, OVCAR8, SKOV3, and NCI/ADR-RES were cultured in RPMI 1640 medium. HEK293T cell line was maintained in Dulbecco’s Modified Eagle Medium (DMEM) with high glucose media supplemented with 10% fetal bovine serum (FBS), sodium pyruvate, non-essential amino acids, 100 units/ml penicillin and 100 µg/ml streptomycin. All cell lines were cultured at 37°C and 5% CO_2_. Sublines derived from OVCAR3, OVCAR8, and SKOV3 were cultured in the same conditions as the parental cell lines.BPPNM (*Brca1*^−/−^*Trp53*^−/−*R*172*H*^*Pten*^−/−^*Nf1*^−/−^*Myc*^OE^) cell line and its sublines were cultured DMEM supplemented with 1% insulin–transferrin–selenium (ITS-G), EGF (2 ng/mL), 4% heat-inactivated fetal bovine serum, and 100 units/ml penicillin and 100 µg/ml streptomycin

### Lentivirus packaging, transduction, and shRNA knockdown

For packaging lentiviral shRNAs, 1.0×10^6^ HEK293T cells were transfected with lentiviral transgene vectors and packaging plasmids psPAX2 and pMD2.G were mixed in a 2:2:1 ratio. Transfection was performed using Effectene transfection reagent (Qiagen). The next day, the medium was replaced to remove DNA complexes. After 48 hours, post-transfection medium containing lentiviral particles was collected and filtered through a 0.45 μm filter. For shRNA knockdown, 1×10^5^ cells per well were seeded in 6-well plates and transduced with 500 µl lentivirus particles (∼2×10^6^ TU/ml) packaged with an shRNA-expressing TRC lentivirus vector in a total volume of 1 ml of appropriate medium supplemented with 8 µg/ml polybrene. The medium was replaced after overnight incubation to remove polybrene and viral particles, and cells were incubated for another 24 hours and then subjected to puromycin selection (2 µg/ml) for 3-5 days. The shRNA sequences are listed in the Key Resource Table.

### CRISPR/Cas9 screen

To generate a stable Cas9-expressing OVCAR8 cell line, the plasmid lentiCas9-blast was packaged into a lentivirus as described above, and the viral supernatant was used to transduce OVCAR8 cells. Cells were selected for 5 days with blasticidin (5 µg/ml) and single-cell clones were isolated and tested for Cas9 gene editing efficiency using the Cas9 reporter vector pXPR_011. For the screen, 8×10^7^ OVCAR8/Cas9 cells were transduced with the Human Brunello CRISPR Knockout Pooled Library^16^ at a multiplicity of infection (MOI) of 0.5. Cells were selected with 2 µg/ml puromycin for 15 days and then stained with an APC-conjugated anti-human CD24 antibody for 30 min on ice in the dark. Five to 10 min prior to FACS, 7-AAD Viability Staining Solution was used to exclude dead cells from the analysis. At least 1×10^8^ CRISPR-edited cells were FACS sorted to isolate the CD24^low^ (defined as cells with the 5% lowest CD24 staining) and 7AAD^neg^ population. Total genomic DNA was extracted from the CD24^low^ and unsorted populations and 20 µg was used to prepare a next-generation sequencing library as previously described ^83^, which was sequenced using Illumina technology. The quality of the raw reads was assessed using FastQC (version 0.11.5). The 20-bp sequences immediately following the pre-guide RNA sequence “GGCTTTATATATCTTGTGGAAAGGACGAAACACCG” was extracted using a customized Perl script. For mapping the 20 bp reads to the human Brunello library’s single guide RNA (sgRNA) sequences, Bowtie (version 1.2.2) was employed with the default parameters, except for -m 1 –best -v 3^75^. To identify candidate genes for further validation and analysis, Fisher Exact test was performed, and P-values were adjusted using the BH-method to counteract the effects of multiple hypothesis testing. The statistical analysis and generation of figures were carried out using the *CRISPRscreen* R package, available at https://github.com/LihuaJulieZhu/CRISPRscreen. Gene Ontology (GO) enrichment analysis was performed using the function get enriched GO in Bioconductor package ChIPpeakAnno. GO terms with P-value < 0.001 were considered significant.

### Analysis of *GPAA1* genomic amplification and Kaplan-Meier analysis

Pan-cancer analysis of genomic amplification of *GPAA1* was performed using the online cBioPortal database (http://www.cbioportal.org). To generate the survival curves for patients expressing high (upper 25%) versus low (other 75%) levels of *GPAA1*, the interactive web server Online consensus Survival analysis for Ovarian cancer (OSov) found at the Long-term Outcome and Gene Expression Profiling Database of pan-cancers (LOGpc) (https://bioinfo.henu.edu.cn/OV/OVList.jsp) was used to query the dataset GSE63885 (for GPAA1).

### Analysis of scRNAseq dataset

The Bioturing Talk2data online interactive web tool was used to query the publicly available single-cell RNA (scRNA) dataset (GSE165897) as previously described^76^. Default parameters were utilized to create tSNE plots illustrating the expression of CD24 and GPAA1 using BioVinci tool. Cell clusters in the ovarian cancer tumor microenvironment (TME) were identified based on author-defined cell types defined^84^.

### CRISPR knockout generation

For GPAA1 and CD24 CRISPR-mediated knockouts, sgRNAs targeting the *GPAA1 and CD24* genes were cloned into the vector lenti-CRISPRv2. The sgRNA constructs (listed in Key Resource Table) were packaged into a lentivirus as described above. Cells were transduced with lentiviral particles and selected in puromycin (2 mg/ml) for 10 days. Single-cell clones were isolated using the serial dilution method in 96-well plates. Individual single-cell clones were grown, and knockout was confirmed by immunoblotting using a polyclonal GPAA1 or CD24 antibody.

### Flow cytometry analysis

For cell surface protein staining, 1×10^5^–1×10^6^ cells were first incubated with Fc receptor blocking solution, for 10 minutes at room temperature. Cells were washed once in FACS buffer (1X PBS, 1.0 % bovine serum albumin (BSA) and 0.2% sodium azide, and then incubated with antibodies for 30 minutes at 4°C in the dark. Cells were also stained with 7-AAD or DAPI for dead cell exclusion, and flow cytometry analysis was performed using a Bio-Rad ZE5 Cell Analyzer. For intracellular staining, cells were fixed with 1% paraformaldehyde for 5 min and permeabilized using the True-Nuclear Transcription Factor Buffer Set before the addition of the antibody.

### Quantitative RT-PCR

Total RNA was extracted from cells using Trizol. The cDNA was synthesized using Proto Script II reverse transcription kit (NEB) and real-time PCR reactions were performed using Quant Studio 3 (Applied Biosystems) using primer sequences listed in Key Resource Table. Expression was normalized to that of *GAPDH*. Experiments were performed three independent times and the results from one representative experiment are shown in technical triplicate.

### Immunoblot analysis

Cells seeded in 6-well plates were harvested and lysed in RIPA buffer (1% Triton-X100, 0.1% SDS, 0.5% deoxycholic acid, 20 mM Tris, pH 7.5, 10% Glycerol) containing 1X protease inhibitor cocktail and 1 mM PMSF. Total lysates (30 µg) were run on 10% SDS-PAGE and transferred to nitrocellulose membrane. Membranes were incubated with anti-GPAA1 (1:1000 dilution), anti-CD24 (1:1,000 dilution), or anti-β-actin (1:2000 dilution) antibodies overnight at 4°C. The blots were imaged by exposing them to x-ray film. Experiments were conducted three independent times and the results from one representative experiment are shown.

### Confocal microscopy

To prepare cells for confocal microscopy, cells were grown on coverslips and fixed with 4% paraformaldehyde for 20 minutes at room temperature followed by three washes with cold 1X PBS. Cells were then permeabilized with 0.3% Triton X-100 for 10 minutes, washed three times with cold PBS, blocked for 1 hour at room temperature with 10% goat serum in PBS, and then incubated at 4°C overnight with recombinant anti-CD24 primary antibody. The cells were then washed three times with PBS, incubated with a fluorescently conjugated secondary antibody (goat anti-rabbit IgG H&L (Alexa Fluor 594) for 1 hour at room temperature, and then washed again three times with PBS. For ER staining, cells were incubated with ER-Tracker Blue-White DPX in PBS for 10 minutes at room temperature. Thereafter, cells were washed three times with PBS before being mounted in ProLong Gold Antifade Mountant. Images were taken using a Leica SP8 Laser Scanning Confocal Fluorescence microscope using a 100X objective lens (oil). The z-stack images were taken with a 0.2 mm thick frame. Images were processed and analyzed using Fiji software.

### Small-molecule inhibitor and drug treatments

Ovarian cancer cells were seeded in 96-well plates, and when reached 70% confluency, cells were subjected to treatment with aminopeptidase inhibitors for 48 h with the concentrations as indicated in the figure legends. Dimethyl sulfoxide (DMSO) served as the vehicle control. In case of chemotherapeutic drugs Docetaxel (100 nM), Carboplatin (100 nM), and Doxorubicin (50 nM), cells were treated for a duration of 12 hours.

### Docking analyses of bestatin to the GPAA1 active sites

Ten replicates were extracted from the molecular dynamics (MD) trajectories of the Zn(s)-bound GPAA1 in our previous study^14^ and used to first explore the potential BES-binding pocket on the GPAA1^Zn^ and GPAA1^ZnZn^ structures. Blind docking was first initiated on the 10 replicates (10×1000 binding modes) using AutoDock Vina^48^ with a grid box centering and covering the whole GPAA1 structure (as receptor). The bestatin molecule retrieved from PubChem (CID: 72172) was used as the ligand. Only those pockets that involved the GPAA1 active sites and the Zn atom(s) and that were most frequently visited by the ligand were of interest. The previous 3×300 ns MD samplings of the GPAA1^Zn^ and GPAA1^ZnZn^ models were extended to 3×500 ns using the same parameters as previously. Ten replicates were extracted from the last 300 ns of the new sampling for further docking refinement. In the subsequent focus docking refinement, we used Glide version 8.8 implemented in the Schrödinger package, release 2020-3.^79^ The bestatin molecule was prepared and minimized using the LigPrep module. Then, it was docked attentively to the detected region of interest above, i.e., the proximity of the Zn(s)-bound active sites of the GPAA1 structures resulted from the blind docking experiment. For the receptor, a grid box of 7 Å (or 10 Å) was set centering the Zn (or Zns, respectively) coordinated with the GPAA1 active sites that include D153, D188, Y328 and/or E226.^14^ The docking protocol with default settings was performed on the extracted 10 replicates of the GPAA1^Zn^ or GPAA1^ZnZn^ conformations. Only those resulting docked complexes with the bestatin binding modes satisfying the detected pharmacophores (i.e., similar to those in the controls PDB: 3mdj and 5ib9) were selected for further analyses. The binding energies were estimated using the PRODIGY-LIG server.^80^

### Cellular Thermal Shift Assay (CETSA)

CETSA assays were performed as described previously.^49^ In brief, 1×10^7^ OVCAR8 cells were treated with DMSO or 100 μM bestatin for 2 hours. Cells (50 µl) were aliquoted in PCR tubes and incubated at room temperature (25°C) or at a range of temperatures (37°C to 65.2°C) in a thermal cycler for 3 min and cooled to 4°C. Cells were lysed using Alpha SureFire Ultra Lysis Buffer supplemented with Protease Inhibitor Cocktail. Insoluble proteins were separated by centrifugation at 15000xg for 30 mins at 4°C. Equal volumes of each sample were separated by SDS-PAGE and detected by immunoblotting blotting using an anti-GPAA1 polyclonal antibody. Three independent experiments were performed. Densitometry of the blots was performed using ImageJ software and the graph was plotted using GraphPad Prism 9.0.

### Macrophage generation and stimulation

Primary human donor-derived macrophages were generated as described previously.^85^ In brief, leukopaks from anonymous donors were obtained from the Rhode Island Blood Center (Providence, RI) and peripheral blood monocytes (PBMCs) were isolated by density gradients centrifugation using Ficoll Paque Plus followed by RBC lysis using RBC lysis buffer. For monocyte isolation, PBMCs were incubated in Monocyte Attachment Medium for 1 hour at 37°C and 5% CO_2_ and then washed three times in Iscove’s Modified Dulbecco’s Medium (IMDM) to remove non-adherent cells. The remaining monocytes were detached and seeded in 96-well plates and cultured at 37°C and 5% CO_2_ in IMDM supplemented with 10% FBS and 20 ng/ml recombinant human M-CSF. After 4 days monocytes were then treated with 20 ng/ml recombinant human IL-4 and 10 ng/mL recombinant human IL-13 for another 3-4 days to obtain M2 polarized macrophages. Macrophage differentiation was confirmed morphologically using microscopy. M2-macrophage polarization was confirmed by immunophenotyping of the M2 marker CD206 using PE-conjugated anti-human anti-CD206 antibody.

### *In vitro* phagocytosis assays

Ovarian cancer cells were labeled with pHrodo Red on ice for 30 min and then co-cultured with PBMC-derived M2 macrophages (prepared as described above) at a ratio of 1:1 (ovarian cancer cells: macrophages) for 4 hours in a humidified 5% CO_2_ incubator at 37°C. Phagocytic events were quantified by counting the number of red fluorescent (pHrodo+) cells per well using a Celigo Imaging Cytometer. GFP was used for the visualization of cancer cells in imaging. Phagocytic events were also quantified using a previously described flow cytometry-based assay.^10,65^ In brief, M2-like macrophages co-cultured with EmGFP+ cancer cells were detached using TrypLE Express, stained with APC anti-human CD11b and analyzed by flow cytometry. The percentage of macrophages undergoing phagocytosis (eaters) was calculated as the percentage of CD11b+ EmGFP+ double-positive cells in the population. For phagocytosis assay with mouse cells (BPPNM), Murine RAW264.7 (RAW) macrophages were cultured in RPMI supplemented with 10% fetal bovine serum and 2% penicillin/streptomycin. For differentiation, cells were stimulated with M-CSF (20 ng/ml) for 48 h, followed by a 72 h stimulation of IL4 (40 ng/ml) and IL13 (40 ng/ml) for M2 polarization.

### Tumor measurement

6-7-week-old female NOD *scid* gamma (NSG) or C57BL/6 mice were purchased from the Jackson Laboratory. All mouse studies were performed in accordance with the Guide for the Care and Use of Laboratory Animals from NIH, and a protocol (202000157) approved by the UMass Chan Medical School Institutional Animal Care and Use Committee (IACUC).

Female NSG mice, 6-7 weeks of age, were injected intraperitoneally with either OVCAR8 (4×10^6^ cell/mouse) or SKOV3 (5×10^6^ cells/mouse) EmGFP-luc cells or the corresponding GPAA1knockout EmGFP-luc sublines. Similarly, C57BL/6J mice were implanted with luciferase expressing syngeneic BPPNM cells (4×10^6^ cells/mouse). Tumors were analyzed using bioluminescence imaging beginning 7 days post-engraftment and continuing every 7 days until day 28. Mice were injected intraperitoneally with luciferin at 140 mg/kg in PBS and images were acquired 10 min later using an IVIS Spectrum CT In Vivo Imaging System. Total flux was quantified using Living Image 4.0 software. For survival analysis, deaths were recorded as instances when the tumor burden reached 10 cm of abdominal circumference and/or the body condition scoring values dropped below the threshold specified in our IACUC protocols.

For bestatin treatment, 5×10^6^ OVCAR8 or SKOV3 cells were implanted in the peritoneal cavity of mice as described above (n=5 per group), and 7 days later bestatin (100 mg/kg) was administered daily by intraperitoneal injection for 4 weeks. Tumors were analyzed by *in vivo* bioluminescence imaging as described above. For the drug combination experiment, 5×10^6^ OVCAR8/EmGFP-luc cells were mixed with Matrigel in a 1:1 volume and were implanted subcutaneously in the right flank of NSG mice (n=5 per group). When tumors reached ∼100-150 mm^3^, mice were treated with either vehicle control (0.9% saline), bestatin (100 mg/kg twice daily by oral gavage), docetaxel (5 mg/kg per week by tail vein injection) or a combination of both bestatin and docetaxel for 28 days. Tumors were measured every week using digital callipers and tumor volumes were calculated using the formula (V = ½ (Length × Width^2^). Differences between control and treated groups were determined using the two-way ANOVA followed by Turkey’s multiple comparison test.

### Macrophage depletion

For macrophage depletion experiments, 200 μl of clodronate liposomes was administered intraperitoneally every 4 days for 16 days before tumor implantation, followed by injection of 100 μl of clodronate liposomes every 4 days. Depletion of macrophages was verified by flow cytometry analysis peritoneal lavage using an APC anti-mouse F4/80^+^ antibody.

### *In vivo* phagocytosis assay

An *in vivo* phagocytosis assay was performed as described previously.^10^ Mice were implanted with EmGFP+ ovarian cancer cells as described above and 7 days later bestatin (100 mg/kg) was administered daily by intraperitoneal injection. After 2 weeks of bestatin treatment, peritoneal ascites was harvested. Single-cell suspensions of ascitic cells were blocked using TruStain FcX (anti-mouse CD16/32 antibody for 10 min at room temperature and then stained with an APC anti-mouse F4/80 antibody for 30 min on ice in the dark. The percentage of cells undergoing phagocytosis was calculated as the percentage of F4/80+ EmGFP+ double-positive cells in the population.

### Immunophenotyping

The omental tumors harvested from C57Bl/6 mice were minced and dissociated using the Human Tumor Dissociation Kit from Miltenyi Biotec, following the manufacturer’s instructions. Single cell suspensions were filtered, and cells were subsequently resuspended in FACS buffer. Cells were stained with anti-mouse; CD45-Pac-Blue, CD3-PE, and CD8-APC antibodies and analyzed by flow cytometry. DAPI was used for viability assessment.

### Immunohistochemistry

Tumor samples were collected from mouse xenografts, preserved in 10% buffered formalin phosphate overnight, and subsequently embedded in paraffin. Blocks were sliced into 5 µm sections slides and stained with an anti-CD8a antibody (at a 1:100 ratio). Morphological examinations were performed at the UMass Chan Medical School Morphology Core Facility.

### Cell proliferation assays

Ovarian cancer cells were transduced with a lentivirus expressing non-silencing (NS) or CD24 shRNA (see the key resource table) and analyzed for cell viability using PrestoBlue Cell Viability Reagent as per the manufacturer’s instructions, and relative cell growth was determined. Relative cell viability of OVCAR8 or SKOV3 cells treated with either DMSO or bestatin 10 and 50 mM was determined using a PrestoBlue assay. For crystal violet staining, OVCAR8, OVCAR3 or SKOV3 cells (1 X10^4^) were seeded in a 6-well plate and cultured for 5-6 days. Cells were washed with PBS and fixed using 1 ml of 4% paraformaldehyde for 20 minutes at room temperature. Following two rinses with PBS, the cells were stained with 0.5ml of 0.1% crystal violet solution (in 10% ethanol) for 20 minutes. The cells were rinsed twice with PBS, and the plate was scanned.

### Statistical analysis

Statistical analyses were performed as described in the corresponding figure legends. Statistical significance was determined by one-way ANOVA or two-way ANOVA followed by multiple comparisons test or unpaired Student’s t-test or log-rank test with GraphPad Prism 9. Data represented as ± SD of at least 3 independent biological replicates, and sample numbers (n) are indicated in the figure legends. *\*P<0.05, **P<0.01, ***P<0.001, ****P<0.001*

## KEY RESOURCES TABLE

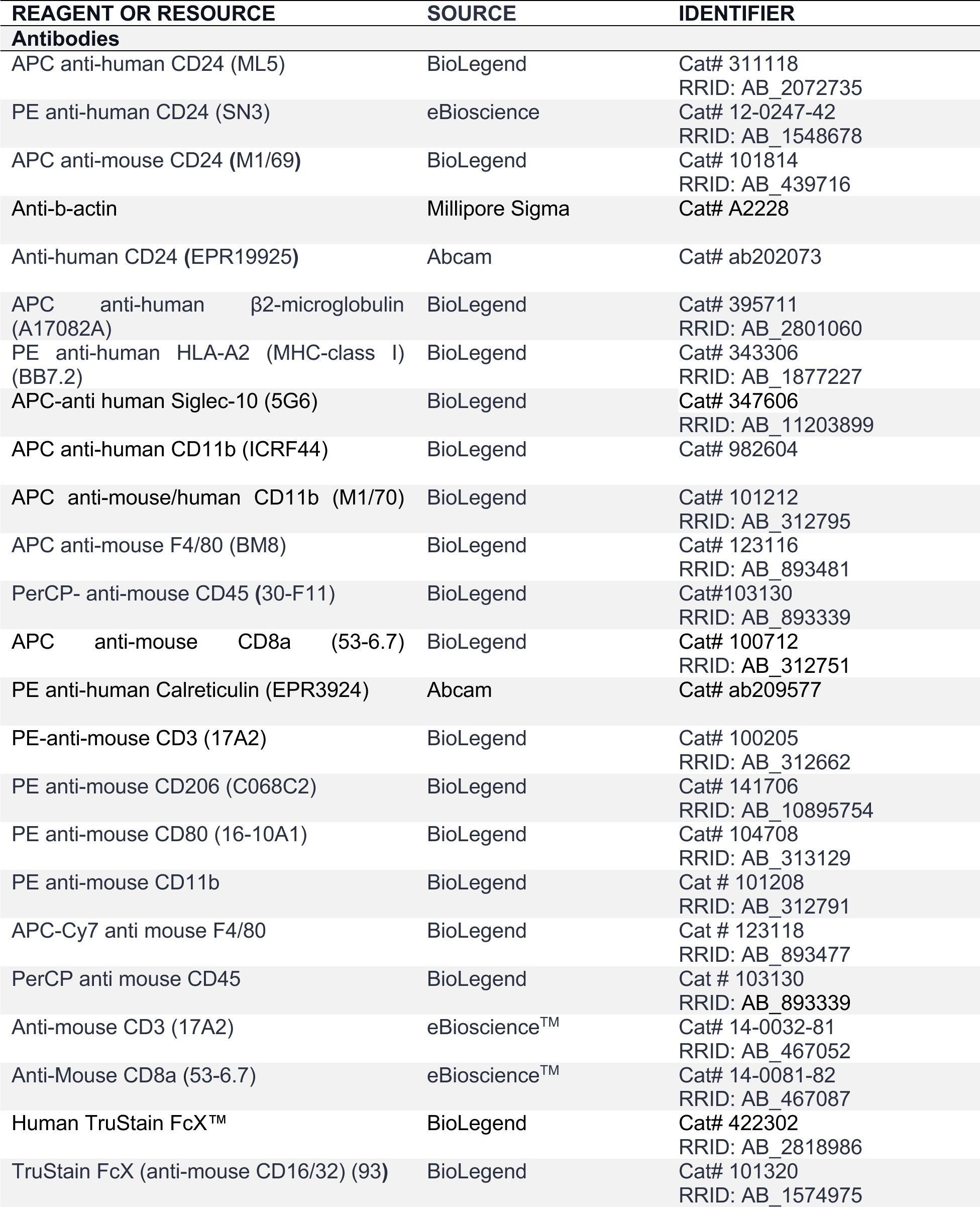
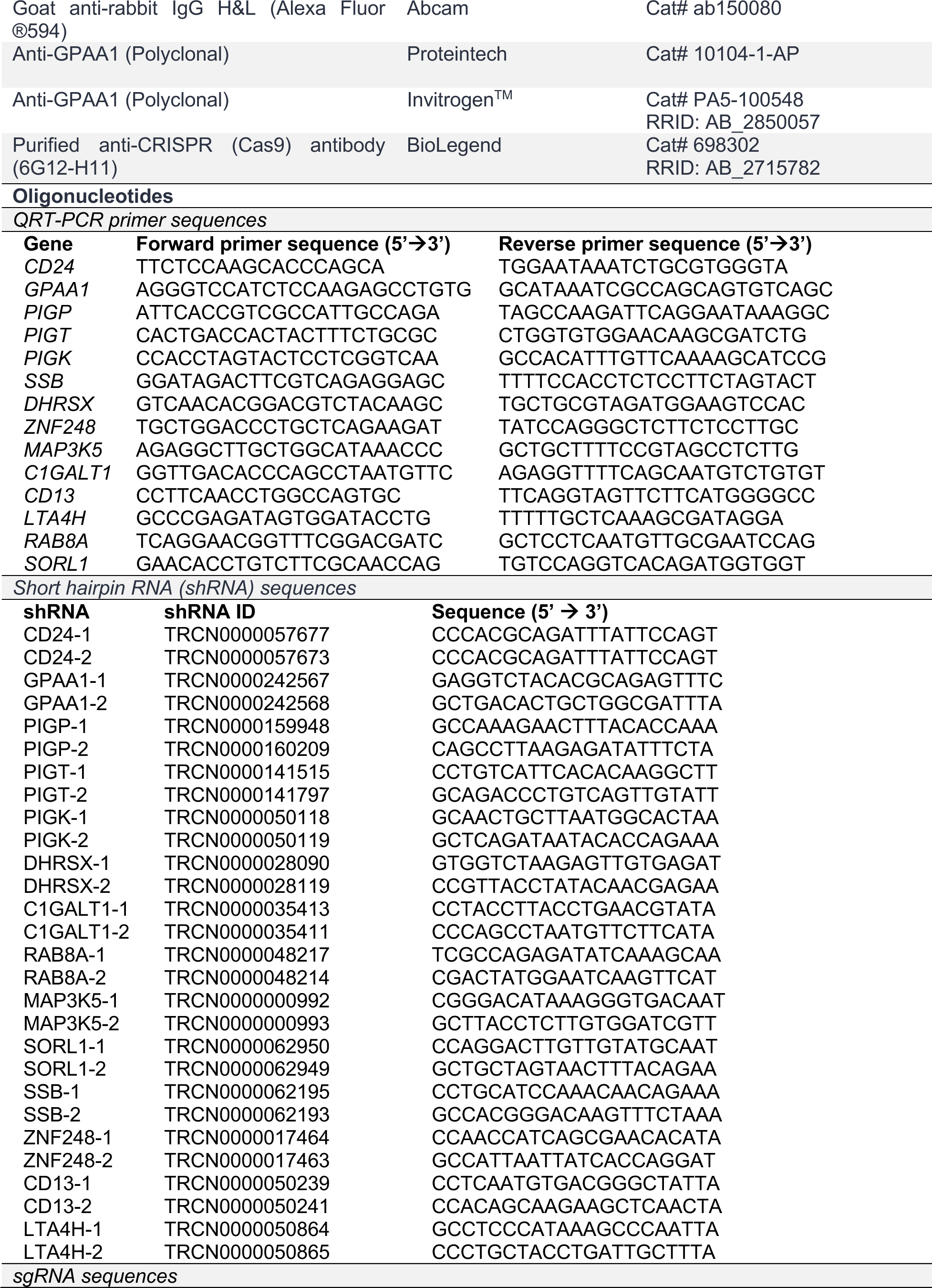
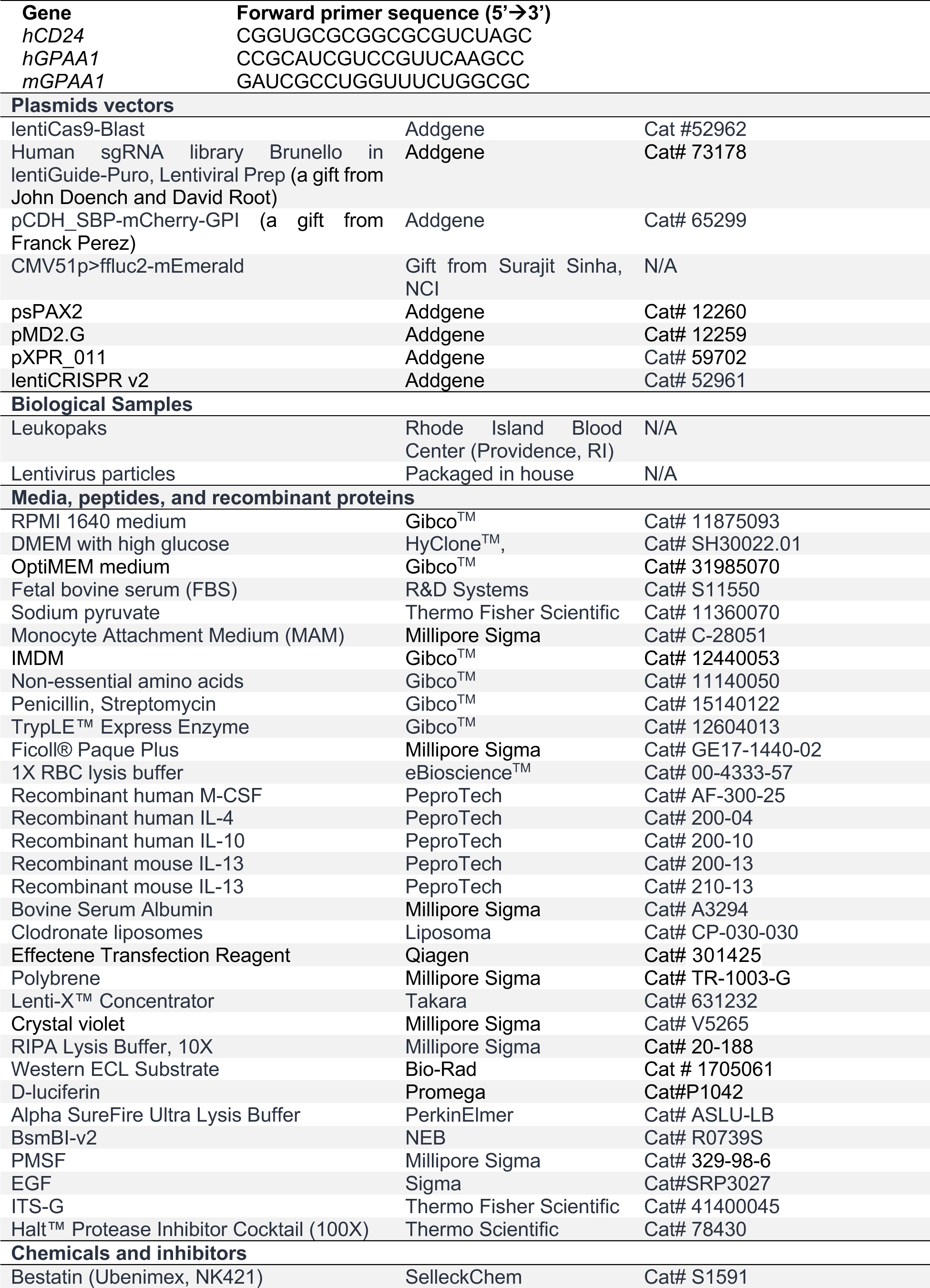
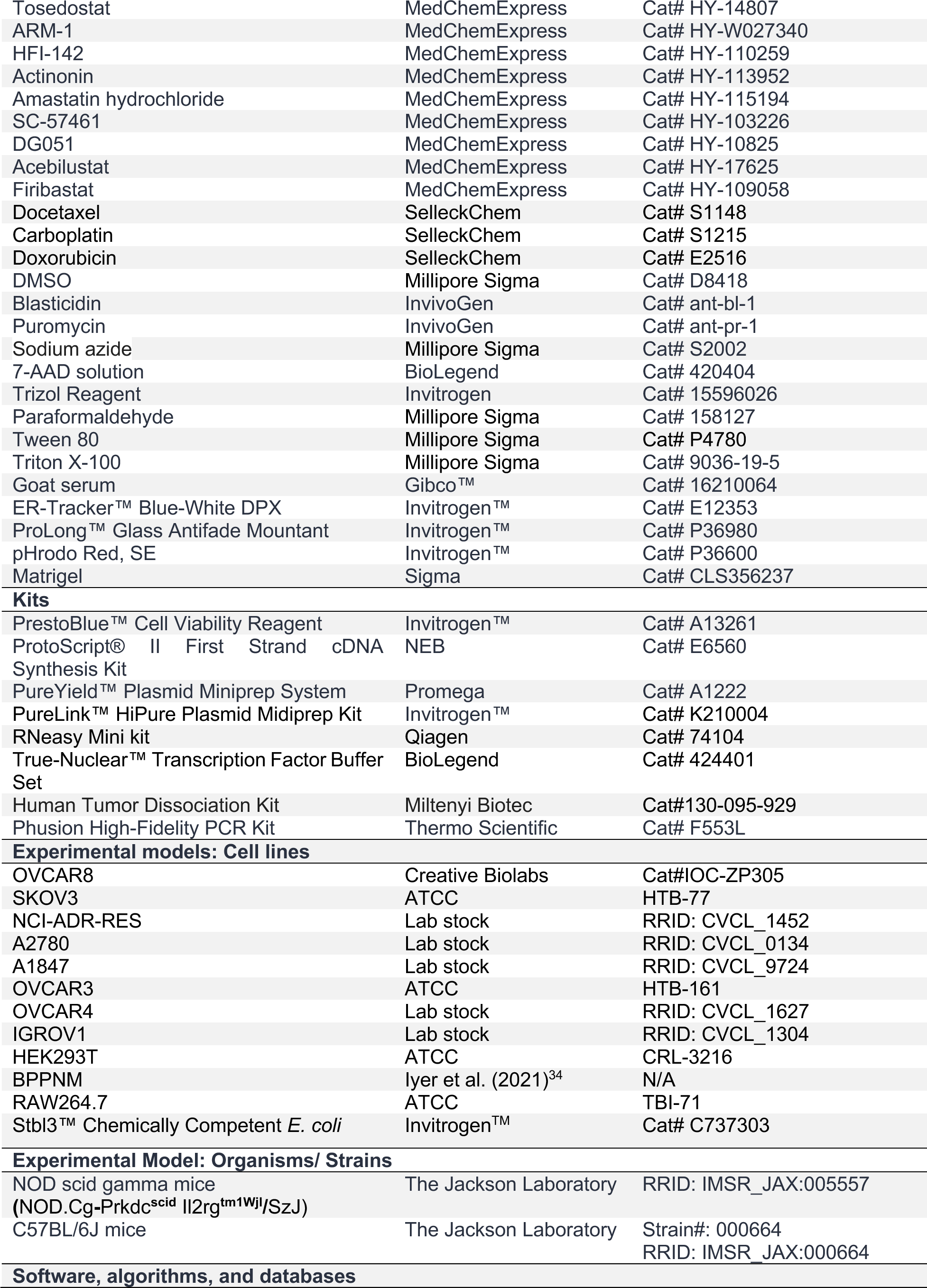
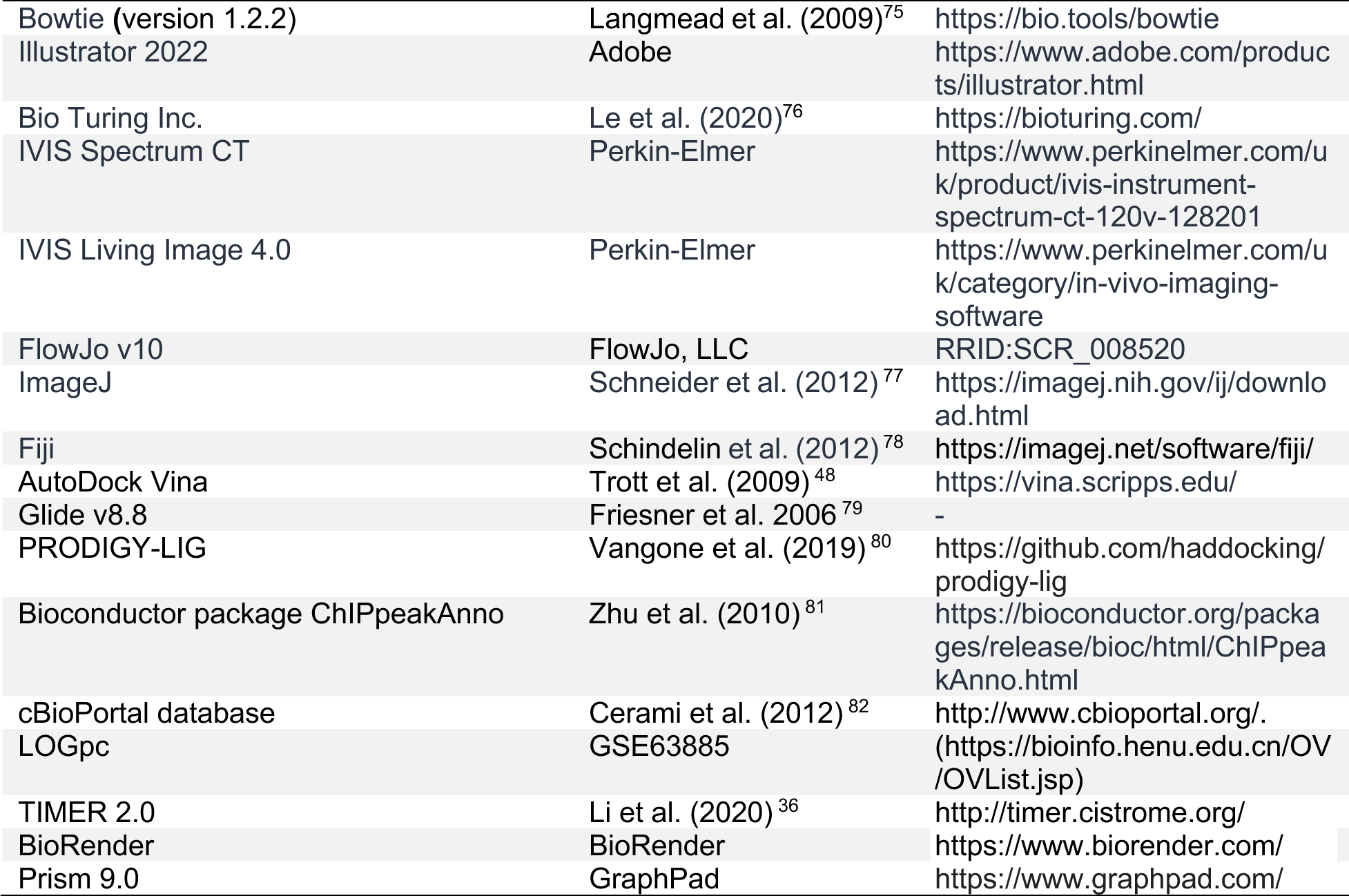

## AUTHOR CONTRIBUTIONS

Conceptualization and study design: A.K.M., S.K.M., M.R.G. Methodology: A.K.M., T.Y., S.B., R.P.T., C.T.T.S., N.N.H.P., A.K., S.R.C., T.M.S., K.S., L.J.Z., S.K.D., P.R.T; S.K.M. Investigation: A.K.M., T.Y., S.B., R.P.T., A.K., F.E., S.K.M. Formal analysis: A.K.M., F.E., S.K.M., M.R.G. Computational analysis: L.J.Z, C.T.T.S., N.N.H.P., B.E, F.E. Resources: M.R.G., F.E. Supervision: F.E, S.K.M., M.R.G. Writing-original draft: A.K.M., S.K.M., M.R.G. Writing-reviewing & editing: A.K.M., C.T.T.S., B.E, S.K.D., K.S., F.E., S.K.M. Funding acquisition: M.R.G. All authors read and approved the final manuscript.

## DECLARATION OF INTERESTS

A.K.M., S.K.M., and M.R.G. are listed as inventors on a patent application filed by the University of Massachusetts Chan Medical School on ‘Targeting Glycosylphosphatidylinositol (GPI) pathway proteins to treat Ovarian cancer’. All other authors declare no competing interests.

## INCLUSION AND DIVERSITY

We support inclusive, diverse, and equitable conduct of research.

## RESOURCE AVAILABILITY

### Lead contact

Further information and requests for resources should be directed to and will be fulfilled by the lead contact, Sunil Malonia.

---

**Figure S1.**
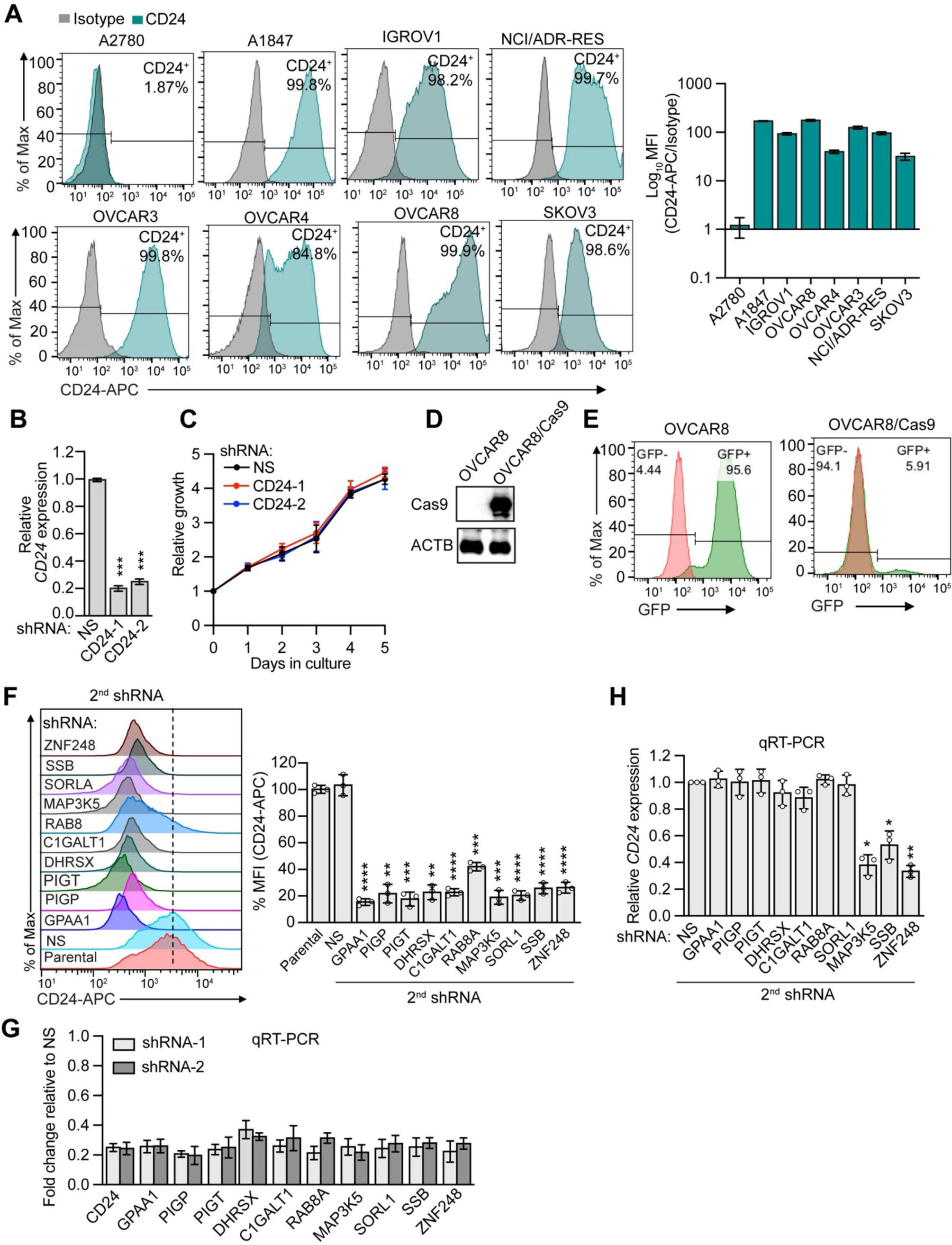
Genome-scale CRISPR screen identifies regulators of CD24. (A) (Left) Representative flow cytometry histograms of CD24 expression in a panel of ovarian cancer cell lines. The isotype is shown in gray. (Right) Quantification of MFI. (B) QRT-PCR analysis monitoring *CD24* knockdown efficiency in OVCAR8 cells expressing an NS or two independent shRNAs targeting CD24. (C) Cell proliferation assay in OVCAR8 cells expressing an NS or CD24 shRNA cultured for the indicated times and analyzed by PrestoBlue assay. (D) Immunoblot monitoring Cas9 levels in parental or OVCAR8 cells stably expressing Cas9. β-actin (ACTB) was monitored as a loading control. (E) Cas9 activity assay. Flow cytometry histograms showing GFP levels in parental or Cas9-expressing OVCAR8 cells transduced with a lentivirus-expressing GFP ORF and GFP-targeting sgRNAs. The results show that the GFP signal was diminished only in cells expressing Cas9, indicative of Cas9 activity. (F) (Left) Representative histograms of CD24 cell surface expression in OVCAR8 cells expressing second shRNA targeting candidate factors or as a control an NS shRNA. (Right) Quantification of MFI. The results were normalized to that obtained in parental OVCAR8 cells. (G) QRT-PCR analysis monitoring knockdown efficiency of two independent shRNAs against candidate factors in OVCAR8 cells. The results were compared to NS (not shown), which was set to 1. (H) QRT-PCR analysis monitoring *CD24* expression in OVCAR8 cells expressing second shRNA targeting candidate factors or an NS shRNA as a control. Data represents ± SD (n=3). P values were calculated using one-way ANOVA followed by Dunnett’s multiple comparisons test. *\*P<0.05, **P<0.01, ***P<0.001, ****P<0.0001*.

**Figure S2.**
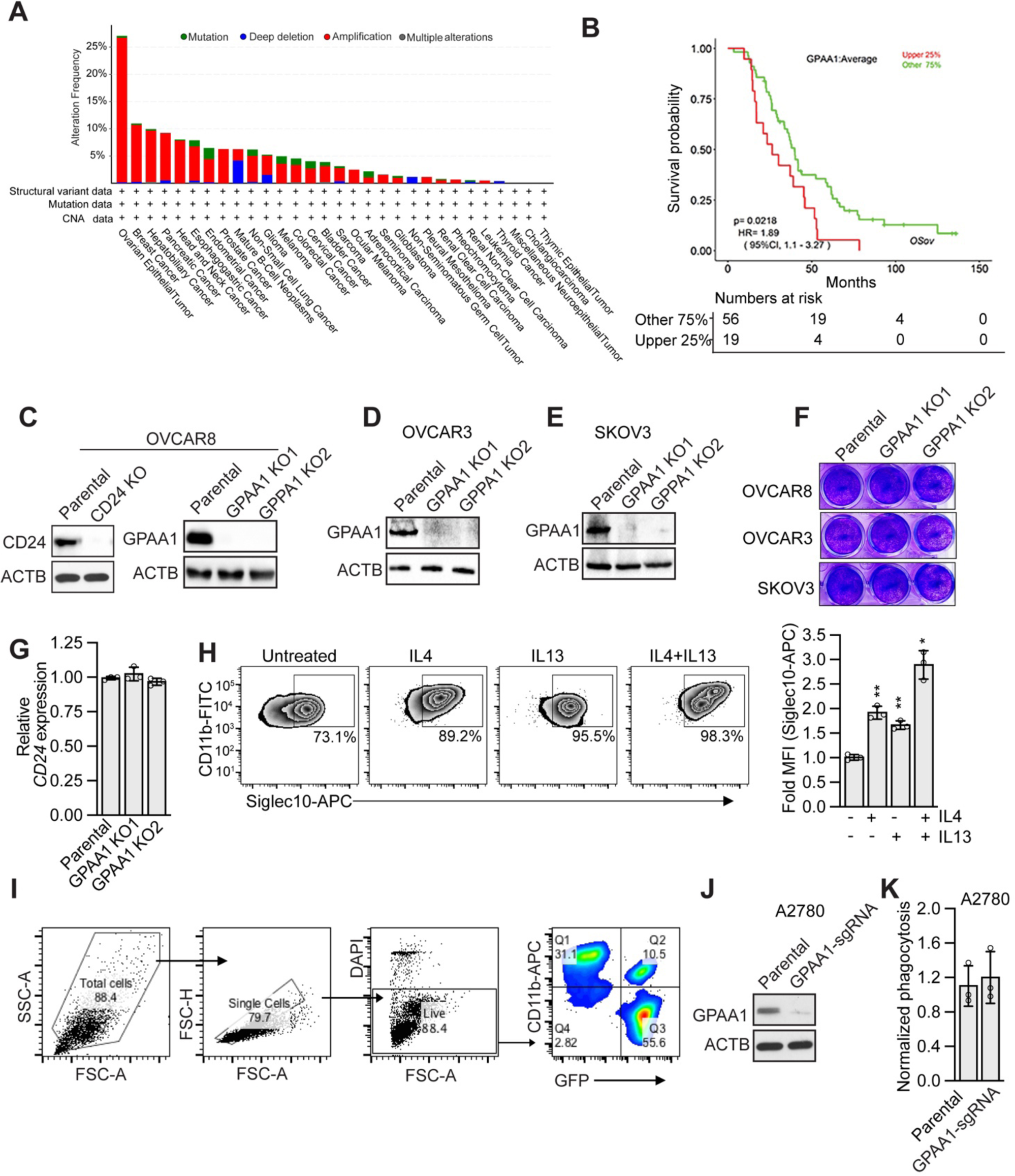
GPAA1 is amplified in ovarian cancer and its genetic depletion augments ovarian cancer cell phagocytosis. (A) Plot showing the alteration frequency of GPAA1 in multiple cancers; data were obtained through the cBioPortal web service. (B) Kaplan-Meier survival analysis showing the correlation between *GPAA1* expression and overall survival in HGSOC patients. (C) Immunoblot monitoring CD24 levels in CD24 KO (left) and GPAA1 KO (right) OVCAR8 cells. (D-E) Immunoblot monitoring GPAA1 expression in GPAA1 KO OVCAR3 (D) and SKOV3 (E) cells. (F) Cell proliferation assay using crystal violet staining in parental or GPAA1 KO OVCAR8, OVCAR3 and SKOV3 cells. (G) QRT-PCR analysis monitoring *CD24* expression in parental or GPAA1 KO OVCAR8 cells. (H) (Left) Representative flow cytometry histogram of Siglec-10 expression in human PBMC-derived macrophages stimulated with IL-4 or Il-13 or both, or untreated. (Right) Quantification of MFI. The results were normalized to those obtained in untreated macrophages. Data represents ± SD (n=3). P values were calculated using one-way ANOVA followed by Dunnett’s multiple comparisons test. (I) Gating strategy for the *in vitro* phagocytosis assay. (J) Immunoblot monitoring GPAA1 levels A2780 cells expressing GPAA1 sgRNA. (K) Quantification of *in vitro* phagocytosis in parental or GPAA1 KO A2780 cells. Data represents ± SD (n=3). P values were calculated using an unpaired student’s t-*test*. *\*P<0.05, **P<0.01*.

**Figure S3.**
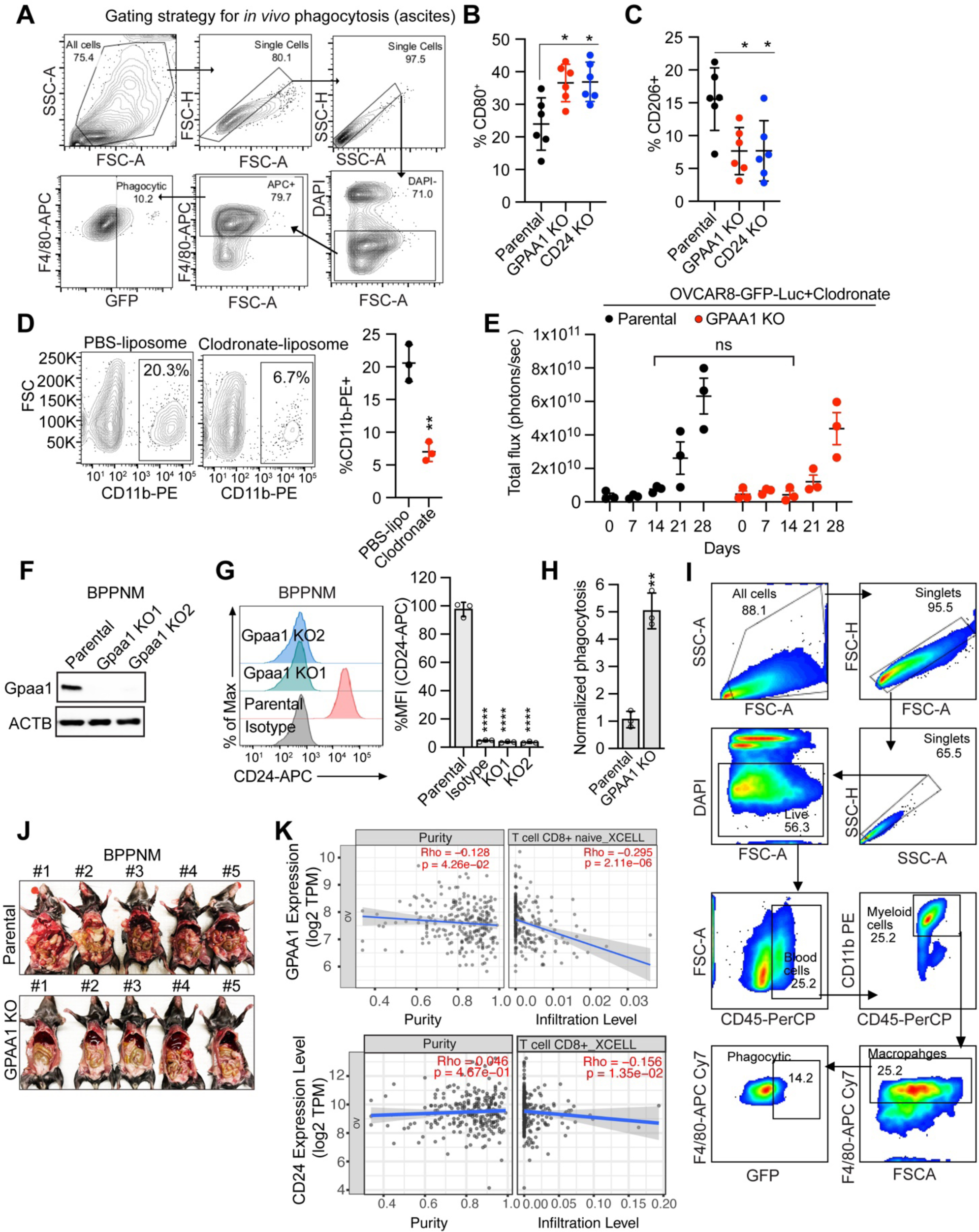
GPAA1 knockout augments *in vivo* phagocytosis and suppresses the growth of ovarian tumors in mice. (A) Gating strategy for the *in vivo* phagocytosis assay in ascites harvested from NSG mice. (B-C) Quantification of CD80+ or CD206+ macrophages by flow cytometry in ascites from mice bearing tumors derived from parental, GPAA1 KO, or CD24 KO OVCAR8 cells. Error bars indicate SD (n=6 mice per group). P values were calculated using one-way ANOVA followed by Dunnett’s multiple comparisons test. (D) (Left) Representative flow cytometry histograms monitoring macrophage depletion in ascites of mice treated with PBS- or clodronate-liposomes. (Right) Quantification of percent macrophage depletion. Error bars indicate SD (n=3 mice per group). P values were calculated using an unpaired student’s t-test. (E) Tumor growth measurement, as assessed by total flux using bioluminescence imaging, in mice implanted with parental or GPAA1 KO OVCAR8 cells and treated with clodronate-liposomes. P values were calculated using one-way ANOVA followed by Dunnett’s multiple comparisons test. (F) Immunoblot monitoring mouse Gpaa1 levels in Gpaa1 KO BPPNM cells. (G) (Left) Representative flow cytometry histograms of CD24 cell surface expression in parental and Gpaa1 KO BPPNM cells. (Right) Quantification of MFI. P values were calculated using one-way ANOVA followed by Dunnett’s multiple comparisons test. (H) Quantification of *in vitro* phagocytosis of parental or Gpaa1 KO BPPNM cells by murine RAW264.7 macrophages. (I) Gating strategy for the *in vivo* phagocytosis in BPPNM tumors in C57BL/6. (J) Images of C57BL/6 mice harboring omental tumors derived from parental or Gpaa1 KO BPPNM cells. (K) Correlation of *GPPA1* and *CD24* expression with CD8+ T cell infiltration levels in ovarian cancers. Data was obtained from the TIMER2.0 database.

**Figure S4.**
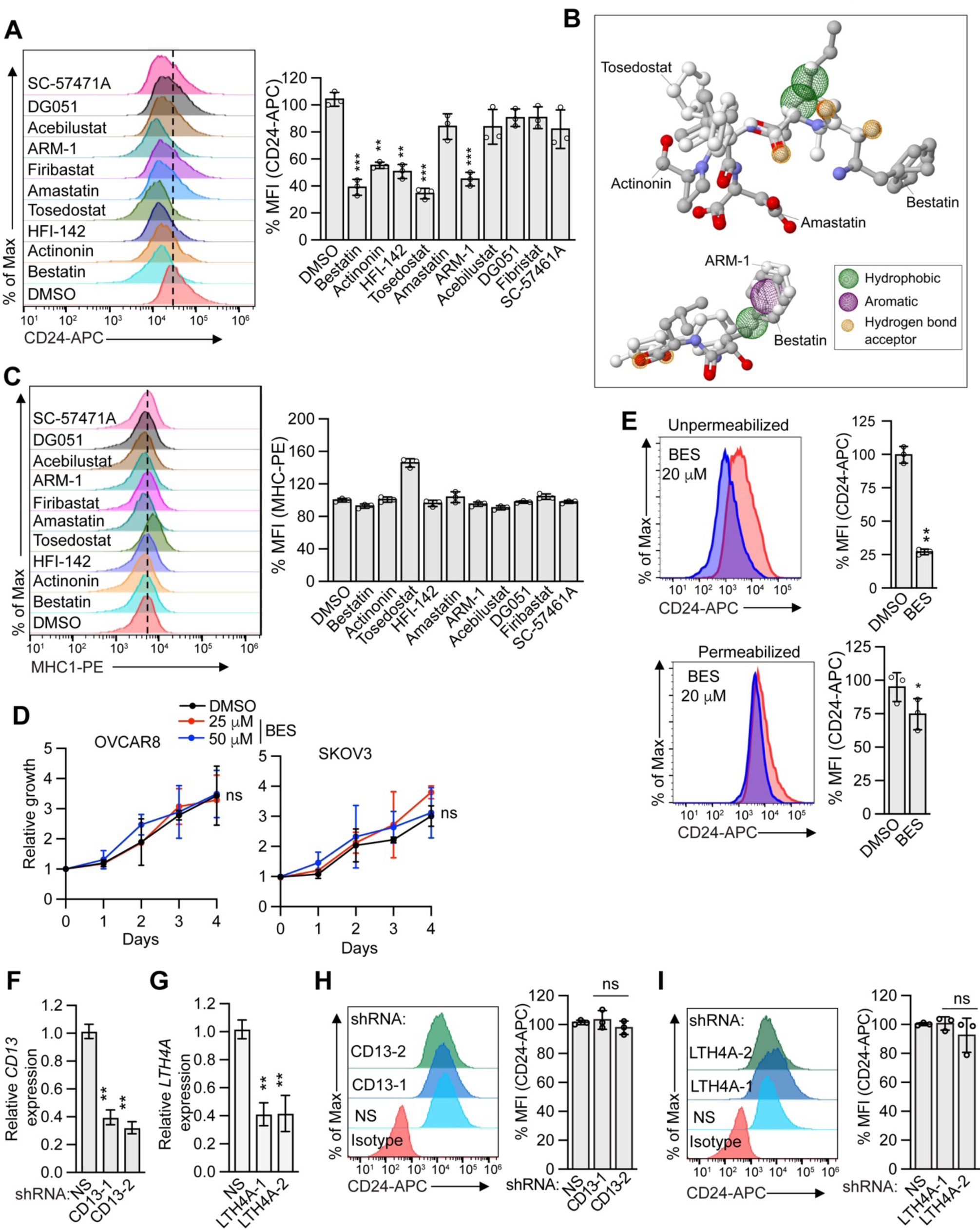
Aminopeptidase inhibitors inhibit GPAA1. (A) (Left) Representative flow cytometry histograms of CD24 cell surface expression in OVCAR8 cells treated with either DMSO or inhibitors; bestatin (20 µM), actinonin (10 µM), HFI-142(1 µM), tosedostat (5 µM), amastatin (10 µM), Firibastat (10 µM), ARM-1 (1 µM), Acebilustat (5 µM), DG051(1 µM), SC-57461A (5 µM) (Right) Quantification of MFI. (B) Supposition of structures of actinonin, amastatin, ARM-1, bestatin, and tosedostat showing similar pharmacophores such as hydrophobic and/or aromatic groups and hydrogen bond acceptors. These functional groups enable the compounds to interact with active sites of aminopeptidases via a similar interaction mechanism. The figure was generated using the Schrödinger suite. (C) (Left) Representative flow cytometry histogram showing MHC-1 cell surface expression in OVCAR8 cells treated with either DMSO or various APIs. (Right) Quantification of MFI. (D) Cell proliferation assays in OVCAR8 (left) or SKOV3 (right) cells treated with either DMSO or bestatin (20 or 50 mM), cultured for indicated times and analyzed by Prestoblue assay. (E) Representative flow cytometry analysis showing CD24 expression in OVCAR8 cells treated with DMSO or 20 µM bestatin in un-permeabilized (left) or permeabilized (right) conditions. Quantification of MFI is shown. (F-G) QRT-PCR analysis monitoring knockdown efficiency of *CD13* (left) or *LTA4H* (right) shRNAs. (H-I) (Left) Flow cytometry analysis showing CD24 cell surface expression in OVCAR8 cells expressing an NS or two unrelated shRNA targeting CD13 (H) or LTA4H (I). (Right) Quantification of MFI. Data represents ± SD (n=3). P values were calculated for (A), (C), (D), (F), (G), (H), (I) using one-way ANOVA followed multiple comparisons test, and for (E) using unpaired student’s t-test.

**Figure S5.**
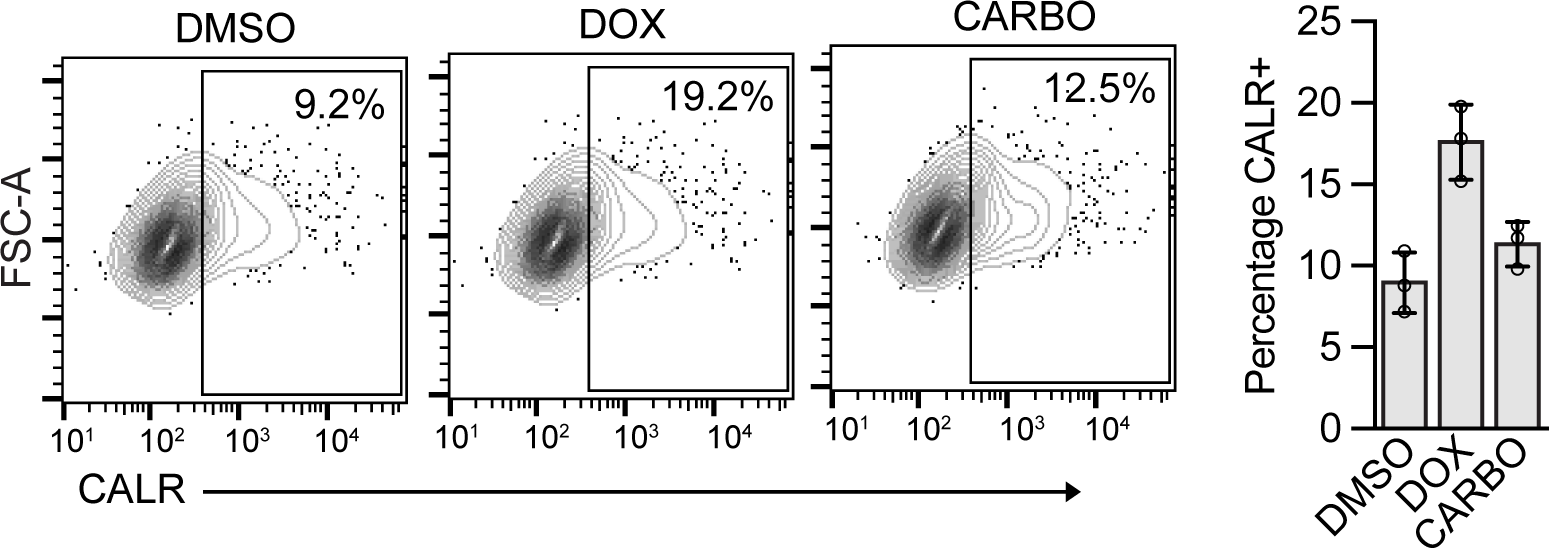
Effect of Doxorubicin and carboplatin treatment on calreticulin expression. (Left) Representative flow cytometry plots showing calreticulin (CALR) surface expression in OVCAR8 cells treated with DMSO or doxorubicin (DOX) or carboplatin (CARBO). (Right) Quantification of percent CALR-positive cells. Data represents ± SD (n=2).

