## Supplementary-file-computational analysis for "Inhibiting the GPI Transamidase Subunit GPAA1 Abolishes CD24 Surface Localization and Enhances Macrophage-Mediated Phagocytosis of Ovarian Cancer Cells"

### Computational Analysis of the GPAA1 Structure and of its Binding of Inhibiting Compounds

#### Computational Supplement (CS) for the Article

[1]Department of Molecular, Cell and Cancer Biology, University of Massachusetts Chan Medical School, Worcester, Massachusetts 01605, USA
[2]Bioinformatics Institute (BII), Agency for Science, Technology and Research (A*STAR), 30 Biopolis Street, Matrix, #07-01, Singapore 138671
[3]Faculty of Biology and Biotechnology, University of Science, Vietnam National University, 227 Nguyen Van Cu Street, District 5, Ho Chi Minh City, Vietnam
[4]Department of Molecular Medicine, and Program in Bioinformatics and Integrative Biology, UMass Chan Medical School, Massachusetts 01605, USA
[5]LASA – Lausitz Advanced Scientific Applications gGmbH, Straße der Einheit 2-24, D-02943 Weißwasser, Federal Republic of Germany
[6]Department of Biochemistry and Molecular Biotechnology, University of Massachusetts Chan Medical School, Worcester, Massachusetts 01605, USA
[7]School of Biological Sciences, Nanyang Technological University (NTU), 60 Nanyang Drive, Singapore 637551

The results of the GPAA1-inhibitor complex computational analyses are described in five sections:

I. Structural descriptors of aminopeptidase inhibitors and their binding pockets
II. Docking analyses of aminopeptidase inhibitors to the Gpaa1 active site
III. Computational analysis of known clinical mutations of Gpaa1
IV. Comparison of structural models of Gpaa1 and cryo-EM structures of the transamidase complex
V. Implication of the GPAA1 structure on the aminopeptidase-like function of GPAA1 and a proposed GPAA1 reaction mechanism

The first section provides a comprehensive analysis of aminopeptidase active sites in known 3D structures with inhibitors in the Protein Data Bank. The culminating conclusion is that Gpaa1 most likely interacts with inhibitors via the metal ion (most likely zinc) as seen in other related aminopeptidases with two hydrogen acceptor functional groups encircling the zinc ion, the dominant arrangement in known structural complexes. Notably, one zinc ion is the preferred interaction partner for inhibitor molecule even in aminopeptidases with two zincs.

The molecular mechanics/dynamics calculation results presented in the second section show that this binding mode is highly plausible in our Gpaa1 models [1] from the structural and energetic/binding energy point of view.

The third section delivers an analysis of the structural effects of clinically observed mutations, how they affect Gpaa1's stability. We find that most mutations destabilize GPAA1. The mutation L291P is a notable exception as it apparently tends to entropically stabilize the conformation of one of the two flaps surrounding the active site of Gpaa1 by reducing the backbone flexibility.

In the fourth section, we compare the published GPIT structures of the Alphafold model[2] (UnitProt: O43292), of the Baker group model[3] as well as of the two experimental cryo-EM structures in the literature[4,5]. Additionally, we analyze structural differences between our two Gpaa1 models $GPAA1^{Zn}$ and $GPAA1^{ZnZn}$ and the Gpaa1 subunit structures in the four published GPIT structures. We find that the latter three models are largely identical (with the implication how independent they are as the experimental restraints alone are insufficient to achieve a high resolution). Most importantly, all four feature a large distance between the active sites of PigK and Gpaa1 that seem to make any catalytic cooperation between the two molecules impossible.

As we see below in the fifth section, this result is in conflict with previously reported biochemical data about the function of Gpaa1 in the genesis of GPI lipid anchored proteins as well as with sequence-analytic evidence [6]. Surprisingly, this contradiction was not worth even be mentioned in any of the three GPIT structure publications or the starting point of a critical rethink. This conflict seems even more severe in the light of the experimental provided in this work that show specific, aminopeptidase-like inhibition of Gpaa1 can disrupt the native GPIT complex' activity.

Thinkable conflict exit options out of the contradiction are (1) GPIT acting as a multimer with PigK and Gpaa1 from two different complexes working together (following the thought developed by the Gamage, Hendrickson, et al.; see references[7-9]) or (2) the published complex structures are partly artifacts and are not biologically relevant in some aspects that concern the catalytic process. At the same time, our Gpaa1 models are quite similar to the Gpaa1 structures in all three published GPIT complexes except for some details regarding the active site arrangements and the inclusion of zinc ions there. To note, our models are in principal arrangement with all accumulated data from cell and molecular biology regarding GPIT, both from previous literature as well as from experiments in this work as well as with sequence-analytically based homology consideration derived from other aminopeptidases [6].

#### I. Structural descriptors of aminopeptidase inhibitors and their binding pockets

The GPAA1 structural fold resembles that of the M28 metallo-protease family including aminopeptidases [6]. In the present article, GPAA1 is shown to be inhibited by a number of aminopeptidase inhibitors such as bestatin. Therefore, we investigated the nature of the GPAA1 enzymatic activity by exploring physico-chemical and structural descriptors to reveal binding properties of the aminopeptidase active sites including that of GPAA1. Additional methodical details are explained in the Methods Appendix of this document.

Among the downloadable (118/121) complex structures from the Protein Data Bank (PDB), 107 aminopeptidase complexes contain small molecule-type inhibitors. The remaining set of the aminopeptidase inhibitors are peptide-like types (the full list in Supplementary File CSE1). The latter PDB structures were excluded from further analysis in this computational study. The majority of the remaining structures (82/107) include metals in the inhibitor-binding regions, with 89% (73/82) involving direct metal binding. On the other hand, no information of metals was captured in the other (25/107) complexes.

With these variations in mind, we first computed various characteristics of ion and inhibitor binding by aminopeptidases such as numbers of metals in the aminopeptidase active sites and the amino acid residues directly interacting with the ions and the corresponding inhibitors. Subsequently, structural descriptors of the inhibitor-bound pocket properties such as amino acid propensity, pocket/ligand volumes, hydrophobicity scores, charge scores, as well as the binding energies were assessed using the tool *dpocket* [10]. Noticeably, we found only a weak correlation ($R_{Pearson}$ = 0.65, $p < 10^{-4}$) between the volumes of the binding pockets and of the ligands, suggesting sometimes only fragments of the inhibitors are involved in the pocket binding.

Then, the inhibitor pharmacophores (e.g. aromatic, hydrophobic, hydrogen bond donor, and/or acceptor groups, identified using the PharmaGist server [11]), were used together with the detected feature of pocket residue propensity in the subsequent grouping of aminopeptidase-inhibitor complexes. The set of 107 inhibitors was clustered using *k-means* clustering incorporated with principal component (PC) analysis (Supplementary Figure CSF1) implemented in the sklearn v.0.23.2 package [12].

**Supplementary Figure CSF1 - *Clusters of the 107 aminopeptidase-inhibitor-complexes***

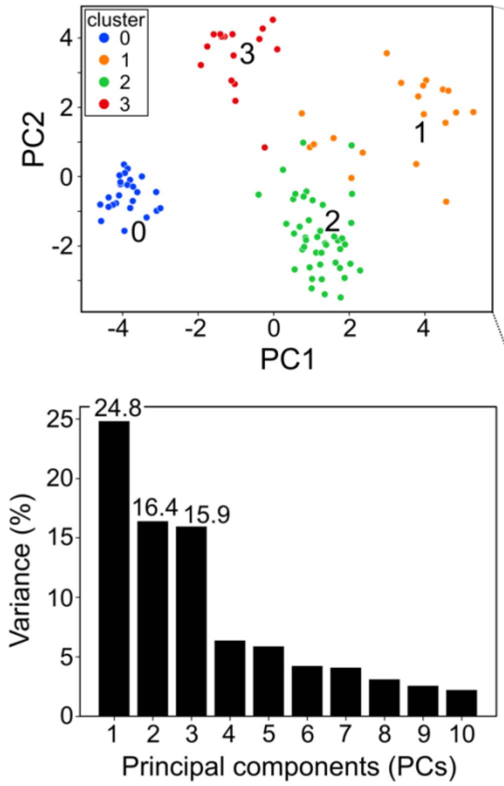


| | | # metal binding to LIG |
|---|---|---|
| cluster 0 | 0a | 0 |
| | 0b | no metal |
| cluster 1 | 1a | 0 |
| | 1b | no metal |
| | 1c | 1 to 2 (ZNs) |
| cluster 2 | 2a | 1 (ZN) |
| | 2b* | 2 (ZNs) |
| cluster 3 | 3a | 0 |
| | 3b | 1 (CO) |
| | 3c | 1 to 2 (ZNs) |
| | 3d | 2 (MNs) |
| | 3e | 2 to 3 (NIs) |

*a singleton, different from the others in cluster 2

*Clusters of the 107 aminopeptidase inhibitors using a combination of PCA and k-means clustering. For cluster visualization, the inhibitor dataset was projected on the first two principal components PC1 and PC2. The explained variances of the first 10 PCs with the variance ratio (%) of the first 3 PCs were labelled. Details of the metal binding capacity of each sub-cluster were tabulated as also shown in Supplementary Table CST1.*

Low variances were observed. We found that ~80% of the total variances explained by the first 6 PCs with the variances maximized within the first 3 PCs. The latter were then used for the *k-means* clustering as labelled in Supplementary Figure CSF1. Notably, the same clustering procedure using the first 6 PCs resulted ~99.1% similar membership (106/107 members remained in the same clusters, data not shown). Four clusters ($k$ = 4) were finalized as the best number of clusters (see clustering evaluation in Supplementary Figure CSF2). The 4 clusters were further sub-grouped according to the metal binding capacity of the aminopeptidase active sites shown in Supplementary Table CST1, e.g., lacking metal (*no_metal*), with direct metal binding (*#num_metal* > 0), without direct metal binding (*#num_metal* = 0), or different types of metals (zinc (ZN), cobalt (CO), manganese (MN), or nickel (NI), etc.).

**Supplementary Figure CSF2 - *Evaluation of number of clusters for the clustering.***

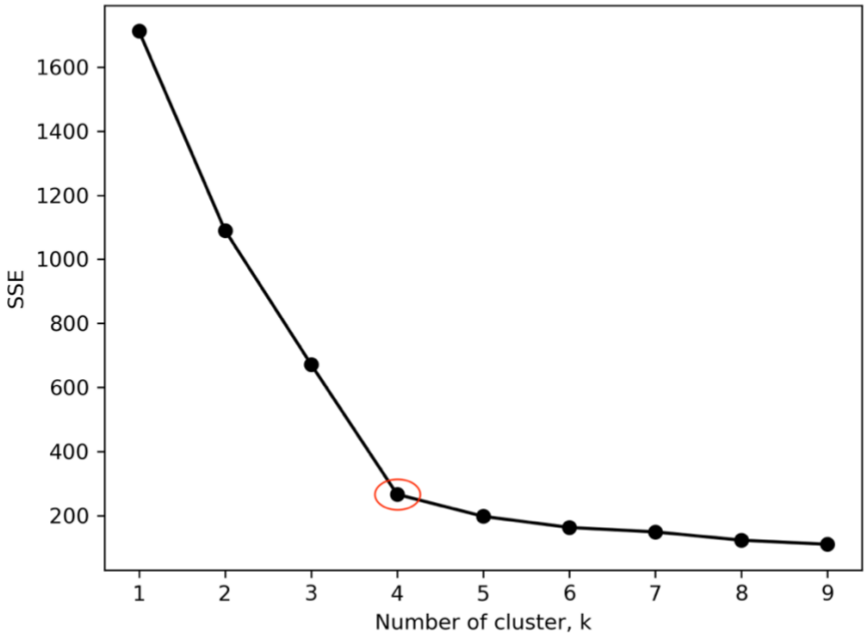


*The number k = 4 shows the optimal number of clusters with the optimal decrease (elbow point, circled in red) of inertia values (SSE). The analysis was performed using kmeans clustering in the sklearn v.0.23.2 package.*

**Supplementary Table CST1 - Clusters defined by metal binding capacity of the aminopeptidase active sites**

| Cluster | Sub-cluster | Number of metals binding to ligand |
|---|---|---|
| Cluster 0 | 0a | 0 |
| | 0b | No metal |
| Cluster 1 | 1a | 0 |
| | 1b | No metal |
| | 1c | 1 to 2 (ZNs) |
| Cluster 2 | 2a | 1 (ZN) |

| | | |
|---|---|---|
| | 2b* | 2 (ZNs) |
| Cluster 3 | 3a | 0 |
| | 3b | 1 (CO) |
| | 3c | 1 - 2 (ZNs) |
| | 3d | 2 (MNs) |
| | 3e | 2 - 3 (NIs) |

*a singleton, different from the others in cluster 2

*Clusters defined by metal binding capacity of the aminopeptidase active sites extracted from the retrieved set of 107 aminopeptidase complexes. The metals include zinc (ZN), cobalt (CO), manganese (MN), or nickel (NI).*

Among the 28 features used for the clustering (Supplementary File CSE2), the residues' physicochemical properties within the aminopeptidase binding pockets appeared critical for determining the distinct clusters of the respective inhibitor-aminopeptidase complexes. For example, numbers of tyrosine (Y), alanine (A), methionine (M) contributed the most in maximizing the group variances along the PC1, etc. (Supplementary Table CST2).

**Supplementary Table CST2 – top features for principal component analysis of binding pockets**

| | Top 3 important features of the inhibitors | Weight* of the features (The higher, the more important on the corresponding PCs) |
|---|---|---|
| PC1 | Number of tyrosine (Y) in the binding pocket | 0.342 |
| | Number of alanine (A) in the binding pocket | 0.331 |
| | Number of methionine (M) in the binding pocket | 0.325 |
| PC2 | Number of valine (V) in the binding pocket | 0.399 |
| | Number of glutamate (E) in the binding pocket | 0.362 |
| | Number of cysteine (C) in the binding pocket | 0.315 |

| | | |
|---|---|---|
| PC3 | Number of serine (S) in the binding pocket | 0.378 |
| | Number of isoleucine (I) in the binding pocket | 0.296 |
| | Size of inhibitors (#num atoms) | 0.293 |

*Absolute values of the components in the eigenvectors

*The top 3 important features contributing to maximizing the variances along the first three principal components PC1, PC2, and PC3. The full analysis can be found in Supplementary File CSE2.*

It was also noticed that metal bindings were present in various complexes across the clusters (Supplementary Figure CSF1, right), highlighting the potential key impact of the metal interactions in the aminopeptidase activity. Indeed, the derived structural features eliciting the differing inhibitor-binding pocket properties, particularly in the presence or absence of metal ions, e.g., $Zn^{2+}$, $Co^{2+}$, $Mn^{2+}$, or $Ni^{2+}$.

Common pharmacophores of each sub-cluster were illustrated relative to their respective pivot molecule (Supplementary Figure CSF3). E.g., the inhibitor labelled “677” in the aminopeptidase-inhibitor complex PDB: 3hab represents the sub-cluster 0a. Differently coloured presentations of the pharmacophore groups, for example by using the ZINC-Pharmer [13], reveal similar arrangements of the pharmacophores across the sub-clusters. To note, all highlighted pharmacophore groups were found to be involved in direct interactions between the inhibitors and the respective aminopeptidase pockets.

**Supplementary Figure CSF3 - *Common pharmacophores of the retrieved set of 107 aminopeptidase inhibitors***

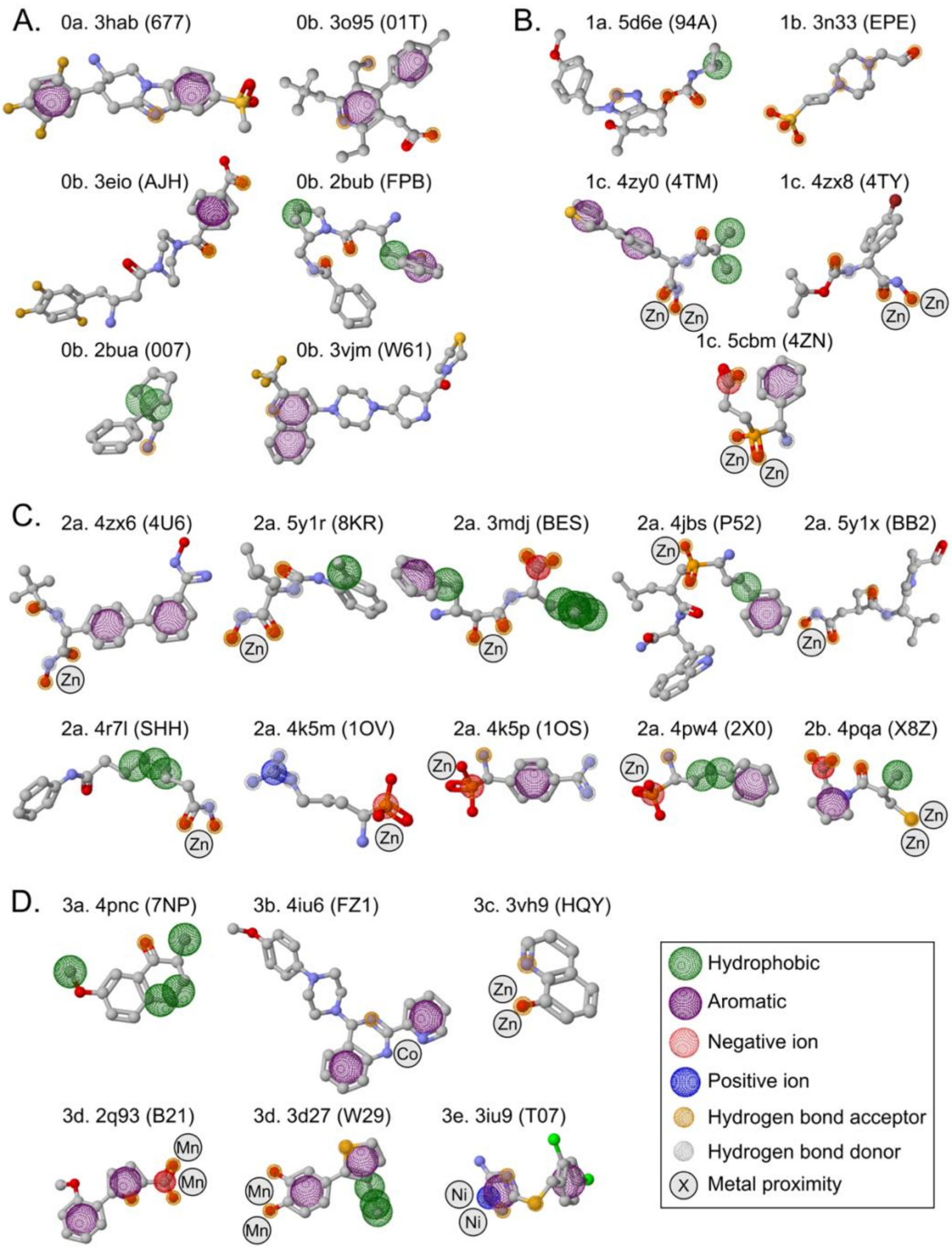


*The common pharmacophores presented on the pivot molecule (with attached PDB id and the respective inhibitor labelled and shown in sticks) of each sub-cluster. The sub-cluster labels are tabulated in Supplementary Table CST1.*

A comparable pattern of oxygen atom pairs (~2.6 Å apart) acting as hydrogen bond acceptors is present in various sub-clusters of the inhibitors, particularly in those binding to zinc, e.g., sub-clusters 1c and 2a (Supplementary Figure CSF3). Interestingly, one of the oxygens could predominantly share the binding with two zinc ions. These hydrogen bond acceptors arrangement appeared to be crucial for the inhibitors to bind the ZN metal in the aminopeptidase pockets. Besides, hydrophobic and/or aromatic groups were found predominant in those inhibitors, and with an additional negative charged group detected near the Zn-binding oxygen pair (sub-cluster 2a, PDBs: 4k5m, 4k5p, and 4pw4).

However, in the of sub-clusters 2b and 3c (Supplementary Figure CSF3, section C and D), the inhibitors also bind two zinc ions but in a different manner. In the sub-cluster 2b (PDB: 4pqa, with the L-captopril inhibitor X8Z), the two zincs were bridged by the sulfhydryl group [14] instead, whereas in the other (PDB: 3vh9, with an 8-quinolinol derivative as an inhibitor, HQY), the zinc ions bound in a bidentate manner to both the inhibitor nitrogen and oxygen atoms [15].

On the other hand, there were no consistent pharmacophore patterns among the sub-clusters of inhibitors binding to aminopeptidases that lack metals (sub-clusters 0b and 1b) or contain no direct metal and inhibitor interactions (sub-clusters 0a, 1a, and 3a).

The metal ions appear to drive the binding modes of the inhibitors and the corresponding formation of the aminopeptidase pockets (see Supplementary Tables CST3 and CST4 further down). For example, presence of both negatively charged residues glutamic acid (E) and aspartic acid (D) was detected in the metal-containing pockets of clusters 2 and 3, or with the more noticeably predominant D (compared with E) in sub-cluster 1c. In those cases, hydrophobic residues alanine (A), valine (V) and/or leucine (L) are only present in Zn(s)-binding pockets. On the other hand, cysteine (C) was only present in the pockets that bound the other metal ions $Co^{2+}$, $Mn^{2+}$, and $Ni^{2+}$ (except for a single case in sub-cluster 3c, PDB: 3vh9 binding two zincs).

**Supplementary Table CST3 -** ***Properties of the aminopeptidase pockets that bound to the inhibitors present in the clusters 0-1 (A) and cluster 3 (B).***

A.

| | | | | Hydrophobic | | | | | | | | Hydrophilic | | | | | | | Positive | | | Negative | |
|---|---|---|---|---|---|---|---|---|---|---|---|---|---|---|---|---|---|---|---|---|---|---|---|
| pdb | LIG | LIG_clusters | sub_cluster | A | V | L | I | P | M | F | W | G | S | T | C | N | Q | Y | K | R | H | D | E |
| 3hac | 361 | 0 | 0a | 0 | 3 | 0 | 0 | 1 | 0 | 1 | 1 | 0 | 2 | 0 | 0 | 1 | 0 | 4 | 0 | 3 | 2 | 0 | 2 |
| 3hab | 677 | | | 0 | 3 | 0 | 0 | 1 | 0 | 1 | 1 | 0 | 2 | 0 | 0 | 1 | 0 | 5 | 0 | 4 | 1 | 0 | 2 |
| 4pnz | 2VH | | | 0 | 3 | 0 | 1 | 1 | 0 | 1 | 1 | 0 | 3 | 0 | 0 | 1 | 0 | 5 | 0 | 4 | 2 | 0 | 3 |
| 3o95 | 01T | | 0b | 0 | 3 | 0 | 0 | 0 | 0 | 1 | 2 | 1 | 2 | 0 | 0 | 1 | 0 | 4 | 0 | 3 | 2 | 0 | 2 |
| 3sx4 | KXA | | | 0 | 3 | 0 | 0 | 0 | 0 | 1 | 3 | 0 | 3 | 0 | 0 | 1 | 1 | 4 | 1 | 2 | 2 | 1 | 2 |
| 3opm | LUI | | | 0 | 4 | 0 | 0 | 0 | 0 | 1 | 3 | 1 | 2 | 0 | 0 | 1 | 0 | 4 | 1 | 2 | 2 | 1 | 2 |
| 3sww | KXB | | | 0 | 2 | 0 | 0 | 0 | 0 | 1 | 2 | 1 | 2 | 0 | 0 | 1 | 0 | 4 | 0 | 3 | 2 | 0 | 2 |
| 3q0t | LGE | | | 0 | 2 | 0 | 0 | 0 | 0 | 1 | 2 | 0 | 3 | 0 | 0 | 1 | 1 | 4 | 1 | 3 | 2 | 0 | 2 |
| 4g1f | 0WG | | | 0 | 2 | 0 | 0 | 0 | 0 | 1 | 1 | 0 | 2 | 0 | 1 | 1 | 0 | 5 | 1 | 2 | 1 | 0 | 2 |
| 3o9v | 10T | | | 0 | 3 | 0 | 0 | 0 | 0 | 1 | 1 | 1 | 2 | 0 | 0 | 1 | 0 | 4 | 0 | 3 | 2 | 0 | 2 |
| 3eio | AJH | | | 0 | 1 | 0 | 1 | 1 | 0 | 1 | 1 | 0 | 3 | 0 | 1 | 1 | 0 | 5 | 0 | 4 | 2 | 0 | 2 |
| 3g0c | RUF | | | 0 | 3 | 0 | 0 | 0 | 0 | 1 | 2 | 0 | 2 | 0 | 0 | 1 | 0 | 4 | 1 | 1 | 2 | 0 | 2 |
| 3g0b | T22 | | | 0 | 3 | 0 | 0 | 0 | 0 | 1 | 2 | 0 | 2 | 0 | 0 | 0 | 0 | 4 | 1 | 2 | 2 | 0 | 2 |
| 3g0d | XIH | | | 0 | 3 | 0 | 0 | 0 | 0 | 1 | 2 | 0 | 2 | 0 | 0 | 1 | 0 | 4 | 1 | 2 | 2 | 0 | 2 |
| 3vjk | M51 | | | 0 | 3 | 0 | 0 | 0 | 0 | 2 | 1 | 0 | 2 | 0 | 0 | 1 | 0 | 4 | 0 | 4 | 0 | 0 | 2 |
| 2qjr | PZF | | | 0 | 3 | 0 | 0 | 2 | 0 | 1 | 1 | 0 | 4 | 0 | 1 | 1 | 0 | 5 | 0 | 3 | 1 | 0 | 3 |
| 3vjl | W94 | | | 0 | 2 | 0 | 1 | 1 | 0 | 1 | 1 | 0 | 3 | 0 | 1 | 1 | 1 | 6 | 0 | 4 | 1 | 0 | 2 |
| 2bub | FPB | | | 0 | 2 | 0 | 0 | 0 | 0 | 1 | 2 | 0 | 2 | 0 | 0 | 1 | 0 | 4 | 0 | 3 | 1 | 0 | 2 |
| 2buc | 008 | | | 0 | 3 | 0 | 0 | 0 | 0 | 1 | 1 | 0 | 3 | 0 | 1 | 1 | 0 | 5 | 0 | 4 | 2 | 0 | 2 |
| 3h0c | PS4 | | | 0 | 3 | 0 | 0 | 1 | 0 | 1 | 1 | 0 | 3 | 0 | 1 | 1 | 1 | 5 | 1 | 5 | 0 | 0 | 2 |
| 2bua | 007 | | | 0 | 2 | 0 | 0 | 0 | 0 | 1 | 1 | 0 | 2 | 0 | 0 | 1 | 0 | 4 | 0 | 2 | 1 | 0 | 2 |
| 3wqh | SKK | | | 0 | 3 | 0 | 1 | 0 | 0 | 1 | 2 | 0 | 2 | 0 | 0 | 1 | 1 | 6 | 0 | 5 | 2 | 0 | 2 |
| 3w2t | LF7 | | | 0 | 1 | 0 | 0 | 1 | 0 | 1 | 1 | 0 | 2 | 0 | 0 | 1 | 0 | 4 | 0 | 3 | 2 | 0 | 2 |
| 3vjm | W61 | | | 0 | 2 | 0 | 1 | 1 | 0 | 1 | 1 | 0 | 3 | 0 | 1 | 1 | 1 | 6 | 0 | 5 | 1 | 0 | 3 |
| 2rgu | 356 | | | 0 | 2 | 0 | 0 | 0 | 0 | 1 | 3 | 2 | 2 | 0 | 0 | 1 | 0 | 5 | 1 | 3 | 1 | 1 | 2 |
| 3g0g | RUM | | | 0 | 3 | 0 | 0 | 0 | 0 | 1 | 2 | 0 | 2 | 0 | 0 | 1 | 0 | 4 | 1 | 2 | 2 | 0 | 2 |
| 5cls | 52T | 1 | 1a | 2 | 0 | 1 | 1 | 1 | 1 | 1 | 0 | 1 | 0 | 0 | 0 | 2 | 0 | 2 | 0 | 0 | 4 | 2 | 0 |
| 5d6f | 57R | | | 2 | 1 | 2 | 1 | 0 | 1 | 2 | 0 | 0 | 0 | 0 | 0 | 2 | 0 | 2 | 0 | 0 | 4 | 2 | 1 |
| 5d6e | 94A | | | 2 | 1 | 2 | 1 | 1 | 1 | 2 | 0 | 1 | 0 | 1 | 0 | 2 | 0 | 2 | 0 | 0 | 4 | 3 | 1 |
| 1boa | FUG | | | 2 | 1 | 2 | 1 | 0 | 1 | 2 | 0 | 1 | 0 | 1 | 0 | 2 | 0 | 1 | 0 | 0 | 4 | 3 | 1 |
| 3n33 | EPE | | 1b | 1 | 0 | 3 | 1 | 1 | 0 | 1 | 0 | 2 | 1 | 0 | 0 | 0 | 1 | 0 | 0 | 0 | 0 | 1 | 1 |
| 6ee2 | J1V | | 1c | 2 | 1 | 3 | 1 | 0 | 2 | 2 | 0 | 3 | 3 | 2 | 0 | 1 | 0 | 1 | 2 | 1 | 0 | 3 | 0 |
| 6eee | J4V | | | 3 | 0 | 3 | 1 | 0 | 2 | 2 | 0 | 3 | 3 | 2 | 0 | 2 | 0 | 1 | 2 | 1 | 0 | 3 | 0 |
| 4zy0 | 4TM | | | 3 | 0 | 3 | 1 | 0 | 2 | 2 | 0 | 3 | 2 | 2 | 0 | 1 | 0 | 0 | 1 | 1 | 0 | 3 | 0 |
| 4zy1 | 4U5 | | | 2 | 0 | 3 | 1 | 0 | 2 | 2 | 1 | 4 | 2 | 2 | 0 | 1 | 0 | 0 | 1 | 1 | 0 | 3 | 0 |
| 4zy2 | 4TL | | | 2 | 0 | 3 | 1 | 0 | 2 | 2 | 0 | 3 | 3 | 2 | 0 | 1 | 0 | 1 | 1 | 1 | 0 | 3 | 0 |
| 4zyq | 4U6 | | | 3 | 0 | 3 | 1 | 0 | 2 | 2 | 0 | 3 | 2 | 2 | 0 | 1 | 0 | 1 | 1 | 1 | 0 | 3 | 0 |
| 4zx8 | 4TY | | | 2 | 1 | 2 | 1 | 0 | 2 | 1 | 0 | 3 | 3 | 2 | 0 | 1 | 0 | 0 | 1 | 1 | 0 | 3 | 0 |
| 4zx9 | 4TK | | | 3 | 0 | 3 | 1 | 0 | 2 | 2 | 0 | 3 | 2 | 2 | 0 | 1 | 0 | 1 | 1 | 1 | 0 | 3 | 0 |
| 4zla | BES | | | 3 | 1 | 1 | 1 | 0 | 1 | 0 | 0 | 3 | 2 | 4 | 0 | 1 | 0 | 2 | 1 | 1 | 0 | 3 | 1 |
| 5ib9 | BES | | | 1 | 1 | 2 | 1 | 1 | 1 | 0 | 0 | 2 | 1 | 1 | 0 | 1 | 0 | 2 | 1 | 1 | 0 | 1 | 4 |
| 3t8w | DGZ | | | 3 | 1 | 3 | 1 | 1 | 2 | 2 | 0 | 3 | 3 | 2 | 0 | 1 | 0 | 1 | 1 | 1 | 0 | 2 | 0 |
| 5cbm | 4ZN | | | 2 | 0 | 2 | 0 | 0 | 2 | 2 | 0 | 3 | 2 | 2 | 0 | 1 | 0 | 1 | 1 | 1 | 0 | 3 | 1 |
| 4k3n | 1OT | | | 2 | 0 | 2 | 0 | 0 | 2 | 1 | 0 | 3 | 2 | 2 | 0 | 0 | 0 | 1 | 2 | 1 | 0 | 3 | 0 |
| 4x2t | TOD | | | 4 | 0 | 3 | 1 | 0 | 2 | 1 | 0 | 3 | 2 | 2 | 0 | 1 | 0 | 1 | 2 | 1 | 0 | 3 | 0 |

B.

| | | | | Hydrophobic | | | | | | | | Hydrophilic | | | | | | | Positive | | | Negative | |
|---|---|---|---|---|---|---|---|---|---|---|---|---|---|---|---|---|---|---|---|---|---|---|---|
| pdb | LIG | LIG_clusters | sub_cluster | A | V | L | I | P | M | F | W | G | S | T | C | N | Q | Y | K | R | H | D | E |
| 4z7m | 4L9 | 3 | 3a | 0 | 0 | 0 | 0 | 0 | 1 | 1 | 1 | 2 | 0 | 0 | 4 | 0 | 2 | 3 | 0 | 0 | 4 | 2 | 3 |
| 4pnc | 7NP | | | 0 | 0 | 0 | 0 | 0 | 0 | 1 | 1 | 1 | 0 | 1 | 3 | 0 | 0 | 3 | 0 | 0 | 4 | 2 | 1 |
| 4iu6 | FZ1 | | 3b | 0 | 0 | 0 | 0 | 1 | 1 | 2 | 1 | 1 | 1 | 2 | 2 | 1 | 1 | 3 | 1 | 0 | 3 | 2 | 2 |
| 2nq7 | HM5 | | | 0 | 0 | 0 | 0 | 1 | 1 | 2 | 1 | 1 | 1 | 1 | 3 | 1 | 0 | 3 | 0 | 0 | 3 | 2 | 2 |
| 1yvm | TMG | | | 0 | 0 | 0 | 0 | 0 | 1 | 1 | 1 | 1 | 0 | 1 | 3 | 0 | 0 | 3 | 0 | 0 | 4 | 2 | 2 |
| 3ked | DAB | | 3c | 1 | 1 | 0 | 0 | 0 | 1 | 0 | 0 | 1 | 0 | 0 | 0 | 0 | 0 | 3 | 0 | 0 | 1 | 0 | 1 |
| 3vh9 | HQY | | | 0 | 0 | 0 | 1 | 0 | 2 | 2 | 0 | 0 | 0 | 0 | 2 | 0 | 0 | 2 | 0 | 0 | 1 | 1 | 1 |
| 2q93 | B21 | | 3d | 0 | 0 | 0 | 0 | 0 | 1 | 1 | 1 | 0 | 0 | 1 | 4 | 0 | 1 | 3 | 0 | 0 | 4 | 2 | 1 |
| 2q94 | A04 | | | 0 | 0 | 0 | 0 | 0 | 1 | 1 | 1 | 0 | 0 | 1 | 4 | 0 | 1 | 3 | 0 | 0 | 4 | 2 | 1 |
| 2q95 | A05 | | | 0 | 0 | 0 | 0 | 0 | 1 | 1 | 1 | 0 | 0 | 0 | 4 | 0 | 1 | 3 | 0 | 0 | 4 | 2 | 1 |
| 2q96 | A18 | | | 0 | 0 | 0 | 0 | 0 | 1 | 1 | 1 | 1 | 0 | 1 | 4 | 0 | 1 | 3 | 0 | 0 | 4 | 2 | 1 |
| 2q92 | B23 | | | 0 | 0 | 0 | 0 | 0 | 1 | 1 | 1 | 0 | 0 | 1 | 4 | 0 | 1 | 3 | 0 | 0 | 4 | 2 | 1 |
| 1xnz | FCD | | | 0 | 0 | 0 | 0 | 0 | 1 | 1 | 1 | 0 | 0 | 1 | 4 | 0 | 1 | 3 | 0 | 0 | 3 | 2 | 1 |
| 3iu7 | FCD | | | 0 | 0 | 0 | 0 | 0 | 1 | 3 | 1 | 0 | 0 | 3 | 2 | 0 | 1 | 1 | 1 | 0 | 3 | 2 | 1 |
| 3d27 | W29 | | | 0 | 0 | 0 | 0 | 0 | 1 | 1 | 1 | 0 | 0 | 1 | 4 | 0 | 0 | 3 | 0 | 0 | 3 | 2 | 1 |
| 3iu9 | T07 | | 3e | 0 | 0 | 0 | 0 | 0 | 1 | 3 | 1 | 0 | 1 | 3 | 2 | 0 | 1 | 2 | 1 | 0 | 3 | 2 | 2 |
| 3iu8 | T03 | | | 0 | 0 | 0 | 0 | 0 | 1 | 3 | 1 | 0 | 0 | 2 | 2 | 0 | 1 | 1 | 0 | 0 | 3 | 2 | 1 |

*Properties of the aminopeptidase pockets that bound to the inhibitors present in the clusters 0-1* ***(A)*** *and cluster 3* ***(B).*** *Residue contents of the 20 amino acids arranged according to their physicochemical properties and colored in increasing green intensity from min=0 to max=8.*

**Supplementary Table CST4 -** ***Properties of the aminopeptidase pockets that bound to the inhibitors present in the clusters 2.***

| | | | | Hydrophobic | | | | | | | | Hydrophilic | | | | | | | Positive | | | Negative | |
|---|---|---|---|---|---|---|---|---|---|---|---|---|---|---|---|---|---|---|---|---|---|---|---|
| pdb | LIG | LIG_clusters | sub_cluster | A | V | L | I | P | M | F | W | G | S | T | C | N | Q | Y | K | R | H | D | E |
| 4zx6 | 4U6 | 2 | 2a | 2 | 3 | 0 | 0 | 0 | 1 | 1 | 0 | 1 | 0 | 4 | 0 | 2 | 1 | 2 | 0 | 1 | 1 | 1 | 4 |
| 4zx4 | 4TL | | | 2 | 3 | 0 | 0 | 0 | 1 | 1 | 0 | 2 | 0 | 5 | 0 | 1 | 1 | 2 | 0 | 1 | 1 | 1 | 5 |
| 4zx5 | 4TM | | | 2 | 3 | 0 | 0 | 0 | 1 | 0 | 0 | 1 | 0 | 2 | 0 | 1 | 1 | 2 | 0 | 1 | 1 | 1 | 5 |
| 6ea2 | J1G | | | 2 | 3 | 0 | 0 | 0 | 2 | 1 | 0 | 2 | 0 | 3 | 0 | 1 | 1 | 2 | 0 | 1 | 1 | 1 | 6 |
| 6eab | J2D | | | 2 | 3 | 0 | 0 | 0 | 1 | 1 | 0 | 2 | 0 | 3 | 0 | 1 | 1 | 2 | 0 | 1 | 1 | 1 | 5 |
| 6ee6 | J4P | | | 2 | 3 | 0 | 0 | 0 | 2 | 1 | 0 | 2 | 0 | 3 | 0 | 1 | 1 | 2 | 0 | 1 | 1 | 1 | 6 |
| 6ee4 | J4S | | | 2 | 2 | 0 | 0 | 0 | 2 | 1 | 0 | 2 | 0 | 4 | 0 | 1 | 1 | 2 | 0 | 1 | 1 | 1 | 5 |
| 6eed | J6A | | | 2 | 2 | 0 | 0 | 0 | 1 | 1 | 0 | 2 | 0 | 4 | 0 | 1 | 1 | 2 | 0 | 1 | 1 | 1 | 7 |
| 4k5o | 1OT | | | 2 | 2 | 1 | 0 | 0 | 2 | 0 | 0 | 1 | 0 | 2 | 0 | 1 | 1 | 2 | 0 | 1 | 1 | 0 | 5 |
| 4zw5 | 4SA | | | 2 | 3 | 0 | 0 | 0 | 1 | 1 | 0 | 2 | 0 | 3 | 0 | 1 | 1 | 2 | 0 | 1 | 1 | 1 | 6 |
| 6ea1 | J0Y | | | 2 | 3 | 0 | 0 | 0 | 1 | 1 | 0 | 2 | 0 | 5 | 0 | 1 | 1 | 2 | 0 | 1 | 1 | 1 | 6 |
| 6ee3 | J4V | | | 2 | 3 | 0 | 0 | 0 | 2 | 1 | 0 | 2 | 0 | 5 | 0 | 1 | 1 | 2 | 0 | 2 | 1 | 1 | 6 |
| 5y1r | 8KR | | | 2 | 3 | 0 | 0 | 0 | 1 | 1 | 0 | 1 | 0 | 2 | 0 | 1 | 1 | 2 | 0 | 1 | 1 | 1 | 6 |
| 4zw6 | 4SY | | | 2 | 3 | 0 | 0 | 0 | 1 | 1 | 0 | 2 | 0 | 3 | 0 | 1 | 1 | 2 | 0 | 1 | 1 | 1 | 5 |
| 4zw7 | 4SZ | | | 2 | 3 | 0 | 0 | 0 | 2 | 1 | 0 | 2 | 0 | 3 | 0 | 1 | 1 | 2 | 0 | 1 | 1 | 1 | 6 |
| 4zw8 | 4T2 | | | 2 | 3 | 0 | 0 | 0 | 2 | 1 | 0 | 2 | 0 | 3 | 0 | 1 | 1 | 2 | 0 | 1 | 1 | 0 | 5 |
| 4zx3 | 4TK | | | 2 | 3 | 0 | 0 | 0 | 1 | 0 | 0 | 1 | 0 | 3 | 0 | 1 | 1 | 2 | 0 | 1 | 1 | 1 | 6 |
| 5xm7 | 89X | | | 2 | 4 | 0 | 0 | 0 | 1 | 0 | 0 | 1 | 0 | 3 | 0 | 1 | 1 | 2 | 0 | 1 | 1 | 1 | 7 |
| 5y1v | 8LO | | | 2 | 3 | 0 | 0 | 0 | 1 | 0 | 0 | 1 | 0 | 3 | 0 | 1 | 1 | 2 | 0 | 0 | 1 | 1 | 5 |
| 5y1t | E8G | | | 1 | 2 | 0 | 0 | 0 | 1 | 1 | 0 | 1 | 0 | 1 | 0 | 1 | 1 | 2 | 0 | 0 | 1 | 0 | 5 |
| 3q7j | FBO | | | 2 | 1 | 2 | 0 | 0 | 1 | 1 | 0 | 3 | 1 | 2 | 0 | 0 | 1 | 2 | 0 | 1 | 2 | 1 | 4 |
| 6eaa | J1V | | | 2 | 3 | 0 | 0 | 0 | 1 | 1 | 0 | 2 | 0 | 3 | 0 | 1 | 1 | 2 | 0 | 1 | 1 | 1 | 6 |
| 3mdj | BES | | | 1 | 0 | 1 | 0 | 0 | 2 | 0 | 0 | 1 | 1 | 1 | 0 | 0 | 1 | 1 | 2 | 1 | 2 | 2 | 4 |
| 3ebh | BES | | | 2 | 3 | 0 | 0 | 0 | 1 | 0 | 0 | 1 | 0 | 2 | 0 | 1 | 1 | 2 | 0 | 1 | 1 | 0 | 6 |
| 4fkk | BES | | | 2 | 1 | 0 | 0 | 0 | 1 | 1 | 0 | 1 | 3 | 0 | 0 | 0 | 2 | 1 | 0 | 1 | 1 | 1 | 4 |
| 3q44 | D50 | | | 2 | 3 | 0 | 0 | 0 | 2 | 0 | 0 | 1 | 1 | 3 | 0 | 1 | 1 | 2 | 0 | 1 | 1 | 1 | 6 |
| 4jbs | P52 | | | 2 | 1 | 0 | 0 | 2 | 1 | 1 | 1 | 2 | 2 | 0 | 0 | 1 | 1 | 2 | 1 | 3 | 1 | 3 | 8 |
| 3ebi | BEY | | | 2 | 3 | 1 | 0 | 0 | 1 | 1 | 0 | 1 | 0 | 2 | 0 | 1 | 1 | 2 | 0 | 1 | 1 | 1 | 6 |
| 2zxg | S23 | | | 1 | 2 | 1 | 0 | 0 | 1 | 0 | 0 | 1 | 0 | 1 | 0 | 2 | 1 | 3 | 0 | 2 | 1 | 2 | 5 |
| 5y1x | BB2 | | | 1 | 3 | 0 | 0 | 0 | 1 | 0 | 0 | 1 | 0 | 3 | 0 | 2 | 0 | 2 | 1 | 2 | 1 | 1 | 5 |
| 5y1q | 8L0 | | | 2 | 2 | 0 | 0 | 0 | 1 | 0 | 0 | 1 | 0 | 0 | 0 | 2 | 1 | 2 | 0 | 2 | 1 | 0 | 5 |
| 5y1k | B1B | | | 2 | 2 | 0 | 0 | 0 | 1 | 1 | 0 | 1 | 0 | 1 | 0 | 1 | 1 | 2 | 0 | 1 | 1 | 0 | 5 |
| 4x2u | TOD | | | 1 | 3 | 0 | 0 | 0 | 1 | 0 | 0 | 1 | 0 | 2 | 0 | 3 | 1 | 2 | 0 | 2 | 1 | 1 | 5 |
| 4r7l | SHH | | | 2 | 1 | 1 | 0 | 2 | 1 | 1 | 1 | 2 | 1 | 0 | 0 | 0 | 2 | 3 | 0 | 0 | 1 | 1 | 3 |
| 4zqt | 4QP | | | 2 | 3 | 0 | 0 | 0 | 1 | 1 | 0 | 1 | 0 | 1 | 0 | 1 | 1 | 2 | 0 | 1 | 1 | 1 | 6 |
| 4zw3 | 4S9 | | | 2 | 3 | 0 | 0 | 0 | 1 | 0 | 0 | 1 | 0 | 1 | 0 | 1 | 1 | 2 | 0 | 1 | 1 | 1 | 6 |
| 5y19 | 8KO | | | 2 | 3 | 0 | 0 | 0 | 1 | 0 | 0 | 1 | 0 | 1 | 0 | 1 | 1 | 2 | 0 | 2 | 1 | 1 | 6 |
| 3t8v | BTJ | | | 3 | 3 | 0 | 0 | 0 | 1 | 0 | 0 | 1 | 0 | 3 | 0 | 1 | 1 | 2 | 0 | 1 | 1 | 1 | 6 |
| 3q43 | D66 | | | 2 | 3 | 0 | 0 | 0 | 1 | 0 | 0 | 1 | 0 | 2 | 0 | 1 | 1 | 2 | 0 | 1 | 1 | 1 | 6 |
| 4k5m | 1OV | | | 1 | 2 | 0 | 0 | 0 | 1 | 1 | 0 | 1 | 0 | 1 | 0 | 1 | 1 | 2 | 0 | 0 | 1 | 0 | 5 |
| 4k5l | 19N | | | 2 | 2 | 1 | 0 | 0 | 1 | 1 | 0 | 1 | 0 | 2 | 0 | 1 | 1 | 2 | 0 | 0 | 1 | 0 | 5 |
| 4k5n | 1OU | | | 2 | 1 | 0 | 0 | 0 | 1 | 1 | 0 | 1 | 0 | 1 | 0 | 1 | 1 | 2 | 0 | 1 | 1 | 0 | 5 |
| 4k5p | 1OS | | | 2 | 2 | 0 | 0 | 0 | 2 | 1 | 0 | 1 | 0 | 1 | 0 | 1 | 1 | 2 | 0 | 1 | 1 | 0 | 5 |
| 4pw4 | 2X0 | | | 1 | 1 | 0 | 0 | 1 | 0 | 1 | 0 | 1 | 0 | 0 | 0 | 2 | 2 | 2 | 0 | 0 | 1 | 0 | 4 |
| 4pqa | X8Z | | 2b | 0 | 0 | 0 | 1 | 1 | 0 | 0 | 0 | 2 | 0 | 2 | 0 | 1 | 0 | 0 | 1 | 1 | 1 | 0 | 3 |

*Properties of the aminopeptidase pockets that bound to the inhibitors present in the clusters 2. Residue contents of the 20 amino acids arranged according to their physicochemical properties and coloured in increasing green intensity from min=0 to max=8.*

It was shown that the hydrophilic tyrosine (Y) appeared in majority (102/107) of the aminopeptidase pockets, and both D and E in most of the metal-containing pockets of the aminopeptidase complexes. Since these features are also present in the GPAA1 active sites [1,6], it additionally reaffirms that the GPAA1 could function like an aminopeptidase and, therefore, supports the reliability of our GPAA1 structural models [1,6] where Zn ions recruit the residue Y328 into their active sites.

#### II. Docking analyses of aminopeptidase inhibitors to the GPAA1 active site

The experimental screening of aminopeptidase inhibitors revealed that Bestatin (BES) repressed GPAA1 activity. To explore BES-binding pockets on the GPAA1 structures attaching one or two zinc as previously found [1], namely $GPAA1^{Zn}$ and $GPAA1^{ZnZn}$, we first performed various blind docking attempts without prior knowledge of the BES binding regions using AutoDock Vina [16]. For both the $GPAA1^{Zn}$ and $GPAA1^{ZnZn}$ models, we found a pocket of interest involving the GPAA1 Zn-binding sites (residues D153, D188, Y328 and/or E226 [6] to be the most frequently visited by the BES molecule. This result provided a glimpse of potential BES binding mode. The features were surprisingly in line with the pharmacophore patterns detected in various aminopeptidase-inhibitor-complexes (such as those in PDB: 4zx8 or 3mdj). Therefore, we explored the binding modes of the BES on the GPAA1 by subsequently docking it attentively to the GPAA1 Zn(s)-binding region using various replicates of the $GPAA1^{Zn}$ and $GPAA1^{ZnZn}$ conformations, using Glide v.8.8 [17].

The computations revealed that the BES molecule makes direct contacts with the Zn(s) located in the $GPAA1^{Zn}$ and $GPAA1^{ZnZn}$ structures, with binding energy $\Delta g$ (kcal/mol) of $-5.7 \pm 1.04$ and $-5.14 \pm 0.07$, respectively (Supplementary Figure CSF4). Compared with the reference aminopeptidase-BES complexes attached with one zinc (PDB: 3mdj) and two zincs (PDB: 5ib9), the GPAA1 binding of BES shared similar binding modes where a hydrogen bond acceptor pair was involved in binding to the zincs of the GPAA1 active site. It was also noticed that the BES interacted predominantly to one of the two zinc atoms in the $GPAA1^{ZnZn}$. This suggests that, for the aminopeptidase-like function of the GPAA1, attachment of one zinc ion in its active site might be sufficient and that this might be the more dominant arrangement.

**Supplementary Figure CSF4**

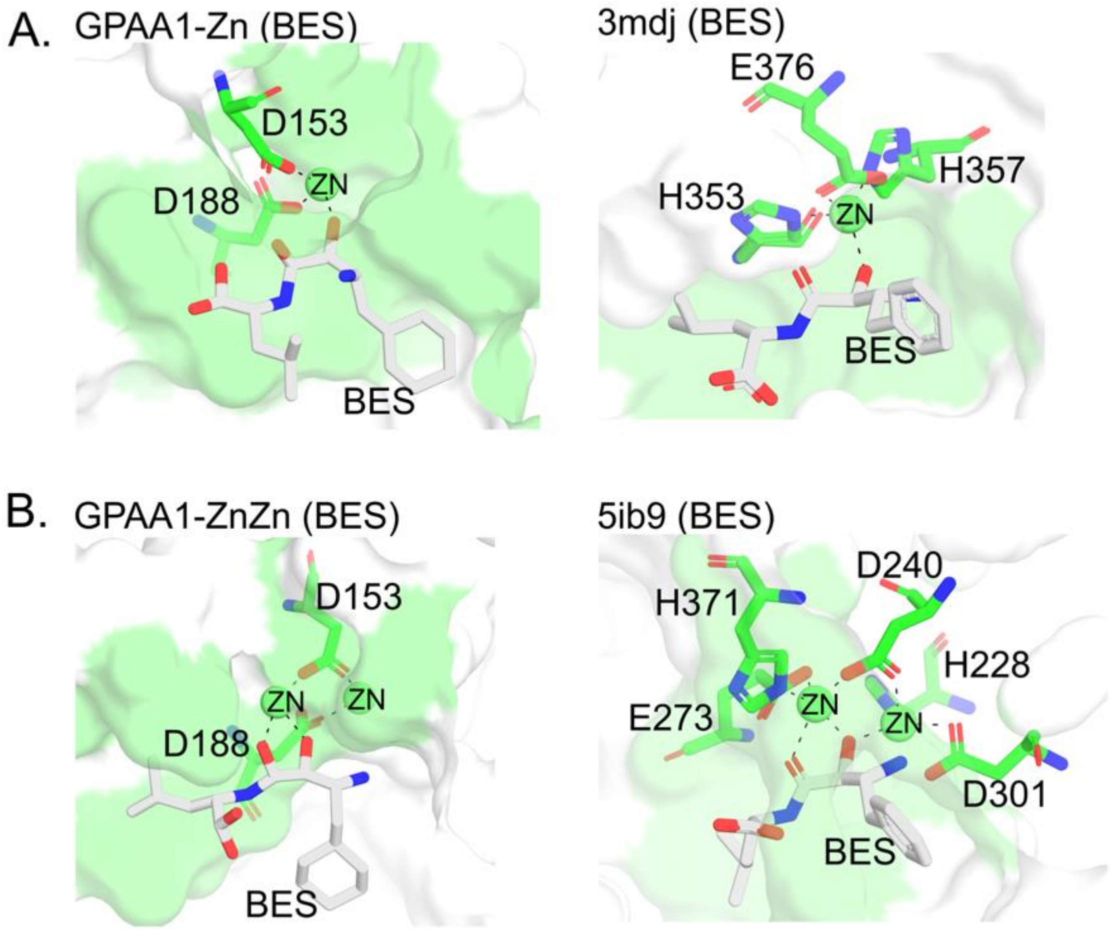


*Analysis of the complex resulting from GPAA1 and bestatin (BES) binding. The BES binding modes in the GPAA1$^{Zn}$* ***(A)*** *and GPAA1$^{ZnZn}$* ***(B)*** *as compared to the referenced aminopeptidase-BES complexes (right panels), respectively PDB: 3mdj (one zinc) and PDB: 5ib9 (two zincs). The BES molecule is shown in sticks (carbon: white, nitrogen: blue, oxygen: red, others: yellow, hydrogen not shown for simplicity), the aminopeptidase pockets are shown in surface with the ZN in spheres. The pocket residues that are within 4 Å around the BES are shown in green. The Zn-interacting residues are labelled.*

#### III. Computational analysis of known clinical mutations of GPAA1

Disorders in neural development or genetic anomalies have been found to be associated with GPAA1 mutations [18-20]. Among these reported mutations, a few missense mutations on GPAA1 were reported to likely affect the protein stability [19,20]. With the goal of analyzing these substitution mutations, we performed mutagenesis on our GPAA1 models to generate the mutant structures, and investigated the protein stability in the presence of the mutations, using FoldX 5.0 [21]. Preliminary results of computational alanine scanning upon the GPAA1$^{Zn}$ structure highlighted the stability sensitivity where the majority of mutated positions would perturb its structural stability (Supplementary Figure CSF5A).

**Supplementary Figure CSF5 – GPAA1 protein stability changed by amino acid residue mutations**

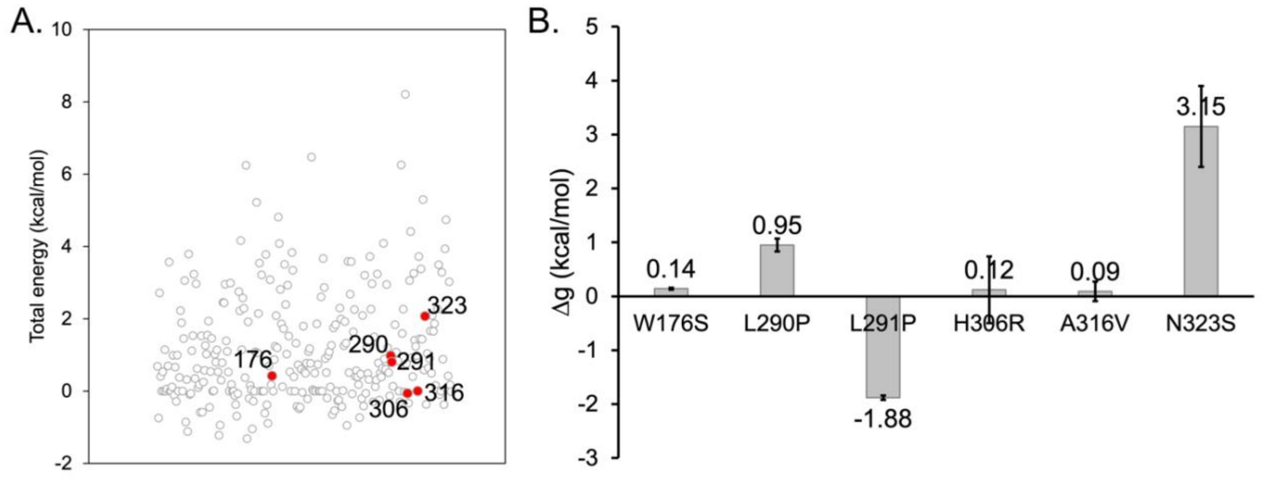


*Computational mutagenesis of clinical mutations on the GPAA1 models.* ***(A)*** *Alanine scanning results of the GPAA1$^{Zn}$ model. The positions of the clinical mutations are highlighted in red and labelled. (**B)** Single point mutagenesis of a few reported clinical mutations depicting destabilizing effects (Δg>0) on the GPAA1$^{Zn}$ structural stability.*

Given that there was no combination of these missense mutations reported in a clinical case (despite some with indel mutations), we modelled the mutations individually. Six mutations W176S, L290P, L291P [20], N323S [19], and H306R, A316V [18] were modelled on the GPAA1$^{Zn}$ structure and their respective structural free energies were estimated. Difference in free energy (Δg) compared to the wildtype GPAA1$^{Zn}$ demonstrates the protein stability upon the occurred mutation, i.e., Δg<0: stabilized or Δg>0: destabilized.

Among these mutations, the residues L290 and L291 are located at the loop4 – one of the two flaps at the Zn-binding sites detected in our previous study [1] that exhibited large fluctuations in the presence of the substrate. Except for the L291P, the mutations demonstrated destabilizing effects (Δg>0), particularly most pronounced with the N323S, possibly leading the GPAA1$^{Zn}$ activity disrupt due to its structural instability (Supplementary Figure CSF5B). The destabilizing effect by the N323S was in consistent with the finding by Li and colleagues [19] where the mutation was reported to induce structure changes at the functional domain. On the other hand, the mutation L291P contributed to stabilizing the GPAA1 structure (Δg<0).

An additional attempt of double mutation L290P-L291P resulted in low Δg (-0.34 ± 0.11 kcal/mol, data not shown). It is speculated that the mutation L291P might be able to overwhelm the effect of the L290P and restore the overall stability. Since the loop4 may play a role in modulating the substrate accessibility to the active sites [1], effects of the mutations might also augment to these modulating effects.

#### IV. Comparison of structural models of Gpaa1 and cryo-EM structures of the transamidase complex (Discussion)

Our models of the lumenal GPAA1 domain maintains similar structural folds as those in the whole GPIT complexes such as the two recently published Cryo-EM structures by Zhang *et al.*[4] (PDB: 7w72) and Xu *et al.*[5] (PDB: 7wld) or models predicted by AlphaFold[2] (UnitProt: O43292) and David Baker's group[3] (UnitProt: P39012).

The individual GPAA1 subunits (with TMs) in these whole GPIT structures were in RMSD range of ~0.8–2.2 Å from each other when structurally aligned, whereas our GPAA1 models (without TMs) were in ~2.7–4.0 Å when aligning with the corresponding luminal region in each whole GPIT (excluding the zinc ions bound to GPAA1 model conformations when aligned to PDB:7w72 and 7wld, RMSD ~5.3 and ~5.9 Å, respectively; see Supplementary Table CST5 placed after the Methods Appendix). It suggests that the GPAA1 subunit, while preserving its main secondary structural fold elements, bears substantial flexibility with respect to the whole GPIT complex (including GPAA1, PIG-U, PIG-K, PIG-T, and PIG-S).

Indeed, the GPAA1 exhibits collective motions with PIG-K and PIG-S, approaching PIG-K in an opposite direction to PIG-S (shown in Supplementary Figure CSF6) while PIG-S holds and embraces PIG-K catalytic domain [5].

**Supplementary Figure CSF6 – Normal mode analysis of the GPIT complex**

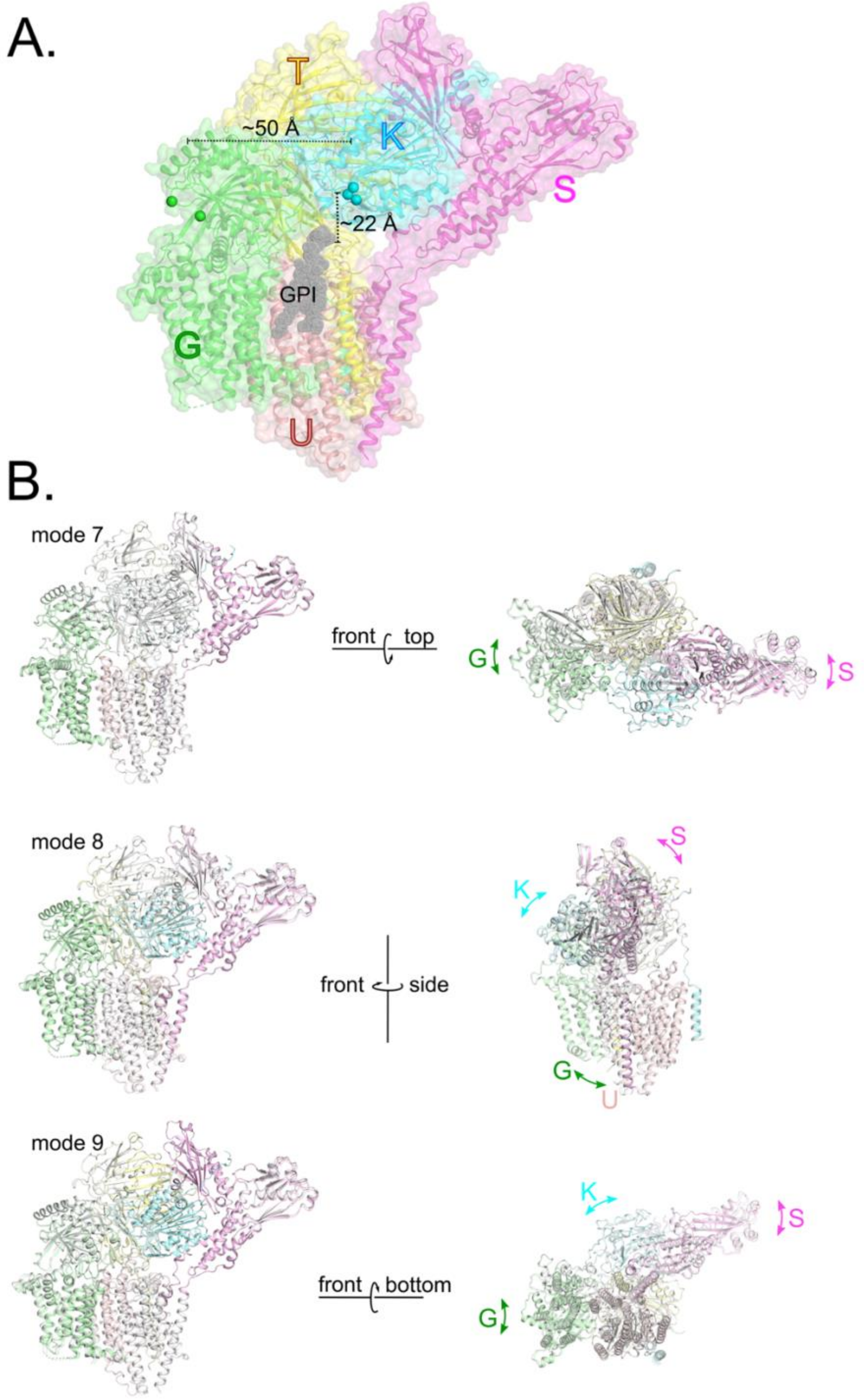


*Sufficient space orienting GPAA1 and PIG-K to accommodate the GPI and proprotein (A). Collective motions of GPAA1, PIG-K, and PIG-S, using normal mode analysis server ElNemo [22]. The first 3 non-trivial modes are presented.*

Given that the GPI-attaching cavity is in ~22–25 Å to PIG-K catalytic dyad and GPAA1 active site is ~50 Å distant from this catalytic site [5], the correlated motions of the three domains is to orient the GPI-attaching cavity as well as accommodate the proprotein substrate for the catalysis. In this scenario, the GPAA1 luminal domain approaching the *ω*-site in the presence of the proprotein and GPI at the PIG-K catalytic site collectively with the motions of both PIG-K and PIG-S (perhaps slightly down towards the GPI cavity), is possible. In addition, it was reported that GPAA1 was physically close to the GPIT-bound proprotein [23-25]; thus, this hypothesis of GPAA1 approaching the PIG-K catalytic site should not be neglected without an experimentally determined structure of the proprotein-bound GPIT complex to demonstrate the otherwise.

##### V. Implication of the GPAA1 structure on the aminopeptidase-like function of GPAA1 and a proposed GPAA1 reaction mechanism

We explored structural descriptors that defined the aminopeptidase active sites and the pharmacophores of aminopeptidase inhibitors to propose the possible aminopeptidase-like function of the human GPAA1. Our hypothesis triggers reconsideration on a few points that Xu and colleagues[5] discussed previously on the GPAA1 function.

Xu *et al.* negated the zinc-binding capability of GPAA1 [5]. We disagree with this hypothesis. First, in the zinc-binding protein families e.g. M28 family, the zinc are coordinated via sites 2, 3 (aspartate, D and/or glutamate, E) and site 5 (histidine, H) in the case of one zinc or additionally to site 1 (H) and site 4 (D or E) in the case of two zincs [6]. Our previous study [1] showed that in the GPAA1 and several M28 protein structures (PDB: 4fwu, 4f9u, 1amp, 1f2o), sites 2, 3 and/or 4 contributed the most to the zinc binding energies, indicating the more essential attribute of D/E than H in binding zinc in the catalytic sites. In particular, sites 2 and 3 in the GPAA1 active site could complement each other to bind zinc in the presence of different substrates [1]. In place of the histidine, a tyrosine (Y) present at the site 5 of GPAA1 also maintains the zinc binding capability.

Second, Xu *et al.* demonstrated that single point mutagenesis to disrupt the zinc-binding residue candidates of GPAA1 (D153A, D188A, etc.) still maintained the GPIT activity in cells, hence again rejecting the GPAA1 zinc binding ability [5]. It is important to note that we could indirectly reproduce the similar resulting effect of the single point mutagenesis (D153A or D188A, see main text of this article) on the GPIT activity. Yet, our double D153A-D188A mutagenesis results (see main text of this article) affirms the zinc-binding capability of GPAA1 and further supports the complementary contribution in binding zinc of the two key residues D153 (site 2) and D188 (site 3).

The PIG-K catalytic site seems not be as legumain-like as Xu *et al.* had proposed [5]. Although PIG-K shares similar topology to legumain/caspase and both the enzymes can be active in the ER lumen neutral pH 7.2 ± 0.2 [26] (with legumain having dual activities switched due to pH shift between acidic and neutral pH [27] and caspase active at pH 6.8–7.2 [28,29]), the PIG-K catalytic region including the trivalent oxyanion hole (C206, H164, and G165, PDB: 7wld numbering) shares more similar distribution of electrostatics potentials to those of caspases than to those in legumain (Supplementary Figure CSF7). In addition, the substrate selectivity pocket S1 of both PIG-K and caspase reflects better zwitterionic characteristic with both positively and negatively charged regions. Also, the trio C206, H164, and G165 are conserved in all human caspase domains [30]. Hence, it suggests PIG-K catalytic mechanism be caspase-related, supporting a previous study[31].

**Supplementary Figure CSF7 - Electrostatics potential analysis of GPIT components**

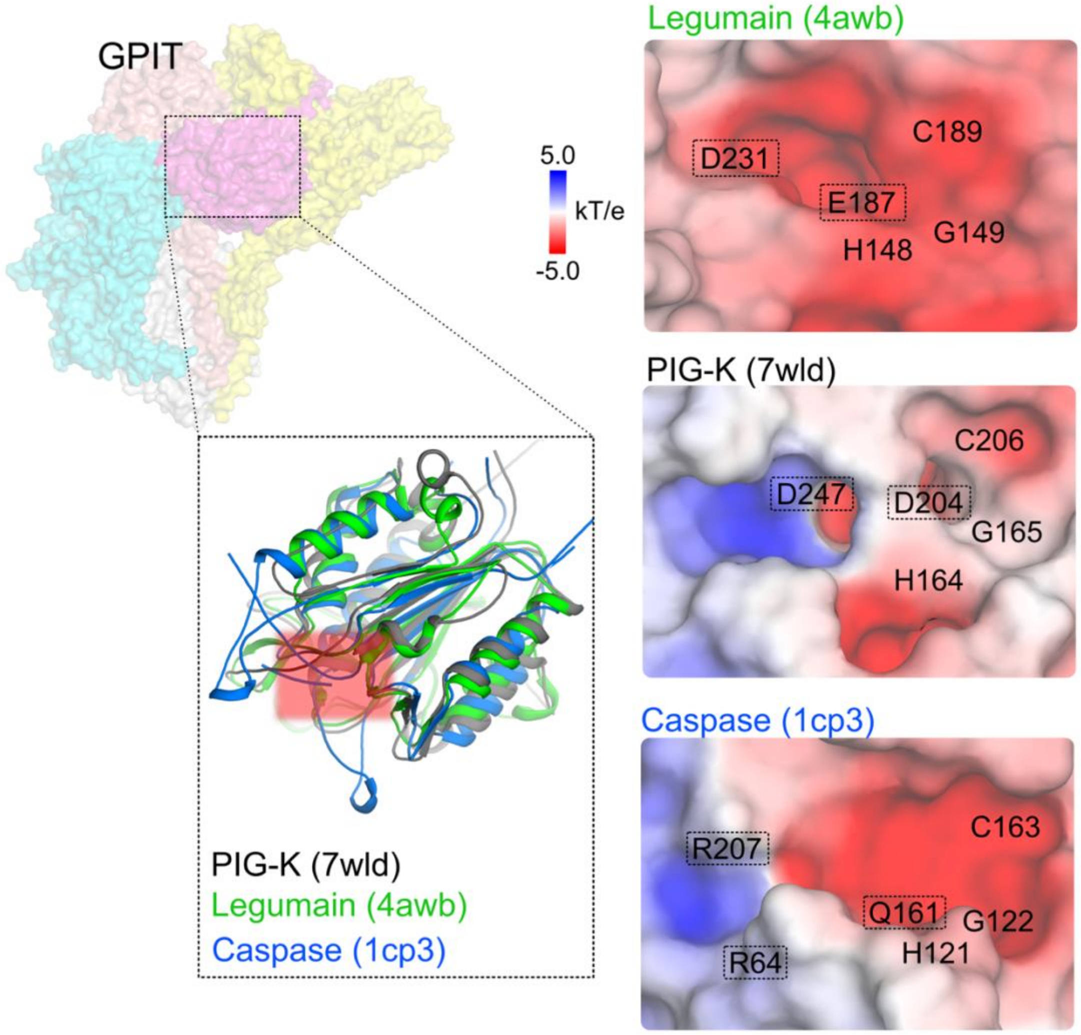


*Electrostatics potentials of the catalytic sites of PIG-K (PDB: 7wld), legumain (PDB: 4awb), and caspase (PDB: 1cp3). The involved residues are labelled, including the trivalent oxyanion hole cysteine (C), histidine (H), and glycine (G) with respective numbering in the PDBs, and the S1 pocket residues (boxed). The S1 pocket residues are referenced from Xu et al.* [5] *for the PIG-K and legumain and from Fuentes-Prior et al.* [30] *for the caspase. The electrostatics calculation was performed using APBS plugin in PyMOL v2.1* [32]*.*

Given the PIG-K caspase-like catalytic activity, which is zinc-dependent [33], together with the GPAA1 zinc-binding capability, we propose a zinc-related regulatory role of

GPAA1 on the GPI attachment at PIG-K catalytic site, as below (Supplementary Figure CSF8).

**Supplementary Figure CSF8 – GPIT reaction mechanism**

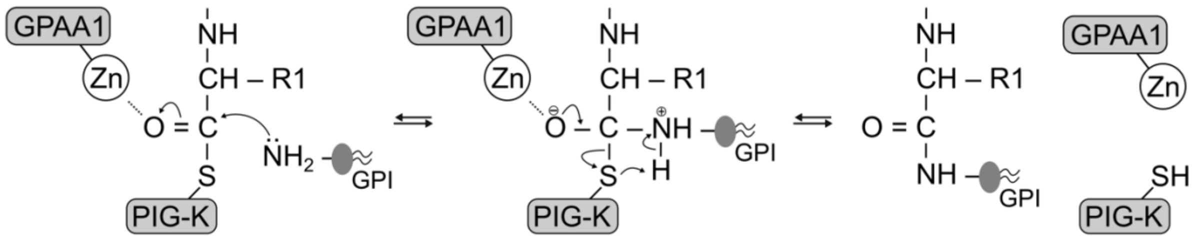


*Proposed mechanism of the GPI attachment to the proprotein substrate involving the zinc-bound GPAA1 at PIG-K catalytic site. In this process, the zinc-bound GPAA1 mimics the electrophilic catalyst of the PIG-K G165 (PDB:7wld) by coordinating to the carbonyl oxygen of the proprotein substrate.*

It was found that GPAA1 was physically close to the GPIT-bound proprotein and crucial for the recognition of the proprotein [23-25]. In addition, Ohishi *et al.* reported that GPAA1 acted "before and during" the carbonyl intermediate formation and was required for the cleavage at $\omega$-site of the proprotein [34]. Therefore, it is reasonable to argue that GPAA1 could also interact with the GPIT-bound proprotein.

There are free zincs in the ER lumen [35], hence it is, to some extent, certain that the GPAA1 active sites (containing aspartate, glutamate, and tyrosine) would bind to zinc (either one or two) in tetra-coordination manners. The zinc-bound GPAA1 active sites could subsequently accommodate and interact with the proprotein via the zinc coordination, e.g. as modelled in our previous work [1]. It is postulated that the zinc-bound GPAA1 could coordinate with the carbonyl oxygen of the scissile peptide bond at the $\omega$-site to trigger an attack to the thioester bond, formed by PIG-K and the proprotein substrate [24], for the GPI attachment. The non-bonded zinc coordinated at the carbonyl oxygen mimics an electrophilic catalyst [36] to facilitate the nucleophilic attack by the GPI amine group on the EtNP3 to form the necessary tetrahedral intermediate for the subsequent hydrolysis (Supplementary Figure CSF8). The zinc-mediated reaction was one of the proposed catalytic mechanisms in the metallo-aminopeptidase family [36], to which GPAA1 belongs [6]. The aminopeptidase-like characteristic of GPAA1 was affirmed by experimental results described in the main text of this work.

On another note, zinc was reported to inhibit various caspases (cysteine-histidine catalytic enzymes) directly or allosterically and/or competitively to the enzyme catalytic site, and in concentration-dependent manner [33]. Given that zincs are present in the ER lumen [35], the presence of zinc-binding GPAA1 alongside with PIG-K could also contribute a regulatory role onto the caspase-like catalytic activity of the PIG-K, e.g. GPAA1 taking up zinc to alleviate the affinity between zinc and PIG-K.

#### METHODS APPENDIX

##### Descriptor extraction of inhibitor-binding pocket features and common pharmacophores of aminopeptidase inhibitors

Inhibitor-bound aminopeptidase complexes were identified from Protein Data Bank (PDB, accessed in October 2021) using the “Advanced Search” with the query containing a combination of additional structure keywords: “aminopeptidase” and “inhibitor”. Only 107/121 aminopeptidase complexes that contain small molecules as inhibitors were retrieved for analysis.

The *dpocket* module implemented in the *fpocket 2.0* package [10] was used to describe the inhibitor-bound pocket properties such as the amino acid propensity, volume, hydrophobicity and charge scores, as well as the ligand volume. The binding energies were estimated using the PRODIGY-LIG package [37].

PharmaGist server [11] was used to identify pharmacophores, e.g. aromatic, hydrophobic, hydrogen bond donor, and/or acceptor groups, present in the set of 107 aminopeptidase-inhibitor complexes. The resulting features of the inhibitor pharmacophores and the inhibitor-bound pocket amino acid propensities were used to first group the inhibitor set using *k-means* clustering method implemented in the scikit-learn v.0.23.2 package [12]. Subsequently, the clusters were further sub-grouped based on the metal binding capacity of the aminopeptidase active sites, e.g. lacking metal (*no_metal*), with direct metal binding (*#num_metal* > 0), without direct metal binding (*#num_metal* = 0), or different types of metals (ZN, CO, MN, NI, etc.).

The common pharmacophores in each sub-cluster of inhibitors were identified using the PharmaGist server with the first molecule in each sub-cluster assigned as the pivot molecule. Only the alignment against the pivot molecule with the maximum alignment score was selected for each sub-cluster. In cases of a partial alignment (i.e., not all molecules are fully aligned against the pivot molecule), the remaining "not aligned" set of each sub-cluster would undergo the pharmacophore re-search and a new pivot molecule would be re-assigned. If no alignment could be achieved, pharmacophore search for the single molecule would be performed. The detailed process description can be found in Supplementary Figure CSF9. The common pharmacophores were visualized and analysed using the ZINC-Pharmer [13].

**Supplementary Figure CSF9**

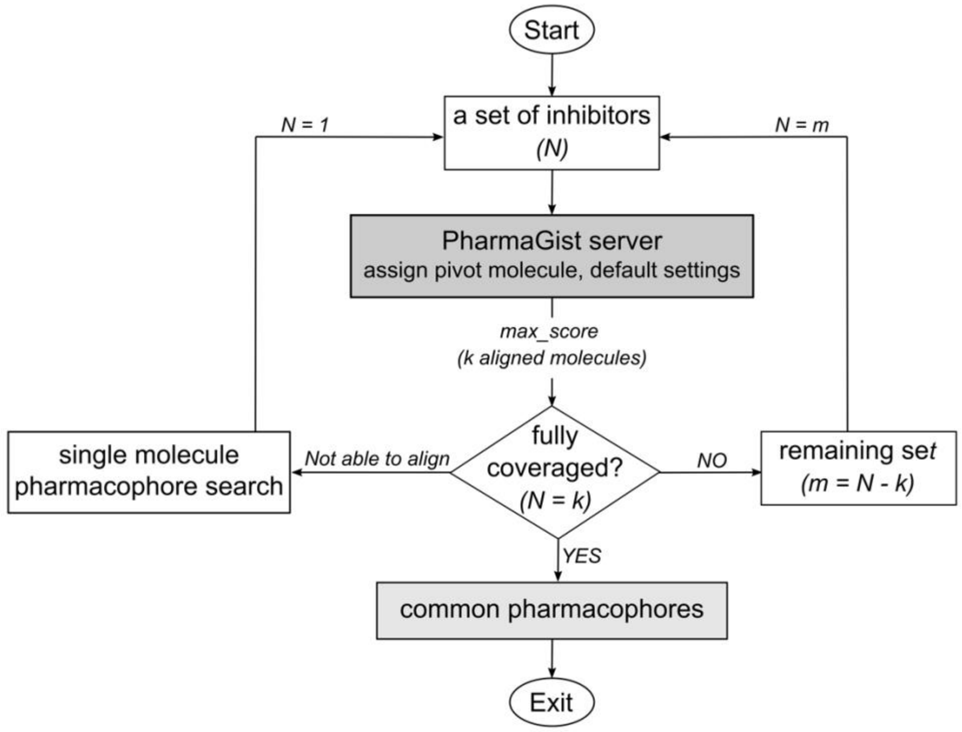


*Pharmacophore search process of the retrieved set of 107 aminopeptidase inhibitors.*

**Docking analyses of bestatin (BES) to the GPAA1 active sites**

Ten replicates were extracted from the molecular dynamics (MD) trajectories of the Zn(s)-bound GPAA1 in our previous study [1] and used to first explore the potential BES-binding pocket on the $GPAA1^{Zn}$ and $GPAA1^{ZnZn}$ structures. Blind docking was first initiated on the 10 replicates (10×1000 binding modes) using AutoDock Vina [16] with a grid box centering and covering the whole GPAA1 structure (as receptor). The BES molecule retrieved from PubChem (CID: 72172) was used as the ligand. Only those pockets that involved the GPAA1 active sites and the Zn atom(s) and that were most frequently visited by the ligand were of interest.

The previous 3×300 ns MD samplings of the GPAA1$^{Zn}$ and GPAA1$^{ZnZn}$ models [1] were extended to 3×500 ns using the same parameters as previously. Ten replicates were extracted from the last 300 ns of the new sampling for further docking refinement.

In the subsequent focus docking refinement, we used Glide version 8.8 implemented in the Schrödinger package, Release 2020-3, [17]. The BES molecule was prepared and minimized using the *LigPrep* module. Then, it was docked attentively to the detected region of interest above, i.e., the proximity of the Zn(s)-bound active sites of the GPAA1 structures resulted from the blind docking experiment.

For the receptor, a grid box of 7 Å (or 10 Å) was set centering the zinc ion (or two zinc ions, respectively) coordinated with the GPAA1 active sites that include D153, D188, Y328 and/or E226 [6]. The docking protocol with default settings was performed on the extracted 10 replicates of the GPAA1$^{Zn}$ or GPAA1$^{ZnZn}$ conformations. Only those resulting in docked complexes with the BES binding modes satisfying the detected pharmacophores (i.e., in sub-cluster 2a or 1c shown in the Supplementary Figure CSF3 because BES is one of the aminopeptidase inhibitors found in those sub-clusters) were selected for further analyses. The binding energies were estimated using the PRODIGY-LIG server [37].

**Computational mutagenesis of the GPAA1 structures**

Among the clinical mutations, six mutations W176S, L290P, L291P [20], N323S [19], and H306R, A316V [18] are covered within the length of our lumenal GPAA1 models [1]. They were individually modelled on the GPAA1$^{Zn}$ structure using FoldX 5.0 [21]. For each of the mutations, 100 replicates of the mutant structures were generated for statistical reliability. The respective structural free energy differences between the wildtype and mutants (Δg) were then estimated to investigate the protein structural stability.

The alanine scanning was also performed on the GPAA1$^{Zn}$ structure using the *AlaScan* module implemented in the FoldX 5.0 package.

*Supplementary Table CST5: Structural alignment of various domain models of the GPIT complex.*

| Our GPAA1 models | AlphaFold models (UnitProt) | Baker's models # (Humphrey *et al.*, *Science*) | PDB:7w72 (Zhang *et al.*, *Nature SB)* | PDB:7wLD (Xu *et al.*, *Nature Comm)* | RMS (Å) | Alignment command in PyMOL |
|---|---|---|---|---|---|---|
| | | Whole GPIT | Whole GPIT | | 2.970 | `cealign 7w72 and ss h+b, bk_GPIT and ss b+h` |
| | | | Whole GPIT | Whole GPIT | 2.962 | `cealign 7w72 and ss h+b, 7wld and ss b+h` |
| | Gpaa1_O43292 | Gaa1_P39012 (w TMs) | - | | 1.907 | `super AF-O43292_Gpaa1, bk_gaa1` |
| | Gpaa1_O43292 | - | Gpaa1 (w TMs) | | 0.797 | `super AF-O43292_Gpaa1, 7w72` |
| | Gpaa1_O43292 | - | - | Gpaa1 (w TMs) | 1.224 | `super AF-O43292_Gpaa1, 7wld` |
| | | Gaa1_P39012 (w TMs) | Gpaa1 (w TMs) | | 2.212 | `super bk_gaa1, 7w72 and chain A` |
| | | Gaa1_P39012 (w TMs) | - | Gpaa1 (w TMs) | 2.193 | `super bk_gaa1, 7wld` |
| | | - | Gpaa1 (w TMs) | Gpaa1 (w TMs) | 1.310 | `super 7w72 and chain A, 7wld and chain G` |

| | | | | | | |
|---|---|---|---|---|---|---|
| GPAA1-Zn | Gpaa1_O43292 (lumen) | - | - | - | 3.029 | super GPAA1-Zn, AF-O43292_Gpaa1 |
| GPAA1-Zn | - | Gaa1_P39012 (lumen) | - | - | 4.035 | super GPAA1-Zn, bk_gaa1 |
| GPAA1-Zn | - | - | Gpaa1 (lumen) | - | 3.020 | super GPAA1-Zn, 7w72 and chain A |
| GPAA1-Zn | - | - | - | Gpaa1 (lumen) | 3.254 | super GPAA1-Zn, 7wld and chain G |
| GPAA1-ZnZn | Gpaa1_O43292 (lumen) | - | - | | 2.763 | super GPAA1-ZnZn, AF-O43292_Gpaa1 |
| GPAA1-ZnZn | - | Gaa1_P39012 (lumen) | - | | 3.302 | super GPAA1-ZnZn, bk_gaa1 |
| GPAA1-ZnZn | - | - | Gpaa1 (lumen##) | | 5.375** | cealign 7w72 and chain A and resi 46-368 and ss h+b, GPAA1-ZnZn and ss h+b |
| GPAA1-ZnZn | - | - | - | Gpaa1 (lumen##) | 5.973** | cealign 7wld and chain G and resi 46-368 and ss h+b, GPAA1-ZnZn and ss h+b |
| | Gab1_P41733 | Gab1_P41733 | - | - | 0.295 | super AF-P41733-Gab1, bk_gab1 |

| | | | | | |
|---|---|---|---|---|---|
| Gab1_P41733 | - | Gab1_PIG-U | - | 1.741 | super AF-P41733-Gab1, 7w72 |
| Gab1_P41733 | - | - | PIG_U | 1.280 | super AF-P41733-Gab1, 7wld |
| - | Gab1_P41733 | Gab1_PIG-U | - | 1.959 | super bk_gab1, 7w72 |
| - | Gab1_P41733 | - | PIG_U | 1.290 | super bk_gab1, 7wld |
| - | - | Gab1_PIG-U | PIG_U | 0.997 | super 7w72 and chain U, 7wld and chain U |
| Gpi8_Q92643 | Gpi8_P49018 | - | - | 0.444 | super AF-Q92643-Gpi8, bk_gpi8 |
| Gpi8_Q92643 | - | Gpi8_PIG-K | - | 0.733 | super AF-Q92643-Gpi8, 7w72 |
| Gpi8_Q92643 | - | - | PIG_K | 0.899 | super AF-Q92643-Gpi8, 7wld |
| - | Gpi8_P49018 | Gpi8_PIG-K | | 0.852 | super bk_gpi8, 7w72 |
| - | Gpi8_P49018 | - | PIG_K | 0.951 | super bk_gpi8, 7wld |
| - | - | Gpi8_PIG-K | PIG_K | 0.932 | super 7w72 and chain K, 7wld and chain K |
| Gpi16_Q969N2 | Gpi16_P38875 | - | - | 1.177 | super AF-Q969N2-Gpi16, bk_gpi16 |
| Gpi16_Q969N2 | - | Gpi16_PIG-T | - | 0.709 | super AF-Q969N2-Gpi16, 7w72 |

| | | | | | |
|---|---|---|---|---|---|
| Gpi16_Q969N2 | - | - | PIG_T | 1.070 | super AF-Q969N2-Gpi16, 7wld |
| - | Gpi16_P38875 | Gpi16_PIG-T | - | 1.406 | super bk_gpi16, 7w72 |
| - | Gpi16_P38875 | - | PIG_T | 1.368 | super bk_gpi16, 7wld |
| - | - | Gpi16_PIG-T | PIG_T | 1.175 | super 7w72 and chain T, 7wld and chain T |
| Gpi17_Q04080 | Gpi17_Q04080 | - | - | 0.813 | super AF-Q04080-Gpi17, bk_gpi17 |
| Gpi17_Q04080 | - | Gpi17_PIG-S | - | 3.535 | super AF-Q04080-Gpi17, 7w72 |
| Gpi17_Q04080 | - | - | PIG_S | 3.404 | super AF-Q04080-Gpi17, 7wld |
| - | Gpi17_Q04080 | Gpi17_PIG-S | - | 2.67 | super bk_gpi17, 7w72 |
| - | Gpi17_Q04080 | - | PIG_S | 2.937 | super bk_gpi17, 7wld |
| - | - | Gpi17_PIG-S | PIG_S | 1.365 | super 7w72 and chain S, 7wld and chain S |

*To obtain best alignment, two alignment commands were used, i.e. *cealign* is used in sequence-independent manner, whereas *super* is used when decent structural similarity is known, e.g. comparing domain models between our models, AlphaFold, Baker's, PDB:7w72, and PDB:7wLD

***super* resulted in only a few fitting residues, so *cealign* was applied instead.

#Baker's models were retrieved from https://www.modelarchive.org/doi/10.5452/ma-bak-cepc

##Lumen (residues 46-368)
